# Contact-phagy remodels ER-mitochondria contact sites

**DOI:** 10.64898/2026.11.10.765636

**Authors:** Adriana Covarrubias-Pinto, Alexis González, Natalia Jimenez-Moreno, Carla Salomo-Coll, Sergio Alejandro Poveda-Cuevas, Rubina Kazi, Ashwin Balakrishnan, Pratima Verma, Javier Lizarrondo, Marisa Di Monaco, Borna Markusic, Isabeli Y.A. Osama, Ah Jung Heo, Matthias Koch, Yangxue Fu, Uxia Gestal-Mato, Jana Hoffman, Rajeshwari Rathore, Marion Basoglu, Elena Polishchuk, Thorsten Mosler, Stefan Eimer, Gerhard Hummer, Ana J. García-Sáez, Mike Heilemann, Florian Wilfling, Ramachandra M. Bhaskara, Simon Wilkinson, Ivan Dikic

## Abstract

Membrane contact sites between the endoplasmic reticulum (ER) and other organelles coordinate cellular organization, metabolism and stress responses, yet how these interfaces are selectively remodeled and removed remains poorly understood. Here we identify a selective autophagy pathway that targets defined ER-mitochondria contact-site (ERMCS) subdomains for turnover, a process we term Contact-phagy. The ER-resident chaperone SIGMAR1 forms a functional complex with the ER-phagy receptor CCPG1, and SIGMAR1– CCPG1 nanoclusters define ERMCS-associated sites of autophagy initiation. Contact-phagy is selectively engaged by metabolic and mitochondrial stress and promotes turnover of defined contact-site components without broadly affecting general ER-phagy, mitophagy or bulk autophagy. At these sites, SIGMAR1–CCPG1 function is coupled to mitochondrial fission: MFF promotes DRP1 recruitment to ERMCS, whereas loss of CCPG1 or SIGMAR1 causes abnormal persistence and clustering of DRP1 and MFF, accompanied by altered mitochondrial remodeling. Loss of CCPG1 further disrupts ER-mitochondria organization and mitochondrial function in pancreatic tissue, while pancreatic cancer cells depend on CCPG1 and SIGMAR1 for organelle integrity and bioenergetic capacity. Together, our findings establish Contact-phagy as a selective pathway for the turnover of defined organelle contact site subdomains and suggest a broader principle by which interorganelle interfaces are maintained.

## Main

The endoplasmic reticulum (ER) forms a dynamic network of sheets and tubules that establishes diverse membrane contact sites (MCSs) with other organelles to coordinate calcium exchange, lipid metabolism, and organelle dynamics^1^. Among these interfaces, ER– mitochondria contact sites (ERMCSs) function as central hubs for metabolic communication, autophagy initiation, and mitochondrial fission^2–4^. ERMCSs host key components of the autophagy initiation machinery ^5–7^. For example, the ER-tethers VAPA/B recruit ULK1 and FIP200 to nascent phagophores, promoting autophagosome formation^8^.

The ER-resident chaperone SIGMAR1 has emerged as an important regulator of ER-mitochondria signaling, particularly in stabilizing Ca^2+^ transfer^9,10^, and mutations in SIGMAR1 are linked to neurodegeneration marked by ER and mitochondrial dysfunction^10,11^. Yet, the molecular basis linking SIGMAR1 function at ERMCSs to autophagy remains largely unresolved.

In parallel, selective autophagy of the ER (ER-phagy) has become recognized as a key mechanism for maintaining ER homeostasis ^12–15^. The ER-phagy receptor CCPG1, which targets stressed or damaged ER regions for lysosomal degradation, plays an essential role in ER quality control^16,17^. Recently, it has been shown that ER contact sites can trigger ER-phagy at liquid-like condensates^18^. A major unresolved question is whether ER-phagy participates in tuning the structure or turnover of ERMCSs and how such mechanisms might influence organelle behavior and cellular metabolic homeostasis.

### SIGMAR1 forms a stable complex with CCPG1

To gain insights into the regulation of ERMCS and a potential role for autophagy, we focused on the ER chaperone SIGMAR1, which localizes to ERMCSs and frequently appears in interactome analyses of ER-phagy receptors, alongside other components of the mitochondrial membrane proteome ^19–21^. SIGMAR1 molecules form homo-trimeric structures important for ligand binding ^22^ and their function within a calcium-sensitive module also containing BiP/Grp78/HSPA5 (Glucose-Regulated Protein 78) and IP3R3^9^. To define the SIGMAR1 interactome, we generated an inducible cell line expressing HA-tagged SIGMAR1 and performed co-immunoprecipitation followed by mass spectrometry analysis **(Fig. 1a)**. This analysis identified 173 interacting proteins, including known partners such as BiP/Grp78, as well as previously unrecognized candidates, notably the ER-phagy receptor CCPG1, which binds FIP200 and LC3B **(Fig. 1b)**. The interaction between endogenous CCPG1 and SIGMAR1 was confirmed by immunoprecipitation and colocalization analysis **(Extended Data Fig. 1a-b)**. Non-denaturing blue native PAGE of immunoprecipitated CCPG1 further demonstrated that both proteins reside within the same complex **(Extended Data Fig. 1c)**. Consistently, both HA–SIGMAR1 and GFP–CCPG1 proteins colocalized along the ER network in tubular and punctate structures **(Extended Data Fig. 1d, arrowheads)**.

**Figure 1.**
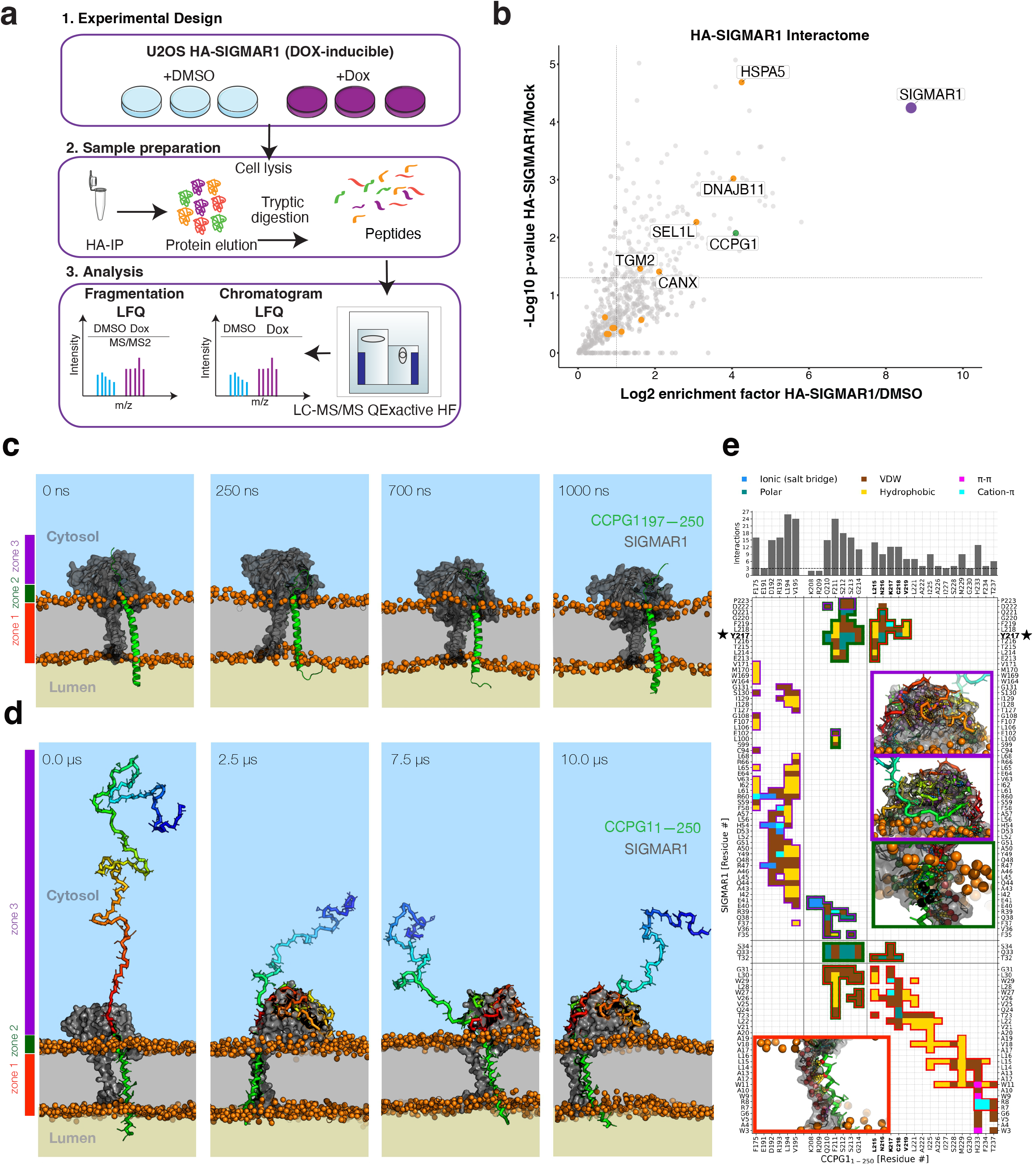
SIGMAR1 forms a complex with the ER-phagy receptor CCPG1. **a)** MS workflow for the analysis of HA-SIGMAR1 co-immunoprecipitated from cell lysates following the treatment doxycycline (Dox) (basal, purple) and DMSO (HA-empty, light blue). **b)** Volcano plot showing proteins enriched in HA-SIGMAR1 IP-MS. Data were analyzed using MaxQuant and Perseus (n = 3). Significant interaction partners were defined by log_2_ enrichment > 1.0 and −log_10_ *P* > 1.3 (one-sided unpaired Student’s *t*-test). The bait protein SIGMAR1 is highlighted in purple, CCPG1, the most significant interaction partner after BiP, is highlighted in green, and all other significant interaction partners are shown as orange dots. **c)** Representative snapshots from atomistic MD simulations showing interactions between the CCPG1_197-250_ (green ribbon) and SIGMAR1 (grey surface) in the POPC bilayer (phosphate groups, orange spheres). The transmembrane (zone 1, red), interfacial (zone 2, green), and cytosolic (zone 3, purple) regions of CCPG1 mediate contacts with SIGMAR1. **d)** Representative snapshots from coarse-grained MD simulations showing interactions between the extended CCPG1_1-250_ containing cytosolic IDR and SIGMAR1 (grey surface) within a POPC bilayer (phosphate groups shown as orange spheres), where the CCPG1-IDR (rainbow-colored sticks) scaffolds the cupid-domain of SIGMAR1 (grey). **e)** Heatmap of the pairwise residue contact map, obtained by averaging intermolecular contacts from coarse-grained MD trajectories. The plot highlights residue interactions (colored by interaction type) with high probability (*P*_contact_ > 0.5; also see **Extended Data Fig. e**). The number of interactions for key interface residues of CCPG1 is plotted above. Insets show zoomed-in views of stable contacts within the three interaction zones. CCPG1 residues mutated to alanine are highlighted in bold on the y-axis label. Y217 of SIGMAR1, which forms a central hub mediating multiple contacts, is marked with a star.

To elucidate the molecular basis of this interaction, we modeled the three-dimensional structure of the binary SIGMAR1–CCPG1 complex. SIGMAR1 contains a single transmembrane (TM) helix and a folded cupid-like domain associated with the cytosolic leaflet of the ER membrane ^22^. The topology of CCPG1 indicates that it contains a single TM helix, flanked by a cytosolic N-terminal intrinsically disordered region (IDR) housing the LC3B- and FIP200-interacting regions (LIR and FIR motifs) ^16^ and luminal C-terminal coiled-coil cargo-interacting regions (CIRs) ^23^. We therefore hypothesized that the TM helix and the N-terminal IDR of CCPG1 could potentially mediate direct physical interactions with SIGMAR1. AlphaFold2 modeling of full-length SIGMAR1 in complex with a CCPG1 fragment encompassing residues 197–250 revealed direct TM helix-helix interactions, with high confidence in the interacting CCPG1 region (residues 212-250; pLDDT ≥ 70) **(Fig. 1c)**.

To test the stability of this interaction in a membrane environment, the top-ranked SIGMAR1-CCPG1_197-250_ complex model was embedded in two different membrane systems: one a homogeneous PC-rich bilayer (POPC lipid-only) and the other that closely mimics ER composition, and both were subjected to all-atom molecular dynamics (MD) simulations (up to 1 µs; *n* = 2 replicates). The binary membrane-embedded complex remained highly stable throughout both simulations exhibiting minimal conformational changes **(Fig. 1c)**. Extension of the model to include the N-terminal IDR (residues 1-250) of CCPG1 followed by extensive coarse-grained MD simulations (up to 10 µs; *n* = 3 replicates) revealed additional contacts between the IDR and the cupid-like domain of SIGMAR1, further stabilizing the complex **(Fig. 1d)**. Mapping of pair-wise residue interactions identified three major interaction zones: (i) hydrophobic contacts within the membrane core, (ii) interactions at the membrane-water interface, and (iii) cytosolic contacts involving the CCPG1 IDR **(Fig. 1e)**. Further, these contacts were consistent in both atomistic and coarse-grained MD simulations **(Fig. 1e and Extended Data Fig. 1e-g)**. A detailed interaction analysis, including a breakdown of the physicochemical properties of these stabilizing contacts across all three zones, reveals primarily extensive hydrophobic contacts **(Supplementary Tables 1-2)** and a few specific polar interactions, including salt bridges, charged/polar contacts, and cation-π interactions. Cation-π contacts represented the dominant class of polar interaction, with SIGMAR1 Y217 emerging as a central interaction hub **(Fig. 1e)**. Hydrophobic and polar interactions within zones 1 **(red in Fig. 1d, e)** and 2 **(green in Fig. 1d, e)** contributed most strongly to complex stability, with a prominent interaction network spanning CCPG1 residues 215-219 **(Fig. 1e and Extended Data Fig. 1e-g)**. Consistent with these structural predictions, deletion of the CCPG1 TM helix (CCPG1 _ΔTM_) abolished binding with SIGMAR1 **(Extended Data Fig. 1h-i)**, whereas alanine substitution of L215, K217, and C218 (TM mutant), or of all five residues L215, N216, K217, C218, and V219 (CCPG1^5A^ mutant), only partially reduced the interaction **(Extended Data Fig. 1j)**. Together, these findings demonstrate that SIGMAR1 and CCPG1 form a stable transmembrane complex within the ER membrane.

### SIGMAR1 stabilizes CCPG1 clusters to drive efficient ER-phagy

We next examined whether SIGMAR1 regulates CCPG1-mediated ER-phagy. Upon nutrient starvation (EBSS) combined with bafilomycin A1 (BafA1) to block lysosomal degradation, mCherry–SIGMAR1 and GFP–CCPG1 accumulated within lysosomes, indicating active ER-phagy flux **(Fig. 2a and 2b, line inset)**. Degradation of both CCPG1 and SIGMAR1 during EBSS/CHX treatment was FIP200-dependent **(Extended Data Fig. 2a)**. SIGMAR1 showed a modest but reproducible decrease during prolonged EBSS/CHX treatment in control cells, whereas this loss was attenuated upon FIP200 depletion, resulting in significantly higher SIGMAR1 levels in FIP200-depleted cells at 6 and 8 h **(Extended Data Fig. 2b)**. Consistently, doxycycline-inducible HA–SIGMAR1 co-localized with the autophagosome and lysosome markers LC3B and LAMP1 under these conditions **(Extended Data Fig. 2c)**, indicating that SIGMAR1 itself is targeted to autophagic compartments.

**Figure 2.**
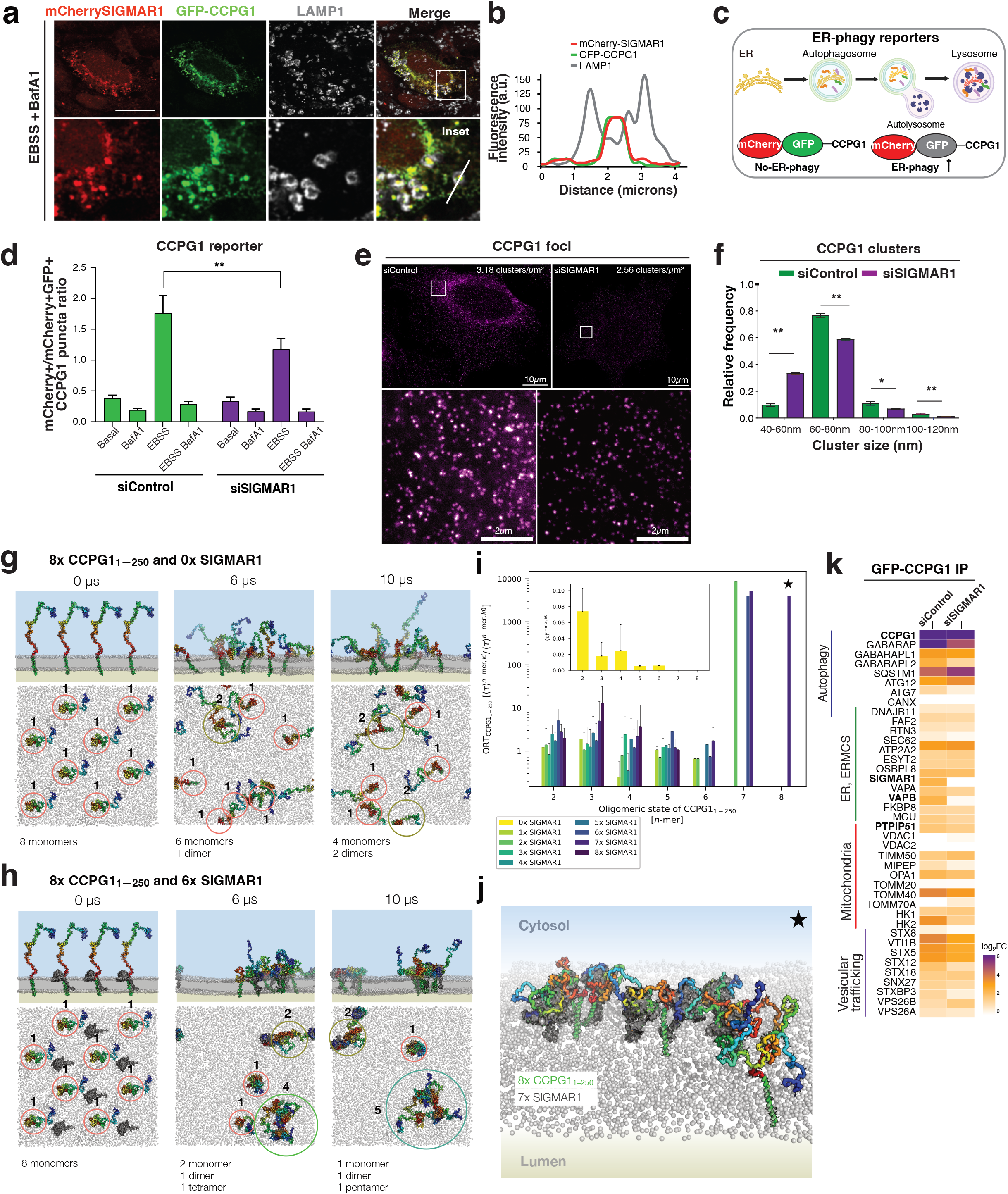
SIGMAR1 stabilizes the CCPG1 clusters and interactions for productive ER-phagy. **a)** Confocal fluorescence microscopy analysis of cells co-expressing mCherry-SIGMAR1 and GFP-CCPG1 and treated with EBSS and BafA1 (6 hours). Fixed cells expressing mCherry-SIGMAR1 (red) and GFP-CCPG1 (green) were stained with LAMP1 (grey). The line indicates co-localization of mCherry and GFP signals colocalized within the LAMP1^+^ vesicles. Scale bar, 10 µm. **b)** Histogram analysis of the fluorescence intensity distribution reveals that mCherry-SIGMAR1 and GFP-CCPG1 colocalizes within the lysosomes. **c)** ER-phagy reporter system: mCherry-GFP-CCPG1. **d)** ER-phagy flux was quantified as the ratio between mCherry^+^GFP^−^ and mCherry^+^GFP^+^puncta. n=3 independent experiments. The CCPG1 reporter was analyzed by high-throughput microscopy analysis, with the number of cells per condition: siControl (basal: 745; EBSS:669) and siSIGMAR1 (basal:379; EBSS:275), respectively. Data are mean ± s.d.; one-way ANOVA, Bonferroni post-hoc test. **e)** Representative STED images of HeLa cells immunostained for CCPG1 and their zoom-ins (bottom) in siControl and siSIGMAR1 conditions. **f)** CCPG1 cluster size distribution of siControl (green) and siSIGMAR1 (magenta) cells. The cluster sizes (diameter) are binned with a width of 20 nm, ranging from 40– 120 nm. Scale bar, 10 µm. **g)** Representative snapshots (side and top views) of coarse-grained MD simulations assessing CCPG1_1-250_ (green/rainbow spheres) oligomerization in the absence of SIGMAR1, and **h)** presence of SIGMAR1 (6x). 8 CCPG1_1-250_ molecules are highlighted (red circles) along with their nascent higher-order oligomers (4- and 5-mers). **i)** The distribution of the ratios between the average oligomeric residence times (ORTs) per CCPG1_1-250_ *n*-mer (see Methods) in the presence of a defined number of SIGMAR1 molecules is extracted from 9 different simulations (*n* = 3). Increasing the number of SIGMAR1 facilitates the formation of higher-order protein clusters with more CCPG1_1-250_. **j)** Representative snapshot of a high-order protein cluster in the ER containing eight CCPG1_1-250_ molecules co-clustered with seven SIGMAR1 molecules. SIGMAR1 cupin domains mediate bridging interactions with CCPG1-IDRs, increasing the lifetime to form stable TM–TM contacts. **k)** Heatmap showing the CCPG1 interaction profile in siControl and siSIGMAR1 knockdown cells. The relative abundance of proteins associated with CCPG1 was compared between the two conditions, including autophagy adaptors, vesicular trafficking proteins, ER and ER–mitochondria contact site (ERMCS) components, and mitochondrial proteins. Data were analyzed using MaxQuant and Perseus (n = 3).

To assess the functional contribution of SIGMAR1 to ER-phagy, we measured ER-phagy flux using a general ER reporter (ss-mRFP–GFP–KDEL ^24^, hereafter as KDEL) and a CCPG1-specific tandem reporter (mCherry–GFP–CCPG1, hereafter as CCGP1) **(Fig. 2c)**. High-content imaging revealed a significant reduction in starvation-induced ER-phagy upon siRNA-mediated knockdown (KD) of either SIGMAR1 or CCPG1 **(Extended Data Fig. 2e-g)**. Notably, SIGMAR1 depletion reduced lysosomal delivery of both CCPG1 and KDEL in starved cells **(Fig. 2d and Extended Data Fig. 2f)**, suggesting that SIGMAR1 facilitates CCPG1-driven ER-phagy.

CCPG1 forms discrete ER-associated foci that recruit FIP200, WIPI2 and LC3B upon autophagy induction^16^, consistent with receptor oligomerization. Co-immunoprecipitation experiments showed that CCPG1 self-associates, with this interaction requiring an intact transmembrane domain and partially depending on the N-terminal region **(Extended Data Fig. 2h; heterodimerization between CCPG1**–**CTD mutant and CCPG1**–**WT)**. Given the chaperone function of SIGMAR1 at ERMCS^9^, we tested whether it stabilizes CCPG1 oligomers. Blue native PAGE revealed high-molecular-weight CCPG1 complexes **(Extended Data Fig. 2i)**, which were markedly reduced upon SIGMAR1 KD, accompanied by accumulation of lower-order oligomers **(Extended Data Fig. 2i)**.

Super-resolution STED microscopy further demonstrated that SIGMAR1 depletion significantly reduced CCPG1 cluster size and density **(Fig. 2e-2f and Extended Data Fig. 2j),** indicating impaired cluster stabilization. To provide mechanistic insights into SIGMAR1-mediated CCPG1 clustering, we performed coarse-grained MD simulations of CCPG1 oligomerization in large membrane patches containing increasing numbers of SIGMAR1 molecules (0 to 8 molecules; up to 10 µs; *n* = 3 replicates; **Fig. 2g-h**). By tracking individual CCPG1 molecules and their oligomeric state across all simulation trajectories, we characterized the extent of CCPG1 clustering as a function of SIGMAR1 concentration. In the absence of SIGMAR1, we found that CCPG1 predominantly organized readily into dimers **(Fig. 2g)** and occasionally into trimers and tetramers, and only rarely formed higher-order pentamers and hexamers as indicated by their average residence times (⟨*τ*⟩*^n-mer,k^*^0^ distribution; see inset in **Fig. 2i-2j**). This indicated that CCPG1 has an intrinsic capacity for low-order self-association in PC-rich bilayers. By contrast, the presence of SIGMAR1 molecules within the membrane patches led to the formation of higher-order CCPG1 clusters **(Fig. 2h)**. In these simulations, we found that a heteromeric binary complex of CCPG1–SIGMAR1 often formed first and nucleated the aggregation of higher-order CCPG1 clusters **(Fig. 2i)**. The growth of the largest CCPG1 clusters was often correlated with the presence of SIGMAR1 within the cluster **(Extended Data Fig. 2k)**. Analysis of quaternary interactions in the membrane further indicates that SIGMAR1 increases the persistence of CCPG1–CCPG1 contacts and can act as a membrane-embedded scaffold that bridges CCPG1 molecules into larger assemblies **(Extended Data Fig. 2k)**.

Next, by measuring the ratio of the average residence times of CCPG1 *n*-mers (2–8 mers) in the absence and presence of a fixed number of SIGMAR1 molecules (*ki* = [0, …, 8]), we quantified the effect of SIGMAR1. We found that higher-order CCPG1 oligomers were reproducibly enriched only in the presence of multiple SIGMAR1 molecules (ratio > 1.0; ratio = 1 indicates background CCPG1 self-association into *n*-mers). In particular, with ≥4 SIGMAR1 molecules, CCPG1 oligomerization tendency increased by 10–20-fold to form 3–6-mers and ≥ 1000-fold to form CCPG1-heptamers and -octamers **(Fig. 2i)**. These findings indicate that SIGMAR1 promotes and stabilizes higher-order CCPG1 assemblies, rather than being strictly required for initial CCPG1 oligomerization. This was consistent with STED microscopy and native PAGE analyses, which showed reduced CCPG1 cluster size/density and decreased high-molecular-weight CCPG1 assemblies upon SIGMAR1 depletion **(Fig. 2e, 2f, 2j and Extended Data Fig. 2j)**. These simulations support a model in which the TM and IDR segments of CCPG1 contribute to homo- and hetero-oligomerization, while SIGMAR1 shifts the assembly equilibrium toward larger CCPG1 clusters.

Finally, proteomic analysis of GFP-CCPG1 immunoprecipitates revealed that SIGMAR1 depletion altered the CCPG1 interactome **(Fig. 2k and Extended Data Fig. 2l-m)**. Loss of SIGMAR1 reduced CCPG1 association with autophagy machineries (GABARAPs, ATG7-12, and p62), ER/ERMCS components (RTN3, OSBPL8, VAPA, VAPB), Mitochondria/ERMCS constituents (MCU, PTPIP51/RMD3, TIMM50, TOMM20, MIPEP, HK1-2) and vesicular trafficking proteins (STX8, VTI1B, STX5, STX12, SNX27 and STXBP3) **(Fig. 2k and Extended Data Fig. 2l-m)**. Pathway analysis revealed that loss of SIGMAR1 increased the association of CCPG1 with proteins involved in RNA processing, protein biosynthesis, and ER-associated protein targeting and localization, indicating a remodeling of the CCPG1 interactome in the absence of SIGMAR1 **(Extended Data Fig. 2n)**. Together, these data demonstrate that SIGMAR1 stabilizes CCPG1 clustering at the ER and promotes productive interactions with the autophagic machinery to drive efficient ER-phagy.

### SIGMAR1 and CCPG1 are localized at ERMCS

SIGMAR1 is an integral component of ERMCS ^10^ and contributes to CCPG1 interactions with proteins involved in ERMCS integrity and autophagy **(Fig. 2k)**. To define the molecular environment of SIGMAR1 at ERMCS, we enriched functional SIGMAR1 oligomers using biomolecular complementation affinity purification (BiCAP) ^25^. Full-length SIGMAR1 was fused to complementary, non-fluorescent Venus fragments (V1–and V2–SIGMAR1), such that homomeric complex formation restored the Venus fluorescence. The reconstituted signal localized to ER membranes marked by REEP5, confirming correct assembly and localization of split-Venus-tagged SIGMAR1 complexes **(Extended Data Fig. 3a-b)**. Next, we characterized the interactome of the SIGMAR1 dimers, revealing 475 significantly enriched proteins (log2 enrichment factor ≥ 1; p-value ≤ 0.05). CCPG1 was one of the most enriched interacting partners of SIGMAR1 dimers, confirming our previous results. In addition, other ER-phagy receptors (e.g., RTN3 or TEX264) **(Extended Data Fig. 3c, red dots)**, ERMCSs components such as CANX, PDIA6, and VAPB, and the mitochondrial outer membrane import machinery TOMM1, TOMM22, and TOMM34 **(Extended Data Fig. 3c, green dots),** as well as ER chaperones, ER structural components, and protein secretion machinery were identified in the SIGMAR1 interactome **(Extended Data Fig. 3c, blue dots)**. Altogether, these data provided evidence that oligomeric SIGMAR1 associates with CCPG1, ERMCS proteins, and outer mitochondrial proteins.

Immunofluorescence analysis demonstrated that punctate structures of CCPG1 and SIGMAR1 occurred in proximity to mitochondrial membranes positive for mitochondrial outer membrane component TOMM20^26^ **(Extended Data Fig. 3d, arrowheads)**. Using co-immunoprecipitation assays, the interaction between SIGMAR1, CCPG1 and the ERMCS protein IP3R3 was confirmed in our cellular model **(Extended Data Fig. 3e)**. Then, we performed subcellular fractionation to isolate ER, crude mitochondria, pure mitochondria, and MAM (mitochondria-associated membranes), as previously reported ^27^. SIGMAR1 and CCPG1 were detected in the ER and IP3R3-positive isolated MAM fraction, but were absent from the fraction containing pure mitochondria **(Extended Data Fig. 3f)**, supporting their localization at ERMCS.

### Contact-phagy promotes ERMCS turnover

We determined that CCPG1 and SIGMAR1 are enriched at ERMCS, thus ER-phagy driven by CCPG1 can initiate at ERMCS, raising the possibility that this pathway can influence the organization of these interfaces. To explore this idea, we performed quantitative mass spectrometry in order to reveal CCPG1-dependent proteome remodeling upon starvation **(Fig. 3a)**. Analysis of crude mitochondrial fractions comparing parental and CCPG1-knockout (KO) HeLa cells under basal and EBSS conditions showed increased abundance of ER-resident ERMCS components (ATL3, CALR, CANX, DNAJB11, FAF2, HSP90B1, BiP, NPC1, P4HB, RTN3, SEC62, PDIA4, STIM2, SEL1L, ATP2A2, ERMCS: ESYT1/2, RAB18, SIGMAR1, VAPA/B), mitochondrial proteins (FKBP8, MXU, VDAC1/2/3, HSPA9, TIMM50, TOMM20, TOMM40, HK1/2) and autophagy related proteins (GABARAP, PEX14, PEX19, SQTM1 and TGM2) **(Fig. 3b-c and Extended Data Fig. 3g)**. A similar and additional set of proteins from ER and mitochondria (FAM134A, RTN1 and ATP2A2 and TOMM70) **(Extended Data Fig. 3h-i**) were upregulated in the total proteome of CCPG1-KO cells, further suggesting that a subset of ER and mitochondrial proteins can be targeted for degradation via CCPG1-dependent ER-phagy.

**Figure 3.**
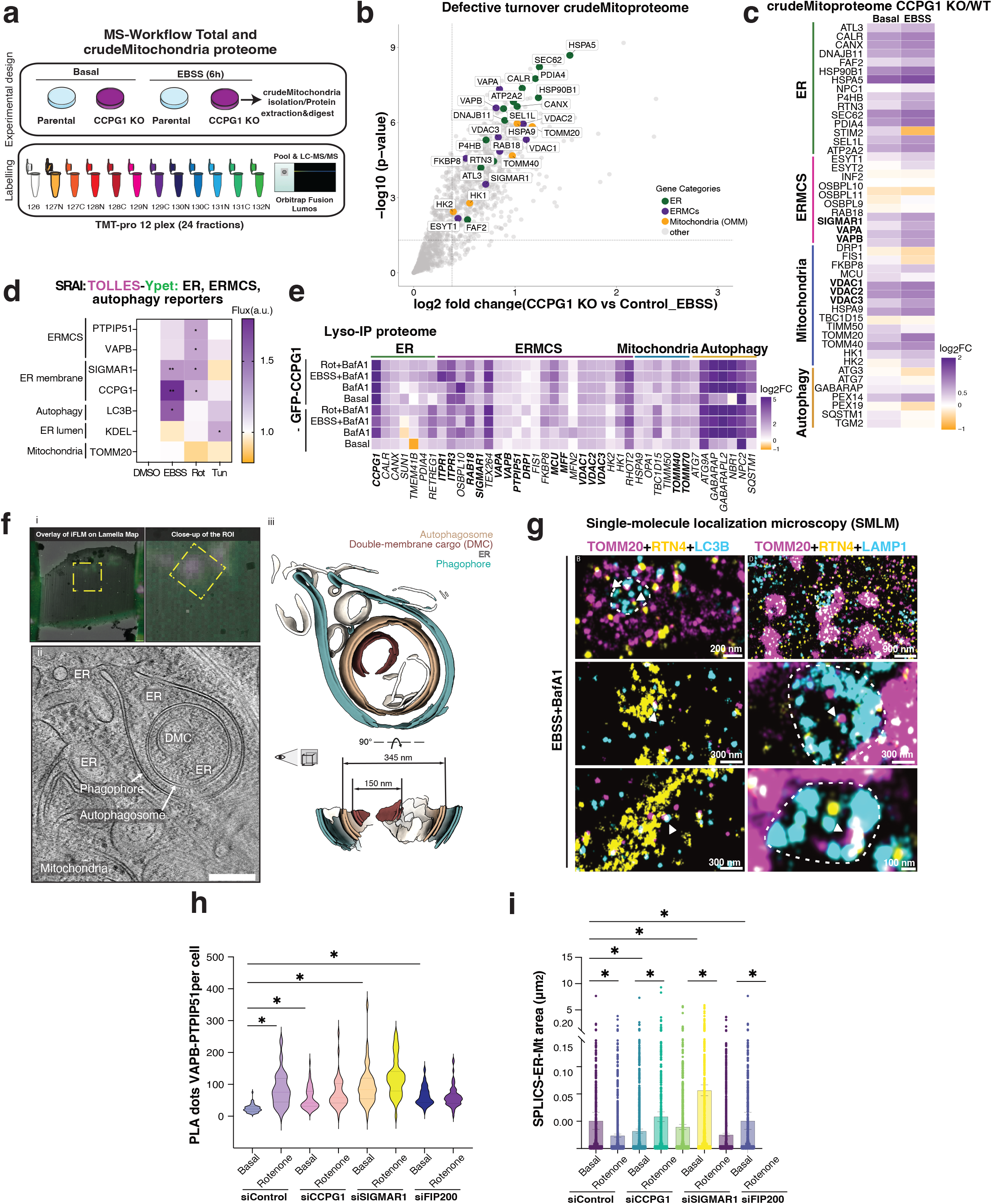
Starvation induces ERMCS degradation depending on the SIGMAR1-CCPG1 complex. **a)** TMT-labeled total cell proteome from WT and CCPG1 KO HeLa cells in Basal and 6h of EBSS, three replicates (3xR) per condition, MS workflow. **b)** Volcano plot showing ER, mitochondrial, and ER–mitochondria contact site (ERMCS) proteins with defective turnover in the crude mitochondrial proteome of CCPG1 KO cells under basal and EBSS conditions. Proteins with differential abundance in CCPG1 KO cells were analyzed using MaxQuant and Perseus (n = 3). Proteins with FC enrichment > 1.3 and −log_10_ *P* > 1.3 were considered significant (two-sided unpaired Student’s *t*-test). **c)** Heatmap showing log_2_ fold changes of ER, mitochondrial, ER–mitochondria contact site (ERMCS) proteins and autophagy proteins in the crude mitochondrial proteome of CCPG1 KO versus control cells under basal and EBSS conditions. Positive log_2_ fold changes are shown in purple, whereas negative log_2_ fold changes are shown in green (n = 3). **d)** Heatmap of fluorescence ratios (flux indices, see Methods) of HeLa cells stably-expressing SRAI fusions with the indicated proteins under DMSO, starvation (EBSS), rotenone (Rot, 1 μM) or tunicamycin (Tun, 10 μg/mL) treatments for 6 h (n = 3 independent replicates, ± s.e.m, 6 fields of 20-100 cells per replicate, normalised to DMSO, 1-sample two-tailed t-tests, **= p<0.01, * = p≤ 0.05). **e)** Heatmap showing log_2_ fold changes of ER, mitochondrial, ER–mitochondria contact site (ERMCS) proteins and autophagy proteins enriched in LysoIP under different conditions. Positive log_2_ fold changes are shown in purple, whereas negative log_2_ fold changes are shown in green (n = 3). **f)** Correlative cryo-ET of HCT116 cells under EBSS starvation and treated BafA1. **i**. Overlay of the lamella fluorescence from the integrated fluorescence microscope at the plasma FIB and the lamella map acquired at the TEM. A close up around the region of interest (ROI) is shown and the position of the tilt series acquisition is denoted by a yellow rectangle. **ii**. Slice of the tomogram reconstructed from the tilt series in the position indicated in **i** (rotated by 90° counter clockwise). The reconstruction shows a phagophore next to the ER engulfing another autophagosome with a mixed cargo, a double membrane cargo (DMC) and ER. **iii**. Segmentation of the membranes corresponding to the phagophore (blue), the ER (white), the autophagosome (red) with a double membrane vesicle (white) and ER-like membrane (orange) in the tomogram shown in **ii**. Scale Bar 200nm. **g)** Exchange-PAINT image of TOMM20+, RTN4+ and LC3B+ or LAMP1+ vesicles in HeLa cells treated with EBSS and Bafa1 (6 h). Arrowheads indicate TOMM20 and RTN4 signal in close proximity within LC3B and LAMP1 vesicles. **h)** Violin plot of PLA dots of VAPB-PTPIP51 coupling per cell in siControl, siCCPG1M, siSIGMAR1 and siFIP200 cells treated with Rot (1uM, 6h). Data are mean ± s.e.m. from three independent experiments Basal n=61 for siControl, n=63 for siCCPG1, n=51 for siSIGMAR1 and n=55 for siFIP200 and Rotenone n=51 for siControl, n=40 for siCCPG1, n=42 for siSIGMAR1 and n=59 for siFIP200; Kruskal–Wallis/Dunn’s post-test. **i)** SCPLICs-ER-Mt contacts circularity index in siControl, siCCPG1 and siSIGMAR1 cells treated with Rotenone. (H) and (I) Data are mean ± s.e.m. from three independent experiments Basal siControl (n=21), siCCPG1 (n=16), siSIGMAR1 (n=14) and siFIP200 (n=14) and Rotenone siControl (n=24), siCCPG1 (n=14), siSIGMAR1 (n=21) and siFIP200 (n=23); Kruskal– Wallis/Dunn’s post-test.

The increased abundance of ERMCS proteins in CCPG1-KO cells, together with the association of the CCPG1–SIGMAR1 complex with ERMCS components, suggested that these proteins may undergo selective autophagic turnover. Among them, VAPB and PTPIP51 (RMDN3) form a well-established ER-mitochondria tether spanning the two organelles. We therefore asked whether ERMCS components could undergo selective autophagic degradation-not correlated simply with increased general ER-phagy flux-by applying different cellular stressors that selectively induce ERMCS degradation. We tested mitochondrial stress (rotenone), ER stress (tunicamycin) and starvation (EBSS) using sensitive SRAI (Signal-Retaining Autophagy indicator) flux reporter^28^ for CCPG1, SIGMAR1, PTPIP51 and VAPB, alongside reporters for general ER-phagy (ss-SRAI–KDEL) and mitophagy (TOMM20). Starvation predominantly induced CCPG1 and SIGMAR1 turnover, whereas tunicamycin promoted general ER-phagy **(Fig. 3d and Extended Data Fig. 3j)**. By contrast, rotenone selectively induced co-degradation of CCPG1, SIGMAR1 and ERMCS proteins without inducing general ER-phagy or mitophagy **(Fig. 3d and Extended Data Fig. 3j)**. Consistent with their autophagic turnover, Lyso-IP proteomics revealed CCPG1-dependent lysosomal enrichment of ERMCS proteins, together with ER and mitochondrial components, which was further enhanced following starvation or rotenone treatment **(Fig. 3e and Extended Data Fig. 3k–l; highlighted in bold)**. Rotenone-induced autophagic turnover of ERMCS proteins was blocked by SIGMAR1 inhibition with haloperidol **(Extended Data Fig. 3m, n)** and by CCPG1 silencing **(Extended Data Fig. 3o)**, demonstrating that stress-induced selective ERMCS turnover requires the SIGMAR1–CCPG1 pathway.

We next asked whether this selective turnover involves co-sequestration of ER and mitochondrial material within the same autophagic structures. Ultrastructural analysis of cells treated with EBSS and bafilomycin A1 revealed autolysosomes containing both ER and mitochondrial material, accounting ~20% of autolysosomes in control cells **(Extended Data Fig. 3p, q)**. Depletion of CCPG1 markedly reduced the proportion of mitochondria-containing autolysosomes, whereas SIGMAR1 depletion almost completely abolished them **(Extended Data Fig. 3p, q),** supporting a role for the CCPG1-SIGMAR1 complex in the coordinated sequestration of ER and mitochondrial material.

To characterize ER-mitochondria structures within autophagic compartments, we employed HCT116 cells, which display robust vitrification and a starvation response comparable to HeLa cells, as assessed by SPLICS–ER–Mt (split-GFP ERMCS reporter which reconstitute fluorescence when the ER and mitochondrial membranes come into close proximity^29,30^) co-localization with LC3B **(Extended Data Fig. 3r)**. Correlative cryo-electron tomography (cryo-ET)^31^ targeted regions of CCPG1/LC3B or CCPG1/TOMM20 colocalization during lamella preparation **(Extended Data Fig.4a-d)**. Tomogram analysis revealed autophagic compartments containing ER-like membranes and the internal double-membrane vesicles **(Fig. 3f and Extended Data Fig. 5a)**, with outer vesicles measuring ~270-330 nm and the internal double-membrane vesicles ~100-200 nm. Owing to the resolution limits of the cryo-fluorescence correlation, the mitochondrial origin of internal vesicles could not be definitively assigned.

To complement cryo-ET, diffraction-limited and super-resolution microscopy approaches were used. Volumetric confocal imaging revealed colocalization of ER (RTN4) and mitochondrial (TOMM20) membranes within LC3B-positive autophagosomes and LAMP1-positive autolysosomes upon autophagy induction **(Extended Data Fig. 6a)**. This was further supported by STED microscopy and exchange-DNA based point accumulation for imaging in nanoscale topography (exchange-PAINT), which revealed ER (CCPG1 or RTN4) and mitochondrial (TOMM20) membranes in proximity within LAMP1- and LC3B-positive vesicles **(Fig. 3g**, **Extended Data Fig. 6b-c)**. Exchange-PAINT measurements showed ER and mitochondrial membranes within <60 nm of each other inside autophagic compartments **(Fig. 3g**, **Extended Data Fig. 6c)**, accounting for antibody linkage error^32^. Consistently, we performed high-resolution confocal imaging followed by LIGHTNING deconvolution using SPLICS–ER–Mt, ERMCS puncta frequently colocalized with LC3B, RTN4 and the mitochondrial marker TOMM20, identifying autophagy-associated structures at ER-mitochondria interfaces **(Extended Data Fig. 6d)**. Exchange-PAINT corroborates this further, where SPLICS–ER–Mt reporter localized within LC3B- and LAMP1-positive structures **(Extended Data Fig. 6e)**. Notably, cryo-ET analysis targeting SPLICS-positive sites revealed ER–mitochondria contact sites in which ER membranes were closely apposed to mitochondria specifically at sites of mitochondrial constriction, consistent with the proposed role in fission^33^ **(Extended Data Fig. 5b)**. Finally, *in situ* PLA detecting endogenous IP3R3-VDAC1 confirmed the presence of ERMCSs within lysosomes under EBSS and bafilomycin A1 treatment **(Extended Data Fig. 6f)**. Together, these data demonstrate that ERMCS can be selectively targeted for autophagic degradation.

To assess ERMCS remodeling and abundance during starvation and rotenone, we examined the coupling between VDAC1–IP3R3 and VAPB–PTPI51 using Proximity ligation assay (PLA). Cells lacking CCPG1, SIGMAR1, or the autophagy initiation factor FIP200 displayed a significant increase in IP3R3–VDAC1 PLA signals compared with control cells **(Fig. 3h and Extended Data Fig. 6g–h)**, indicating increased basal ERMCS abundance or stabilization of these contacts. A similar increase in PLA signal along the VAPB–PTPIP51 axis was observed in CCPG1- and SIGMAR1-deficient cells **(Extended Data Fig. 6i)**. In control cells, starvation or rotenone also increased ERMCS signals, consistent with stress-induced remodeling of ER– mitochondria contacts. In contrast, these stressors did not further increase PLA signals in CCPG1- or SIGMAR1-deficient cells, suggesting that loss of Contact-phagy impairs efficient ERMCS remodeling and resolution, resulting in persistence of these interfaces. To capture remodeling in ERMCS architecture (monitoring area changes), we employed the fluorescent contact site reporter SPLICS–ER–Mt. In siControl cells, either starvation or rotenone led to a marked redistribution of the SPLICS signal, which became smaller **(Extended Data Fig.6j-k)** characterized by a reduction in contact area **(Figure 3i**, **Extended Data Fig. 6j)**, consistent with active remodeling of ERMCS. In contrast, CCPG1- and SIGMAR1-deficient cells exhibited smaller SPLICS-positive area in basal conditions that enlarged upon rotenone or starvation, contrary to the siControl cells thus indicating impaired contact site remodeling **(Figure 3I**, **Extended Data Fig. 6k)**. Notably, pharmacological inhibition of SIGMAR1 (antagonist Haloperidol)^9,34^ prevented stress-induced PLA dots increase **(Extended Data Figure 6l-m)**. Consistent with this finding, SIGMAR1 inhibition also prevented stress-induced SPLICS remodeling **(Extended Data Figure 6n-o)**. Together, these findings suggest that ERMCS abundance and remodeling are regulated by an ER-phagy receptor-dependent process, which we term Contact-phagy.

### DRP1 associates with SIGMAR1-CCPG1 and contributes to Contact-phagy

Thus far, we have defined an ER-resident Contact-phagy complex. However, additional machinery is likely required to fragment ERMCS prior to their autophagic turnover. To identify such factors, we performed proteomic analysis of ER–mitochondria contact sites using a Venus-based VAPB–PTPIP51 complementation system. VAPB–PTPIP51 proved to be a robust substrate for this pathway and underwent CCPG1–SIGMAR1-dependent remodeling upon autophagic induction **(Fig. 3c, d, h)**.

Complementary Venus fragments fused to VAPB and PTPIP51 reconstituted at ER-mitochondria interfaces, enabling immunoprecipitation followed by mass spectrometry (MS) to identify proteins associated with these contact sites **(Fig. 4a and Extended Data Fig. 7a)**. MS analysis identified a broad range of ERMCS-associated proteins under both Basal as well EBSS+BafA1 conditions. Pathway enrichment analysis highlighted proteins involved in ER and mitochondria organization, ERMCSs biology, vesicle-mediated transport, intracellular protein transport, mitochondrial organization and fission, and autophagy **(Fig.4b and c)**. Notably, upon EBSS+BafA1 treatment, several interactions increased their association to VAPB–PTPIP51 contact sites. Among the enriched proteins were components of ERMCS and fission machineries (ESYT1, **SIGMAR1**, ITPR2, VDAC1/2, TOMM40, VPS13C, **DNM1L/DRP1**, FIS1, **MFF**) **(Fig.4d)**. Co-immunoprecipitation of CCPG1, SIGMAR1 with the Venus VAPB–PTPIP51 contact sites tether reporter further confirmed their association **(Extended Data Fig. 7b)**. Together, these data suggest that mitochondrial fission machinery is recruited to ER-mitochondria contact sites during autophagy induction and may cooperate with the CCPG1– SIGMAR1 complex to promote Contact-phagy.

**Figure 4.**
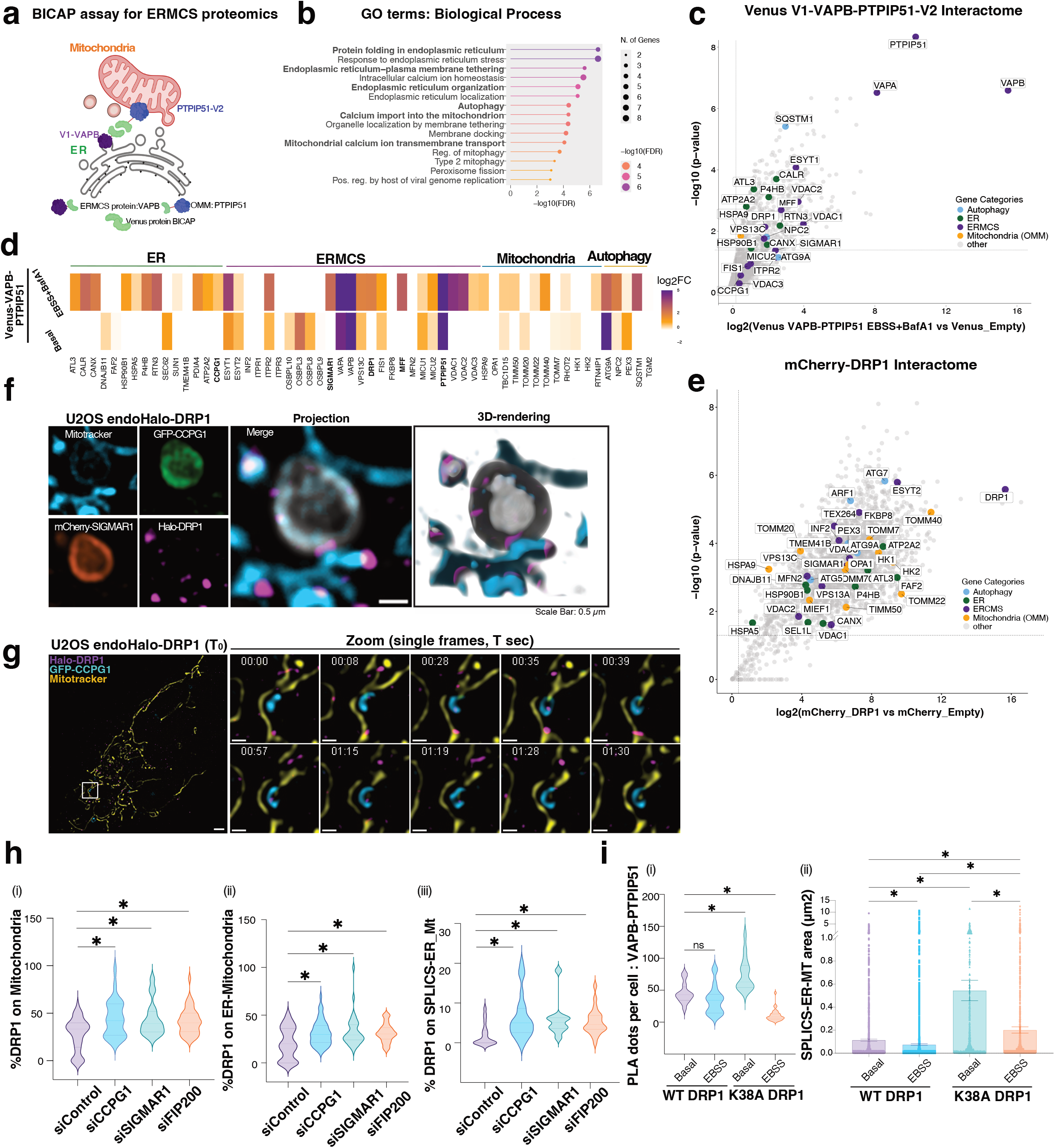
DRP1 associates to SIGMAR1-CCPG1 complex and contributes to Contact-phagy. **a)** Schematic showing the ER and mitochondrial proteins tagged with the TUBRO-ID for proximity proteomics. **b)** Pathway enrichment analysis from the top hits and common partners among all the baits, highlighting mitochondrial morphogenesis and outer mitochondrial membrane organization. **c)** Volcano plot showing proteins enriched in Venus– VAPB–PTPIP51 under EBSS conditions compared with Empty Venus. Data were analyzed using FragPipe (n = 3). Proteins with a fold-change > 1.3 and −log_10_ *P* > 1.3 were considered significantly enriched based on a two-sided moderated *t*-test. The baits VAPB and PTPIP51 (RMDN3), together with other ER–mitochondria contact site (ERMCS) proteins, are highlighted in purple. Significantly enriched ER, mitochondrial, and autophagy-related proteins are also highlighted. **d)** Heatmap showing log_2_ fold changes of ER, mitochondrial, ER–mitochondria contact site (ERMCS) proteins and autophagy proteins Venus–VAPB–PTPIP51 compared with Empty Venus under basal and EBSS conditions. Positive log_2_ fold changes are shown in purple, whereas negative log_2_ fold changes are shown in green (n = 3). **e)** Volcano plot showing proteins enriched in mCherry-DRP1 compared with mCherry-empty. Data were analyzed using MaxQuant and Perseus (n = 3). Proteins with a fold-change > 1.3 and −log_10_ *P* > 1.3 were considered significantly enriched based on one-sided unpaired Student’s *t*-test. The baits DRP1(DNM1L) together with other ER–mitochondria contact site (ERMCS) proteins, are highlighted in purple. Significantly enriched ER, mitochondrial, and autophagy-related proteins are also highlighted. **f)** SIM images showing association of DRP1 assemblies on mitochondria with SIGMAR1-CCPG1 complex in U2OS-endoHalo-DRP1 cells. Left panels show individual channels for mitochondria (cyan; labelled with BioTracker^TM^ 405 Blue mitochondria dye), GFP-CCPG1(green), mCherry-SIGMAR1 (orange), Halo-DRP1 (magenta, labelled with JF646). The middle panel shows the merged projection of all four channels, with mCherry-SIGMAR1 and GFP-CCPG1 merged in gray for simplified representation. The rightmost panel depicts the 3D rendering of the merged image shown in the middle panel. Scale Bar: 0.5 um. **g)** Snapshots from live-SIM imaging of U2OS-endoHalo-DRP1 cell (left panel) mitochondria (yellow; labelled with MitoTracker Orange CMTMRos), GFP-CCPG1 (cyan) and Halo-DRP1 (magenta; labelled with JF646) at time=0 (T_0_), Scale Bar, 2 um. Right panel: Time-lapse images of the boxed region in the left panel showing DRP1 interaction with CCPG1. Scale Bar, 0.5 um. Frames were acquired every 2.21 seconds for ~2 minutes. **h)** Violin plots showing the percentage of DRP1 associated with mitochondria (i), with ER (ii), with SPLICS-ER-Mt for ERMCs, from the quantified SIM images of U2OS-endomGFP-DRP1 and endoHalo-DRP1 cells respectively, with CCPG1, SIGMAR1, and FIP200 knockdown and control cells. (For (i) mitochondria and (ii) ER, n=19, n= 14, n= 24 and n=20, respectively; For (iii) SPLICS-ER-Mt, (n=28, n=15, n=29 and n=28) from 2 independent sets of experiments), Kruskal–Wallis/Dunn’s post-test; *p<0.05. **i)** PLA dots analysis **(i)** VAPB-PTPI51 in WT DRP1 expressing cells (basal n=25 and EBSS n=31) and K38A DRP1 expressing cells (basal n=36 and EBSS n=29). Data represent the mean ± s.e.m. from three independent experiments; Kruskal–Wallis/Dunn’s post-test. **(ii)** SPLICs-ER-MT remodeling by monitoring the area changes upon starvation in WT DRP1 expressing cells (basal n=21 and EBSS n=20) and K38A DRP1 expressing cells (basal n=14 and EBSS n=27). Data represent the mean ± s.e.m. from three independent experiments; Kruskal–Wallis/Dunn’s post-test.

To further investigate this relationship, we analyzed the DRP1 interactome by MS. SIGMAR1 emerged as a prominent DRP1 interactor, together with several outer mitochondrial membrane proteins and ERMCS-associated factors **(Fig. 4e)**. These interactions were validated biochemically, as DRP1 co-immunoprecipitated with both SIGMAR1 and CCPG1 **(Extended Data Fig. 7c),** in a manner dependent on the SIGMAR1 chaperone domain **(Extended Data Fig. 7d)**. Consistently, multi-color live cell imaging using high-resolution structured illumination microscopy (SIM) revealed the association of SIGMAR1 and CCPG1 with mitochondria-positive structures **(Fig. 4f and Extended Data Fig. 7e)**. DRP1 localized in close proximity to these ER-mitochondria interfaces evidencing the recruitment of mitochondrial fission machinery to Contact-phagy sites **(Fig.4f, 3d rendering and Extended Data Fig.7e, insets)**. The organization of these structures is consistent with the sequestration of ER-mitochondria contact-site components, as further illustrated in **Fig 4g**. Time-lapse imaging showed GFP– CCPG1-positive membrane progressively wrapping around a mitochondrial fragment containing DRP1, supporting the engulfment of ER-mitochondria contact-site material into autophagosomal compartments for subsequent degradation. Importantly, we could detect DRP1 accumulation in constriction sites at these ER-mitochondria contact interfaces, where it likely contributes to the final scission of the sequestered mitochondrial and ER fragments.

To further characterize these associations, we performed confocal imaging followed by LIGHTNING deconvolution. The ERMCS reporter Venus VAPB–PTPIP51 structures frequently colocalized with LC3B and DRP1 **(Extended Data Fig. 7f)**. Multi-target STED microscopy of DRP1, LAMP1 and SPLICS-ER-Mt further corroborated this, where LAMP1 positive vesicles could be found containing signal from both DRP1 and SPLICS–ER–Mt **(Extended Data Fig. 7g)**. Notably, DRP1 itself underwent degradation upon starvation in a FIP200-dependent manner **(Extended Data Fig. 7h-i)**, suggesting that DRP1 can be co-degraded with the SIGMAR1–CCPG1 complex during Contact-phagy. Consistent with this, CCPG1-deficient cells showed stabilization of DRP1, SIGMAR1 and PTPIP51 under starvation conditions, with a more modest stabilization of VAPB **(Extended Data Fig. 7j-m)**.

We next examined the subcellular distribution of DRP1 in Contact-phagy-deficient cells using SIM imaging. DRP1 showed increased association with both ER, marked by Halo–VAPB, and mitochondria, labeled with Mitotracker Orange **(Fig. 4h**; **Extended Data Fig. 7n-o)**. Consistently, DRP1 localization at ERMCS, visualized using the SPLICS reporter, was increased in CCPG1-, SIGMAR1-and FIP200-deficient cells **(Fig. 4h and Extended Data Fig. 7p-q)**. This redistribution was accompanied by altered DRP1 nano-assemblies. STED microscopy revealed reduced DRP1 cluster size in siCCPG1 cells and, more prominently, in siSIGMAR1 cells, whereas cluster density was significantly increased upon SIGMAR1 depletion and showed a trend toward an increase following CCPG1 or FIP200 depletion **(Extended Data Fig. 8a–d)**. Thus, defective Contact-phagy alters both the abundance and spatial organization of DRP1 at ER-mitochondria interfaces.

We next asked whether DRP1 recruitment to ERMCS depended on MFF, an outer mitochondrial membrane receptor for DRP1^35–37^. MFF depletion significantly reduced DRP1 recruitment to both mitochondria and ERMCSs, as measured by colocalization with the ERMCS reporter (SPLICS–ER–Mt), whereas FIS1 depletion had no significant effect **(Extended Data Fig. 8e-g)**. Notably, MFF was also identified as a potential Contact-phagy substrate by both Lyso-IP and ERMCS interactome analyses **(Fig. 3e and Fig. 4d)** and accumulated in CCPG1-deficient cells **(Extended Data Fig. 3i)**. Consistent with these findings, CCPG1-, SIGMAR1- and FIP200-deficient cells displayed increased MFF localization at ERMCSs, as indicated by enhanced colocalization with the ER marker RTN4 **(Extended Data Fig. 8h–i)**. These findings suggest that impaired Contact-phagy promotes persistence of MFF at ERMCSs, thereby favoring sustained DRP1 recruitment at these interfaces.

We then tested whether the mitochondrial fission machinery is functionally required for ERMCS remodeling. Cells lacking DRP1 or FIS1 exhibited a significant increase in ERMCS signals specifically along the VAPB–PTPIP51 axis, as assessed by proximity ligation assays **(Extended Data Fig. 8j-k)**, whereas no significant changes were observed for IP3R3–VDAC1. Consistently, SPLICS–ER–Mt analysis revealed an expanded ERMCS area in DRP1- or FIS1-deficient cells, including under rotenone and starvation conditions **(Extended Fig. 8l-n)**. These results indicate that disruption of the mitochondrial fission machinery impairs efficient remodeling and resolution of ER-mitochondria contacts.

Finally, we tested a catalytically inactive DRP1 mutant K38A, which has been shown to inhibit mitochondria fission^38^ **(Extended Data Fig. 8o)**. Expression of DRP1 K38A increased ER-mitochondria contact sites **(Fig. 4i and Extended Data Fig. 8p-q)** and impaired their remodeling in response to starvation, further confirming a requirement for DRP1-dependent fission in ERMCS dynamics.

Altogether, these findings support a model in which DRP1 is recruited to ERMCS, where it contributes to local membrane remodeling and fission, while MFF facilitates its recruitment to these interfaces. Following this remodeling step, DRP1 together with ERMCS components is targeted for autophagic turnover through Contact-phagy. Accordingly, impairment of Contact-phagy results in persistence of MFF and DRP1 at ERMCSs, altered organization of the fission machinery, and defective resolution of these interfaces.

### Role of Contact-phagy in mitochondrial homeostasis

Because Contact-phagy mediates ERMCS turnover, CCPG1–SIGMAR1-mediated remodeling of these interfaces is expected to influence organelle dynamics and homeostasis. CCPG1 has previously been shown to maintain ER structure and proteostasis in the exocrine pancreas16. Consistently, analysis of insoluble protein fractions from *Ccpg1* hypomorphic (gene-trap, GT) mice revealed accumulation of ER-synthesized enzymes and luminal chaperones^16,17^. Reanalysis of this dataset further revealed upregulation not only of ER proteins but also of components of the ER-Golgi intermediate compartment (ERGIC) and mitochondrial pathways in CCPG1-deficient pancreatic tissue compared with wild-type controls **(Fig. 5a, highlighted in red)**. These findings parallel our proteomic observations in CCPG1-knockout (KO) HeLa cells **(Fig. 3b-c and Extended Data Fig. 3g-h)**.

**Figure 5.**
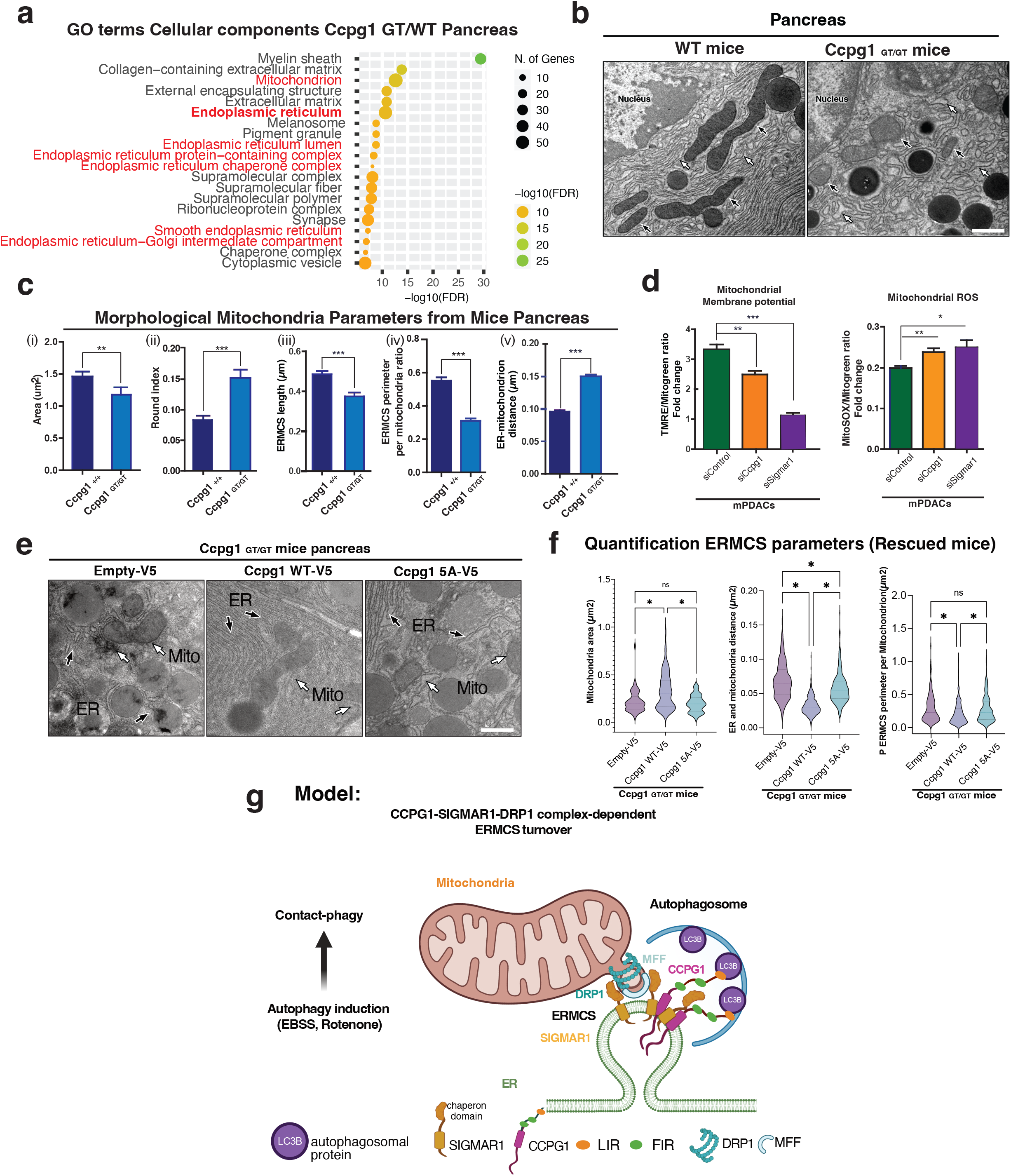
Contact-phagy is important for the maintenance of mitochondrion function and depends of CCPG1-SIGMAR1 complex *in vivo*. **a)** Cellular component pathway analysis of proteins up-regulated in Ccgp1 ^GT/GT^ compared to Ccpg1^+/+^ pancreatic insoluble fraction (Mass spectrometry data from^16^). Depicted Mitochondria and ER cellular components (red) were accumulated in the absence of Ccpg1. **b)** Representative TEM images from WT and Ccpg1 ^GT/GT^ pancreas from 8-week-old mice. ER and mitochondria are indicated by white and black arrows respectively. Scale bar 500 nm. **c)** Morphological mitochondria parameters were analyzed from TEM images (Ccpg1^+/+^: n=3, Ccpg1 ^GT/GT^: n=3 mice per group); **(i)** mitochondria area (**μ**m^2^); **ii)** Round index**; iii)** ERMCSs length (μm), **iv)** ERMCS perimeter per mitochondrion ratio and **v)** ER-mitochondrion distance (μm). **d)** Mitochondrial membrane potential and superoxide levels were measured in murine pancreatic ductal adenocarcinoma cells (mPDACs) in siControl, siCcpg1 and siSigmar1 (number of cells TMRE: n=70, n=62, n=50 and MitoSOX: n=52, n=44, 52, respectively). For this, we used TMRE and MitoSox, respectively, and quantified by analyzing the fluorescence intensity. **e)** Representative electron microscopy images and its morphological analysis from Ccpg1 ^GT/GT^+Empty-V5, Ccpg1 WT-V5 and Ccpg1 5A-V5 mutant expressing pancreas. (Ccpg1^GT/GT^+Empty-V5 n=3, Ccpg1^GT/GT^+Ccpg1 WT-V5 n=3, Ccpg1^GT/GT^+ Ccpg1 5A-V5 n=3 animals per group). **f)** Bar plot results of mitochondria area (µm^2^), ER and mitochondria distance (nm) and ERMCS perimeter per mitochondrion (µm). Data represent the mean ± s.e.m. from three independent experiments; Kruskal–Wallis/Dunn’s post-test. Scale Bar 500 nm. **g)** Model: starvation induces Contact-phagy, SIGMAR1 recruits the ER-phagy receptor CCPG1 to the ERMCSs mediating their degradation through a DRP1-dependent pathway.

Next, immunofluorescence analysis combined with high-content microscopy revealed increased mitochondrial fragmentation in SIGMAR1- and CCPG1-deficient cells, with ~50% more discrete mitochondria per cell **(Extended Data Fig. 9a–b)**. *In vivo*, pancreatic acinar cells from Ccpg1 ^GT/GT^ mice exhibited reduced TOMM20-positive area and increased mitochondrial circularity **(Extended Data Fig. 9e-f)**. TEM revealed pronounced mitochondrial abnormalities **(Fig. 5b)**. Quantitative morphometric analysis showed smaller, more rounded mitochondria **(Fig. 5c, plots i and ii)**, accompanied by reduced ERMCS contact length and ER coverage of the mitochondrial perimeter **(Fig. 5c, plots iii and iv)**, as well as increased ER-mitochondria distance **(Fig. 5c, plot v)**, indicating compromised ERMCS integrity. Similar defects were observed in CCPG1- and SIGMAR1-deficient HeLa cells **(Extended Data Fig. 9c–d, plots i–iv)**, including marked cristae disorganization and mitochondrial swelling **(Extended Data Fig. 9c, arrows)**.

To assess mitochondrial function, murine pancreatic ductal adenocarcinoma (mPDAC) cells were analyzed using live-cell reporters of membrane potential (TMRE) and mitochondrial reactive oxygen species (MitoSOX)^39,40^. Loss of SIGMAR1 or CCPG1 led to a substantial reduction in mitochondrial membrane potential (30–50% relative to control) and increased ROS production (~25%) **(Fig. 5d and Extended Data Fig. 9h)**, with similar effects in HeLa cells **(Extended Data Fig. 9i-j)**. Furthermore, the mitochondrial mass did not change in CCPG1 and SIGMAR1 deficient cells **(Extended Data Fig. 9g)**. Consistent with impaired Contact-phagy, CCPG1-deficient pancreatic acinar cells (mPDACS) failed to degrade ERMCS components **(Extended Data Fig. 9k-l)**, in agreement with proteomic changes in CCPG1 KO HeLa cells **(Fig. 3c and Extended Data Fig. 3g-i)**. Pathway analysis further revealed upregulation of mitochondrial translation, gene expression and metabolic pathways **(Extended Data Fig. 9m)**, underscoring the role of Contact-phagy in mitochondrial homeostasis **(Extended Data Fig. 9n)**.

To test whether SIGMAR1 is required for CCPG1-mediated ERMCS and mitochondrial regulation *in vivo*, we generated a CCPG1 mutant defective in SIGMAR1 binding (Ccpg1^5A^). Based on molecular dynamics simulations, residues within the interfacial (IF; N216, V219) and transmembrane boundary (TM; L215, K217, C218) regions were mutated to alanine **(Fig. 1e and Extended Data Fig. 1e)**. Co-immunoprecipitation and immunofluorescence analyses showed that mutation of either region was sufficient to disrupt CCPG1–SIGMAR1 interaction **(Extended Data Fig. 1j)**. Expression of CCPG1^5A^ reduced mitochondrial area and increased circularity compared with wild-type CCPG1 **(Extended Data Fig. 10a–b)**. *In vivo* rescue experiments demonstrated that re-expression of wild-type *Ccpg1* restored mitochondrial morphology and ERMCS integrity in *Ccpg1* ^GT/GT^ mice, whereas Ccpg1^5A^ failed to fully rescue these defects **(Extended Data Fig. 10c–d)**. TEM analysis confirmed that expression of luciferase control protein in *Ccpg1* ^GT/GT^ mice pancreas did not rescue dilated ER, fragmented mitochondria and reduced ERMCS contacts **(Fig. 5e, left panel)**, while wild-type Ccpg1 restored normal ER and mitochondrial morphology and contact sites **(Fig. 5e, middle panel)**. In contrast and importantly, Ccpg1^5A^ expression resulted in an intermediate phenotype **(Fig. 5e, right panel)**. Quantifications are provided in **Figure 5f**. Notably, the ER luminal Ccpg1 substrate REG3A/B accumulated in the presence of Ccpg1^5A^, indicating that SIGMAR1 binding is required for CCPG1-dependent substrate turnover **(Extended Data Fig. 10e–f)**.

Collectively, these data demonstrate that intact ERMCS remodeling via Contact-phagy is essential for mitochondrial homeostasis **(Extended Data Fig. 9m-n, model)**. Contact-phagy therefore represents a selective autophagy pathway that remodels ER-mitochondria interfaces to regulate mitochondrial dynamics during nutrient deprivation and more specifically during mitochondrial stress supporting cellular adaptation to metabolic stress **(Fig. 5g, model)**.

## Discussion

Here, we identify Contact-phagy as a selective autophagy pathway that promotes turnover of defined ER-mitochondria contact-site (ERMCS) subdomains and contributes to ER and mitochondrial homeostasis. CCPG1 and SIGMAR1 form a receptor-chaperone complex at ERMCS, where SIGMAR1–CCPG1 nanoclusters define sites of ER-phagy initiation and functionally couple local autophagic turnover to the mitochondrial fission machinery. Through this coordination, Contact-phagy links selection of specific contact-site subdomains to remodeling of the apposed membranes and their subsequent lysosomal turnover.

Contact-phagy is not uniformly engaged across ERMCS proteins or stress conditions. Starvation and rotenone promote turnover of defined contact-site components, including VAPB–PTPIP51-associated machinery, whereas tunicamycin does not induce comparable ERMCS degradation. These responses can occur without equivalent activation of bulk autophagy, general ER-phagy or mitophagy, indicating that Contact-phagy does not simply represent a generic consequence of cellular or ER stress. Instead, it appears to constitute a stress-selective remodeling program acting on defined ERMCS subdomains. This selectivity further suggests that ER-mitochondria contacts are heterogeneous structures with distinct molecular compositions, functions and susceptibilities to autophagic turnover.

A key unresolved question is how ERMCS subdomains are selected for Contact-phagy. Metabolic or mitochondrial stress may alter the composition, stability or functional state of particular contacts, thereby favoring their recognition by the SIGMAR1–CCPG1 machinery. Alternatively, prolonged contact persistence or local damage may generate signals that recruit or activate Contact-phagy. Whether persistence itself constitutes a trigger, or instead arises as a consequence of impaired contact-site turnover, remains to be established. Defining the molecular features that distinguish ERMCS destined for turnover from functional contacts will therefore be important for understanding how Contact-phagy selectively remodels the ER-mitochondria interface.

Our findings further reveal that autophagic turnover of ERMCS is closely integrated with mitochondrial fission. DRP1 interacts with SIGMAR1, MFF promotes DRP1 recruitment to ERMCS, and loss of CCPG1 or SIGMAR1 results in abnormal persistence and clustering of DRP1 and MFF at these interfaces. Thus, recruitment of the fission machinery alone is insufficient for productive contact-site remodeling; its spatial coordination with Contact-phagy appears to be critical. These findings suggest that CCPG1–SIGMAR1-dependent autophagic remodeling and DRP1–MFF-dependent mitochondrial remodeling are functionally coupled.

Such coordination addresses an inherent topological challenge of contact-site quality control: selective removal of an interorganelle interface requires remodeling of closely apposed membranes belonging to two distinct organelles. CCPG1-dependent ER-phagy and DRP1-dependent mitochondrial fission may therefore act together to disengage selected ER-mitochondria interfaces and facilitate their autophagic turnover. In this model, Contact-phagy integrates contact-site selection, organelle remodeling and autophagic removal.

The consequences of disrupting this pathway extend beyond contact-site organization. Loss of CCPG1 or SIGMAR1 alters ER architecture and mitochondrial morphology and is associated with increased ROS, impaired mitochondrial membrane potential and reduced bioenergetic capacity. CCPG1 deficiency further perturbs ER-mitochondria organization and mitochondrial function in pancreatic tissue, demonstrating a requirement for this pathway in vivo. Pancreatic ductal adenocarcinoma cells depend on both CCPG1 and SIGMAR1 to maintain organelle integrity and metabolic fitness, suggesting that Contact-phagy becomes particularly important under sustained metabolic and proteostatic demands.

Whether analogous mechanisms operate at other membrane contact sites will be important for establishing Contact-phagy as a broader principle of interorganelle contact-site quality control.

## Materials and Methods

### Mouse breeding

Mice were bred under standard husbandry conditions in i.v.c cages with environmental enrichment. *Ccpg1* mutant animals previously generated ^16^ (considered a non-harmful genetic alteration prior to 1 year of age) were bred under the authority of a UK Home Office Project License held by Simon Wilkinson, subject to local ethical review (University of Edinburgh). The established line was checked on a regular basis for any health concerns.

### Cell Culture and inducible cell lines

Cell lines were cultured in DMEM supplemented with 10% heat-inactivated FBS and penicillin-streptomycin (Thermo Fisher Scientific) in a 5% CO_2_ atmosphere at 37 °C. U2OS, HeLa, HEK293T and HCT116 cells were provided from the American Type Culture Collection (Manassas, VA) and were used to generate inducible cell lines based on lentiviral transduction^41^. U2OS TRex cells were provided by S. Blacklow, Brigham and Women’s Hospital and Harvard Medical School. U2OS-mEGFP-DRP1 and U2OS-Halo-DRP1 cell lines were provided by Ana García-Sáez. Their identities were authenticated by STR analysis. HeLa TRex were provided by Prof. S. Taylor (Manchester University) ^42^, *Hela Control CRISPR KO and CRISPR CCPG1 KO* were previously described ^16^. The mCherry–eGFP–CCPG1 WT constructs were introduced into the vector pcDNA5/FRT/TO using GATEWAY technology and transfected with the Flp recombinase expression plasmid, pOG44, into Flp-In U2OS TRex cells. Following selection with 300 µg/ml hygromycin, the resistant cells were expanded. Using lentiviral transduction, we generated U2OS cell lines expressing HA tagged SIGMAR1-WT under the control of a doxycycline-inducible promoter. Expression was induced with 1 μg ml^−1^ doxycycline for the indicated time. For Cryo-CLEM analysis, we generated stable HCT116 cell lines constitutively expressing mCherryLC3B and GFP-CCPG1 (G418 1 mg/ml and 15 µg/ml blasticidin) or mCherryCCPG1 and TOMM20-GFP (3 ug/ml puromycin, 1mg/ml G418). The lentiviral particles were produced in HEK 293T cells (ATCC). In brief, HEK293T (ATCC) cells were co-transfected with lentiviral transfer plasmids (1.1 µg) encoding either HA-SIGMAR1, mCherry-LC3B, EGFP-CCPG1-WT, or TOMM20-EGFP, together with the packaging plasmids pPAX2 (2.2 µg) and pMD2.G (1.0 µg), using TurboFect reagent (Thermo Fisher Scientific). The lentivirus-containing medium was collected after 48 h. The medium was centrifuged to remove dead HEK293T cells and stored at −80 °C. The medium containing lentiviral particles was transferred to HCT116 cells in the presence of 8 µg/mL polybrene. After 24 h, cells were selected with the appropriate antibiotic.

To generate HCT116 cells constitutively co-expressing EGFP-CCPG1 and mCherry-LC3B, HCT116 cells were first transduced with lentiviral particles carrying the pHAGE (Blasticidin) DEST-EGFP-CCPG1-WT construct. Transduced cells were selected with 15 µg/mL blasticidin. Stable EGFP-CCPG1–expressing HCT116 cells were subsequently transduced with lentiviral particles carrying the pLenti CMV G418 DEST-mCherry-LC3B construct and selected with 1 mg/mL G418. Finally, stable HCT116 cells co-expressing EGFP-CCPG1 and mCherry-LC3B were maintained in culture medium containing 15 µg/mL blasticidin and 1 mg/mL G418. A similar procedure was used to generate HCT116 cells constitutively co-expressing mCherry-CCPG1 and TOMM20-EGFP. Lentiviral constructs pHAGE (puromycin) DEST-mCherry-CCPG1-WT and pLenti CMV G418 DEST-TOMM20-EGFP were used sequentially, with selection using 3 µg/mL puromycin and 1 mg/mL G418. Cells were maintained in the presence of antibiotics for selection and were grown to subconfluence before each experiment.

For cell lines expressing constructs under the control of a doxycycline-inducible promoter, 1 µg/mL doxycycline (Sigma-Aldrich) was used. Cells were treated with 200 ng/mL bafilomycin A1 (LC Laboratories) and EBSS medium (Thermo Fisher Scientific) for the indicated time periods. Transient transfections were performed using Turbofect reagent according to the manufacturer’s instructions. All cell lines were regularly tested for mycoplasma contamination using the LookOut Mycoplasma PCR Detection Kit (Sigma).

Human *CCPG1*-targeting siRNAs (siRNA #1 5′-GGAGAGAGAACGACUAGUA-3’, siRNA #2 5’-UCAGAAGCUCAACAGUUA-3’), human SIGMAR1-targeting siRNAs (siRNA #1 5′-GAAUGCGGGUGGCUGGAUG-3’ and siRNA #2 5’-GGCCUGAGCUCACCACCUA-3’), murine *CCPG1*-targeting siRNAs (J-057836-12-0002; J-057836-10-0002; J-057836-09-0002), SIGMAR1-targeting siRNAs (J-040017-09-0002; J-040017-12-0002; J-040017-12-0002), human FIP200-targeting siRNAs (pooled siRNA#1 5′-GGAGUGGGCUGGUGCUUUA, siRNA#2 5’-AAACUACGAUUGACAUAA, siRNA#3 5’-GCAAAGAAAUUAGGGAAUC, siRNA#4 5’-UAAACUUGACGGACUAAUA), human DRP1-targeting siRNA #5’-GGAGCCAGCUAGAUAUUAA and siRNA #GUGAACCCGUGGAUGAUAA, human FIS1-targeting siRNA #CGAGAAGGCCUUAAAGUAC and siRNA#CAGCAGGCUCGGUGUCCAA, human siRNA targeting MFF (cat. no. L-018261-00-0005) were purchased from Dharmacon. A non-targeting siRNA was used as a control (Qiagen). Transfection with siRNA was carried out using Lipofectamine RNAiMAX (Thermo Fisher Scientific) according to the manufacturer’s instructions. Assays were carried out 72 h post-transfection.

### Generation of mouse pancreatic ductal acinar cell knockouts

Mouse PDAC cells were infected with lentiCas9-Blast and selected with blasticidin. The resulting PDAC-Cas9 cells were then infected with lenti-guide puro (either empty vector or with Ccpg1 gRNA target sequence TTGCGGCGCCGTCTAGGAGC cloned in as per standard protocol). Infected cells were selected with puromycin and then from the resultant pool individual wild-type or Ccpg1 knockout clones were obtained by limiting dilution. lentiGuide-Puro was a gift from Feng Zhang (Addgene plasmid # 52963; http://n2t.net/addgene:52963; RRID: Addgene_52963). LentiCas9-Blast was a gift from Feng Zhang (Addgene plasmid # 52962; http://n2t.net/addgene:52962; RRID: Addgene_52962).

### Generation of Contact-phagy reporter cell lines

Lentivirus (derived from plasmids pLX304 and pLIX) were produced in HEK293FT cells by transient transfection with packaging plasmids, as previously described (https://www.mdpi.com/2073-4409/12/8/1134). Viruses were collected 48 h post-transfection in high-serum full medium (30% FBS) and filtered through a 0.45 µm filter. Viruses were then concentrated using Lenti-X Concentrator (Takara Bio, 631231), combining one volume of concentrator with three volumes of clarified supernatant. The mixture was incubated for 16 h at 4°C, centrifuged at 1,500 g for 45 min at 4°C, and the resulting pellet was resuspended in PBS to a final concentration ten times that of the original supernatant volume. For transduction, HeLa cells were infected for 2 d with 10 µl viruses/ml of media in the presence of 10 µg/mL polybrene (Sigma-Aldrich, TR-1003), then sorted by FACS.

### Flow Cytometry for Contact-phagy receptors sorting

Cells were sorted using a FACSAria II SORP (BD Biosciences, San Jose, CA, USA). TOLLES was excited with a 405 nm laser and its emission collected through a 525/50 BP filter; YPet was excited with a 488 nm laser and its emission collected through the same 525/50 BP filter. Single cells were first gated using FSC-Height (FSC-H) versus FSC-Area (FSC-A), then gated for a narrow range of YPet and TOLLES expression prior to collection. Data were recorded using BD FACSDiva software (v6.3.1).

### Antibodies

**Primary and secondary antibodies are presented in Supplementary Table 1**.

| Antigen. | Company and ID | Application and dilution |
|---|---|---|
| GAPDH (14C10) | Cell signalling # 2118 | WB (1/5000) |
| HA | Roche (11867423001) | WB (1/10,000), IF (1/2000) |
| LC3B | Rabbit mAb #3868 CST | WB 1/1000; IF (1/500) |
| REEP5 | Proteintech (14643-1-AP) | WB, IF (1/1000) |
| FLAG (M2) | Sigma (F3165-5MG) | WB (1/10000), IF (1/1000) |
| SIGMAR1 | Proteintech (15168-1-AP) | WB (1/1000) |
| CCPG1 | Proteintech (13861-1-AP) | WB (1/2000) |
| CCPG1 | Generated in house, <sup>16</sup> | IF (1/100) |
| GFP | Clontech (Cat. 632460) | WB (1/1000) |
| mCherry | Eurogen (AB233) | WB (1/1000) |
| IP3R3 | BD Biosciecnes (610312) | IF (1/300), |
| VDAC1 | Proteintech (55259-1-AP) | IF (1/300, only PLA), WB 1/2000 |
| VAPB mouse | Proteintech (66191-1-Ig) | IF (1/300) |
| PTPIP51 rabbit | Proteintech (2064-1-AP) | IF (1/300) |
| TOMM20 rabbit | Proteintech (11802-1-AP) | IF (1/300), WB 1/2000 |
| RTN4 sheep | Biothechne AF6034 | IF (1/300) |
| TOMM20 mouse | BD Biosciences (612278) | IF (1/300) |
| DRP1 rabbit | BD Biosciences (611738) | IF (1/300), WB 1/2000 |
| FIS1 rabbit | Proteintech 10956-1-AP | IF (1/300), WB 1/2000 |
| MFF rabbit | Proteintech 17090-1-AP | IF (1/300), WB 1/2000 |
| HRP-conjugated secondary antibodies<br>anti-mouse, anti-rabbit and anti-rat | Cell Signaling, #7077S, Jackson ImmunoResearch #309-005-003, AB_2307391 B_2338128 | WB (1/10000) |
| anti-rabbit Alexa 647 | A21244, Life Technology | IF (1/500) |
| anti-mouse Cy3<br>anti-rat Cy3 | MerckMillipore, #AP124C,<br>MerckMillipore #AP189C | IF (1/500) |
| Vinculin | Sigma (V4505) | WB (1/5000) |
| anti-rabbit Cy5<br>anti-sheep Alexa 555 | Invirogen A10523<br>Invitrogen, A21436 | IF (1/500) |
| Anti myc tag | Cell signalling #2276 | WB (1/5000) |
| V5 – Mouse | Invitrogen 14-6796-82 | IF (1/500) |
| REG3A/G – Mouse | Santacruz sc-377038 | IF (1/500) |
| V5 – Rabbit | BioLegend 903801 | IF (1/500) |
| AlexaFlour488 anti-rabbit | Invitrogen A21206 | IF (1/500) |
| AlexaFluor594 anti-mouse | Invitrogen A11005 | IF (1/500) |
| AlexaFlour488 anti-mouse | Invitrogen A11001 | IF (1/500) |
| AlexaFluor594 anti-rabbit | Invitrogen A11012 | IF (1/500) |

### Plasmids

We introduced the cDNAs into vector pDONR223 using the BP Clonase Reaction Kit (Invitrogen) followed by transfer to GATEWAY destination vectors using the LR Clonase Reaction Kit (Invitrogen), resulting in vectors pLTD-N-HA-PURO, pcDNA5-FRT/TO-N-mCherry-eGFP, pDEST-V1ORF, pDEST-V2-ORF, pDEST-ORF-V1 and pDEST-ORF-V2.

Site-directed mutagenesis of *Ccpg1* was carried out using a two-step Polymerase Chain Reaction (PCR) method, using primers containing the corresponding mutations and the KOD polymerase (Sigma, 71842-3). Constructs were then cloned into a pDONR223 destination by BP reactions (Thermo Fisher Scientific, 11789020). Then, constructs were cloned into AAV vectors by LR reaction (Thermo Fisher Scientific, 11791020).

**Plasmids are presented in Supplementary Table 2**.

| Plasmid | Mutation / Tagging | Usage | Source |
|---|---|---|---|
| pDONR233-SIGMAR1 | WT | GATEWAY entry vector for pDEST plasmids. | This study |
| pDONR233-CCPG1 | WT | GATEWAY entry vector for pDEST plasmids. | <sup>16</sup> |
| pDONR233-TOMM20 | TOMM20 WT | GATEWAY entry vector for pDEST plasmids. | This study |
| pDONR233-PTPIP51 non-stop | PTPIP51 | GATEWAY entry vector for pDEST plasmids. | This study |
| pDONR233-VAPB | WT VAPB | GATEWAY entry vector for pDEST plasmids. | This study |
| pLTD-HA-SIGMAR1 | N-terminal -HA tag | Expression under the control of a doxycycline inducible promoter.<br>Lentiviral vector | This study |
| GFP-CCPG1 | N-terminal-HA tag | Transient transfection | <sup>16</sup> |
| pcDNA5-FRT/TO-N-mCherry-EGFP-CCPG1-WT | N-terminal mCherry-EGFP | Expression under the control of a doxycycline inducible promoter | This study |
| pcDNA3.1. 6xmyc-CCPG1-WT | N-terminal Myc-tag (1-757aa) | Transient transfection | This study |
| pcDNA3.1. 6xmyc-CCPG1-NTD | N terminal-Myc-tag, N-terminal domain of CCPG1 (1-230aa) | Transient transfection | This study |
| pcDNA3.1 6xmyc-CCPG1-delta TM | N terminal-Myc-tag, delta TM on CCPG1 (1-757aa with the deletion between 217 and 235aa). | Transient transfection | This study |
| pcDNA3.1 6xmyc-CCPG1-CTD | N terminal-Myc-tag, C-terminal domain of CCPG1 (231 and 757 aa). | Transient Transfection | This study |
| pcDNA3.1 SFB-FLAG-CCPG1. (SFB: S-Tag, 2xFLAG-Tag, Streptavidin-Binding Peptide (SBP)-Tag | N terminal-SFP and FLAG tag | Transient transfection | This study |
| PHAGE-mCherrySIGMAR1 | N terminal-mCherry tag | Transient transfection | This study |
| pAcmCherryDRP1 WT | N-terminal mCherry tag, DRP1 WT (derived from mCherry substituted YFP-Backbone Takara) | Transient Transfection | Addgene #49152 |
| pAcmCherryDRP1 K38A | N-terminal tag, mCherry DRP1 K38A catalytic inactive mutant (derived mCherry Substituted YFP-Backbone Takara) | Transient Transfection | Addgene #232350 |
| V1-SIGMAR1 | N-terminal V1, which is a non-fluorescent N-terminal of the Venus (Met1-Gln157). | Transient Transfection | This study |
| V2-SIGMAR1 | N-terminal V2, which is a non-fluorescent C-terminal of the Venus (Lys158-Lys238) | Transient Transfection | This study |
| pDONOR223 TOMM20-GFP | Gateway entry vector | for pDEST plasmids | This study |
| pHAGE (puro) DEST-mCherry-CCPG1 | N-terminal mCherry tagged to CCPG1 WT full length | Lentiviral production for mammalian constitutive expression | This study |
| pLenti CMV (G4180) DEST-TOMM20-EGFP | C-terminal GFP tagging TOMM20 | Lentiviral production for mammalian constitutive expression | This study |
| pLenti CMV (G418) DEST-mCherry LC3B | N-terminal mCherry tagged LC3B | Lentiviral production for mammalian constitutive expression | This study |
| PHAGE-GFP-CCPG1 (Blast) | N-terminal GFP tagged CCPG1 with | Lentiviral production | This study |
| pDONR223 mCherry-LC3B | Gateway entry vector | for pDEST plasmids | This study |
| pLenti CMV Hygro DEST-SPLICs ER-Mt | Detect the short Mitochondria-Endoplasmic Reticumul contact (8-10nm) | Lentiviral production for mammalian constitutive expression | This study |
| RFP-GFP-KDEL | N-terminal tagged RFP and GFP tandem of KDEL (ER retention sequence) | Lentiviral production for generation of ER-phagy reporter Hela cells | Addgene #128257 |
| pDONR223 Ccpg1<br>Mouse Ccpg1 (1-753) | WT CCPG1 | GATEWAY entry vector for pDEST plasmids. | Bioscience |
| pDONR223 Ccpg1 SIGMAR_IF<br>Mouse Ccpg1 | CCPG1 Interfacial (IF) Mutant residues N217A, V220A that impairs SIGMAR1 binding | GATEWAY entry vector for pDEST plasmids. | This study |
| pDONR223 Ccpg1 SIGMAR_TM<br>Mouse Ccpg1 | CCPG1 Transmembrane (TM) Mutant residues L216A, K218A, C219A that impairs SIGMAR1 binding | GATEWAY entry vector for pDEST plasmids. | This study |
| pDONR223 Ccpg1 SIGMAR_5A<br>Mouse Ccpg1 | CCPG1 5A mutant (5A)<br>Residues mutated in IF and TM (216-220 A5) that impairs SIGMAR1 binding | GATEWAY entry vector for pDEST plasmids. | This study |
| AAV Ela-DEST-V5 | Gateway destination vector for AAV production | Adeno-associated virus (AAV) production with the acinar-specific Elastase promoter. | This study |
| AAV Ccpg1-V5<br>Mouse Ccpg1 (1-753) tagged to V5 | WT CCPG1 tagged with V5 | Adeno-associated virus (AAV) production with the acinar-specific Elastase promoter. | This study |
| AAV Ccpg1 SIGMAR_IF-V5<br>Mouse Ccpg1 N217A, V220A tagged to V5 | CCPG1 Interfacial (IF) Mutant residues N217A, V220A that impairs SIGMAR1 binding tagged with V5 | Adeno-associated virus (AAV) production with the acinar-specific Elastase promoter. | This study |
| AAV Ccpg1 SIGMAR_TM-V5<br>Mouse Ccpg1 L216A, K218A, C219A tagged to V5 | CCPG1 Transmembrane (TM) Mutant residues L216A, K218A, C219A that impairs SIGMAR1 binding tagged with V5 | Adeno-associated virus (AAV) production with the acinar-specific Elastase promoter. | This study |
| AAV Ccpg1 SIGMAR_5A -V5<br>Mouse Ccpg1 (216-220 A5) tagged to V5 | CCPG1 5A mutant (5A)<br>Residues mutated in IF and TM (216-220 A5) that impairs SIGMAR1 binding tagged wth V5 | Adeno-associated virus (AAV) production with the acinar-specific Elastase promoter. | This study |
| pLIX 403 TRE-SRAI-CCPG1 (Dox inducible) | Gateway destination vector for inducible expression of human CCPG1 fused to N-terminal SRAI | Lentiviral production for generation of Contact-phagy reporter Hela cells | This paper, modified from Addgene plasmid #41395 (a gift from David Root) |
| pDONR223 LC3B | Human LC3B | Lentiviral production for generation of Contact-phagy reporter Hela cells | <sup>28</sup> |
| pLX304 SRAI-C1 DEST | Gateway destination vector | Lentiviral production for generation of Contact-phagy reporter Hela cells | <sup>28</sup> |
| pLX304 SRAI-N1 DEST<br>Gateway destination vector | Gateway destination vector | Lentiviral production for generation of Contact-phagy reporter Hela cells | <sup>28</sup> |
| pLX304 SRAI-SIGMAR1 | Human SIGMAR1 fused to N terminal SRAI | Lentiviral production for generation of Contact-phagy reporter Hela cells | This study |
| pLX304 SRAI-PTPIP5 | Human PTPIP51 fused to N terminal SRAI | Lentiviral production for generation of Contact-phagy reporter Hela cells | This study |
| pLX304 SRAI-VAPB | Human VAPB fused to N terminal SRAI | Lentiviral production for generation of Contact-phagy reporter Hela cells | This study |
| pLX304 TOMM20-SRAI | Human TOM20 fused to C terminal SRAI | Lentiviral production for generation of Contact-phagy reporter Hela cells | This study |
| pLX304 SRAI-LC3B | Human LC3B fused to N terminal SRAI | Lentiviral production for generation of Contact-phagy reporter Hela cells | <sup>28</sup> |
| pLX304 ss-SRAI-KDEL | ss= N-terminal ER signal sequence from IgGκSP<br>KDEL= C-terminal retaining ER sequence | Lentiviral production for generation of Contact-phagy reporter Hela cells | <sup>28</sup> |

### Recombinant adeno-associated virus generation

Adeno-associated virus (AAV) serotype 2/8 were prepared as previously described^43,44^. Briefly, AAVpro 293T cells (Takara Bio, #632273) were plated in 15 cm dishes and transfected using Opti-MEM (Gibco, #31985070) and PEI-Max (Polysciences, #24765), at a ratio of 1:1:1 AAV plasmid, pHelper (a gift from L. Boulter, University of Edinburgh) and pAAV2/8 (gift from C. David, Tsinghua University). Viral particles were collected from culture media and cell lysates. Conditioned media was collected in falcon tubes and cleared by centrifugation at 800 × g for 10 min at 4 °C. Cells were scraped into 12 ml of cold PBS, centrifuged at 1,500 × g for 5 min and resuspended in 5 ml lysis buffer (10 mM Tris-HCl pH 8.5, 150 mM NaCl). Lysis was performed by 3 freeze-thaw cycles, samples were frozen in dry ice alternating with heating at 37 °C. After centrifugation at 3,000 × g for 10 min, supernatant was combined with original cell culture medium. 1 volume of concentrator solution (40% PEG-8000, 1.2 M NaCl, PBS pH 7.4) was then added to 3 volumes of viral particles and incubated overnight at 4 °C rotating. The following day, samples were centrifuged at 1600 × g for 60 min at 4 °C. Viral pellets were resuspended in 1 ml cold PBS and incubated overnight at 4 °C. The next day, samples were centrifuged at 13,000 × g for 3 min at 4 °C and supernatants were retained. The final AAV-containing supernatants were titered using qPCR as described^45^ then aliquoted and frozen prior to use.

### CCPG1 GT/GT rescue experiment

Delivery of rAAV expressing *Ccpg1* isoforms or *Luciferase*-V5 negative control to mouse pancreas was performed via a single intraperitoneal injection of AAV preparations (20 ml/kg, 1.5 × 10^9^ viral genome / μL). Mice were monitored to ensure full recovery prior to return to general husbandry. Mice were terminated at 40 days post-injection by cervical dislocation. Freshly excised pancreata was either snap frozen for posterior protein extraction or fixed by immediate immersion in 10% neutral buffered formalin and incubation overnight at RT. The following day fixed samples were washed 2 × 70% EtOH and embedded in paraffin.

### Interactome analysis and sample preparation for MS

For the LC–MS interactome analysis, HA–SIGMAR1 expression was induced in U2OS cells with 1ug/ml doxycycline. After 24 h, cells were lysed (50 mM Tris-HCl pH 7.5, 150 mM NaCl, 0.5 mM ethylenediaminetetraacetic acid (EDTA), 1% Triton X-100, 10 mM N-ethylmaleimide (NEM) and Roche EDTA-free protease inhibitors, freshly added). Next, the samples were incubated with HA-agarose beads (Sigma-Aldrich, A2095) overnight at 4 °C on a rotating platform. Protein-bound HA-beads were washed three times with lysis buffer Samples were prepared for LC-MS analysis. HA-IP samples were incubated with 40 µl denaturing buffer (2% sodium deoxycholate, 1 mM tris(2-carboxyethyl) phosphine, 4 mM chloroacetamide and 50 mM Tris-HCl pH 8.5) and heated at 95 °C for 10 min. Samples were mixed 1:1 with 500 ng LysC (Promega), incubated for 3 h at 37 °C and digested with 500 ng of trypsin in 50 mM Tris-HCl, pH 8.5, overnight at 37 °C. Reactions were stopped by addition of 150 µl of isopropanol containing 1% trifluoroacetic acid (TFA). Peptides were cleaned up by loading them onto SDB-RPS stage tips (Sigma). After one wash with 1% TFA in isopropanol and one wash with 0.2% TFA in water, peptides were eluted using 80% acetonitrile and 1.25% ammonia. Eluted peptides were dried and processed for LC–MS measurements.

For endogenous co-immunoprecipitation of CCPG1 and SIGMAR1, confluent 15 cm dishes of HeLa cells were harvested in 1% Triton X-100 IP buffer (50 mM Tris-HCl, 150 mM NaCl, 0.5 mM EDTA) supplemented with complete protease inhibitor cocktail (Roche) and 1 mM PMSF. Lysates were cleared by centrifugation at 12,000 × g for 10 min and incubated overnight at 4°C with CCPG1 primary antibody. Lysates and antibody were then incubated with Protein G–Agarose beads (Roche) at 4°C. Beads were washed three times with lysis buffer, resuspended in Laemmli buffer, and boiled. The resulting supernatants were loaded onto SDS-PAGE.

### Modeling and simulations of SIGMAR1–CCPG1 complexes

Sequences corresponding to the human CCPG1_197-250_ (UniProt code: Q9ULG6) and full-length SIGMAR1 (UniProt code: Q99720) were stitched with a polyglycine linker (40 residues) and modeled as a single polypeptide using AlphaFold (v2.0 with default parameters and the database dated 05-14-2020)^46^. To remove any bias arising from the directional stapling of the two proteins, we reversed their orientation, built 10 models, and ranked them. The top-ranked models in both orientations were consistent, showing the same binding mode and interface; therefore, we used the top-ranked model in subsequent MD simulations. We constructed an extended model of CCPG1 including the N-terminal IDR. We used the hierarchical chain growth (HCG) method to build the IDR segment^47^ and grafted it onto the TM fragment CCPG1_197-250_, using overlapping residues to provide relative orientation. The linked structure was first optimized using Chimera’s loop refinement module ^48^, followed by energy minimization of the entire complex with SIGMAR1.

All-atom molecular dynamics simulations of the SIGMAR1-CCPG1_197-250_ complex were performed with GROMACS v2021.5 ^49^. The initial membrane orientation of the complex was predicted using the PPM server ^50^. To assess the robustness of the complex in different membrane environments, the top-ranked SIGMAR1-CCPG1_197-250_ model was embedded in two bilayer compositions: (i) a pure POPC bilayer, used as a minimal PC-rich membrane model, and (ii) an ER-like lipid mixture composed of POPC/DOPC 25.47%, POPE/DOPE 15.10%, POPI 9.44%, POPS 3.77%, DOPS 2.83%, POPA/DOPA 0.94%, and cholesterol 0.94%^51^.

In both the membrane systems, the PPM-predicted orientation was retained, placing the amphipathic helix of the SIGMAR1-Cupin domain in contact with the upper leaflet to optimize membrane insertion and avoid unfavorable initial contacts. Each system was equilibrated and simulated using the same all-atom MD protocol. Independent replicate simulations (*n =* 2) were performed for each membrane condition, and the resulting trajectories were used to compare complex stability, protein orientation, and interaction-site reproducibility across lipid compositions. The protein-membrane systems were built using the CHARMM-GUI server^52^, and interactions were modeled with the CHARMM36m force field^53^. Protein protonation states for CCPG1 and SIGMAR1 were assumed at physiological pH=7, following CHARMM-GUI default parameters. Protein complexes were embedded in a pre-equilibrated square POPC and ER bilayer (16 × 16 nm^2^) under periodic boundary conditions and solvated with the TIP3P water model and NaCl (0.15 M). Electrostatic interactions were modeled using the particle mesh Ewald summation method with a minimum cutoff of 1.2 nm. The LINCS algorithm was employed to constrain covalent bonds between hydrogen and heavy atoms. Van der Waals forces were treated using the Verlet cutoff scheme force-switch modifier with a cutoff of 1.2 nm.

The MD simulations followed the traditional protocol suggested by CHARMM-GUI ^52^. Initially, the system was minimized using the steepest-descent algorithm until the maximum force was reduced to 1000 kJ nm^−1^. Then, two different equilibrations were performed assuming NVT (two steps) and NPT (four steps) ensembles. The first equilibration run used the Berendsen thermostat at 310 K (*τ*_T_ = 1 ps)^54^. A semi-isotropic Berendsen barostat^54^ at 1 bar (*τ*_P_ = 5 ps, compressibility of 4.5 × 10^−5^ bar^−1^) was employed from the 2^nd^ equilibration step. Positional restraints were applied to the phosphate groups of POPC lipids and to the side-chain dihedrals of protein residues during the initial equilibration and were gradually removed during subsequent equilibration steps. For the production run, the pressure was maintained at 1 bar (*τ*_P_ = 5 ps, compressibility of 4.5 × 10^−5^ bar^−1^) with the semi-isotropic Parrinello-Rahman barostat ^55^. Production runs were performed for 1000 ns with a 2 fs timestep. Two replicates (*n* = 2) were performed for each system, using different initial velocities at 310K.

Coarse-grained MD simulations were performed using the Martini v2.2. We converted the atomistic models to a coarse-grained representation using the *martinize.py* script (v2.6)^56^. Local secondary structure was retained by enforcing backbone restraints derived from DSSP assignments^57^. To accurately capture the IDR dynamics of the N-terminal part of the CCPG1_1-250_ model, we used an adapted MARTINI model, reducing the IDR interactions by 40 % ^58,59^. Changes for the coarse-grained proteins were set assuming a pH equal to 7. The protein complexes were embedded in square membrane patches 25 × 25 nm^2^ with *insane.py* ^60^. Additional CG systems were designed to study the effect of SIGMAR1 on CCPG1 clustering. We placed eight copies of CCPG1_1-250_ molecules on a square grid and embedded them in a POPC patch of 45 × 45 nm^2^ (negative control). Using this configuration as a base system, we built additional new systems with increasing concentrations of SIGMAR1. SIGMAR1 molecules were gradually increased from *n*_Prot_ = *k_i_* = [0, 1, 2…, 8] by keeping the inter-protein distance at a distance of at least 10 nm.

The MD runs for all CG systems were simulated using GROMACS (v2021.5;^48^). We employed a reaction field (*ε*_rf_ = 15) with a Coulomb cutoff distance of 1.1 nm to describe the long-range electrostatic interactions. Reaction-field electrostatics were used to enable efficient sampling of the large membrane-associated assemblies, rather than to resolve atomistic details of long-range electrostatic interactions. van der Waals interactions were treated with a cutoff of 1.1 nm and a potential-shift-Verlet modifier. We followed a classical coarse-grained MD protocol in which the systems were first minimized in an initial run using a soft-core potential, followed by minimization without positional restraints. Both minimizations were run for 3000 steps with the steepest-descent algorithm. Then, the equilibration of the system was accomplished by first simulating under the NVT conditions with the velocity-rescale thermostat at 310 K (*τ*_T_ = 1.0 ps) ^61^, followed by 5 runs under NPT conditions including the semi-isotropic Berendsen barostat at 1 bar (*τ*_P_ = 12.0 ps, compressibility of 3.0 × 10^−4^) ^54^. During the NPT runs, positional restraints on the BB (protein part) and PO4 (phosphate groups of POPC lipids) beads were gradually released while the integration time step (*dt) was increased* from 2 fs to 20 fs. Production runs were simulated with a timestep of 20 fs with the velocity-rescale thermostat ^61^ and Parrinello-Rahman barostat ^55^ for 10 µs. Three independent replicas per CG system were generated by assigning different random initial velocities. The resulting CG simulations were used to generate experimentally testable predictions regarding CCPG1– SIGMAR1 binding and CCPG1 oligomerization, which were subsequently evaluated using independent biochemical assays.

Contact maps for all systems in atomistic and coarse-grained representation were computed using in-house scripts implementing MDAnalysis ^62^. The pairwise residue contacts across SIGMAR1 (**A**) and CCPG1 (**B**) were computed as 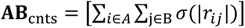, where the sums extend over heavy atom positions of interacting residues (*ij*) of proteins A and B, and 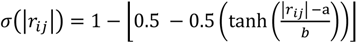, a smooth sigmoidal counting function was implemented to restrict interactions to distances below a cutoff. The cutoff parameter *a* was set to 5 Å for atomistic simulations and 10 Å for coarse-grained simulations. The smoothing parameter *b* was set to 0.5 and 1.0 for atomistic and CG simulations, respectively. Contact maps were averaged over 1000 and 2500 individual frames of the atomistic and coarse-grained MD trajectories, respectively.

We used a hierarchical clustering approach to define oligomeric clusters of CCPG1_1-250_. A PPI interface was considered valid and bound if it exhibited more than 10 contacts. This approach allowed us to progressively identify higher-order assemblies and define the corresponding oligomeric states. The trajectory was partitioned so that CCPG1 molecules were categorized as bound (with variable oligomeric-state annotations *n*-mers) or unbound, to compute the average residence time 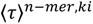 for each oligomeric state (ORT) of CCPG1_1-250_ clusters. Subsequently, the ratio of the average oligomeric residence time for each CCPG1 *n*-mer as 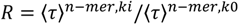 was estimated to quantify the effect of SIGMAR1 on the formation of higher-order CCPG1_1-250_ complexes.

### Blue-native PAGE, SDS–PAGE and Western blot

Blue-native PAGE was performed following the manufacturer’s instructions (#BN1001BOX, protocol pub. no. MAN0007893 Rev. A.0, Life Technology). For SDS–PAGE, proteins were denatured at 90 °C for 5 min in Laemmli buffer. Native or denatured proteins were transferred to methanol-activated polyvinylidene fluoride membranes (Amersham Hybond P, 0.45 µm). Membranes were blocked for 1 h in 5% skimmed milk in 1xPBS and incubated overnight at 4 °C with the specified primary antibody, and for 2 h with the corresponding secondary antibody at room temperature. We used horseradish peroxidase-conjugated anti-rabbit (1:10,000), anti-mouse (1:10,000), and anti-rat (1:10,000) secondary antibodies as appropriate, followed by signal development using Western ECL substrate (sc-2048, Santa Cruz) and the Chemidoc automated detection system (Bio-Rad). Densitometric quantification of western blot bands was carried out using ImageJ (version 1.51w) and Bio-Rad Image Lab software.

### V1-SIGMAR1/V2-SIGMAR1 and V1-VAPB-PTPIP51-V2 BiCAP and mCherry-DRP1 sample preparation for MS

HEK 293T and HeLa cells were transiently co-transfected with the constructs V1– SIGMAR1 and V2–SIGMAR1, V1-VAPB-PTPIP51-V2 and mCherry DRP1. After 16 h, cells were lysed with 1% Triton X-100 in lysis buffer (50 mM Tris-HCl pH 7.5, 150 mM NaCl, 0.5 mM EDTA, Roche EDTA-free protease inhibitor cocktail and NEM), followed by incubation with either GFP-trap or RFP-trap beads (Chromotek/Proteintech) overnight at 4 °C on a rotating platform. Protein-bound beads were washed three times with lysis buffer and one time in the same buffer without detergents before on-bead trypsin digestion. Samples were incubated with 25 µl SDC buffer (2% sodium deoxycholate, 1 mM TCEP, 4 mM chloroacetamide and 50 mM Tris-HCl pH 8.5) and heated to 60 °C for 30 min. We then added 500 ng trypsin in 25 µl 50 mM Tris-HCl (pH 8.5) to each sample and incubated overnight at 37 °C. The reaction was stopped with 150 µl of 1% TFA in isopropanol. Peptides were cleaned up using SDB-RPS stage tips (Sigma-Aldrich). After one wash with 1% TFA in isopropanol and one wash with 0.2% TFA in water, peptides were eluted in 80% acetonitrile plus 1.25% ammonia. Eluted peptides were dried and processed for LC-MS.

### LC-MS analysis (for co-immunoprecipitations)

Dried peptides were reconstituted in 2% acetonitrile containing 0.1% TFA and analyzed on a QExactive HF mass spectrometer coupled to an easy nLC 1200 (Thermo Fisher Scientific) fitted with a 35-cm, 75-µm ID fused-silica column packed in house with 1.9-µm C18 particles (Reprosil pur, Dr. Maisch). The column was maintained at 50 °C using an integrated column oven (Sonation). HPLC solvents consisted of 0.1% Formic acid in water (Buffer A) and 0.1% Formic acid, 80% acetonitrile in water (Buffer B). Peptides were eluted in a non-linear gradient of 4–38% Buffer B over 60 min followed by a step-wise increase to 95% Buffer B in 6 minutes which was held for another 9 minutes and directly sprayed into the mass spectrometer equipped with a nanoFlex ion source (Thermo Fisher Scientific). Full-scan MS spectra (300– 1,650 m/z) were acquired in profile mode at a resolution of 60,000 at m/z 200, a maximum injection time of 20 ms and an AGC (automatic gain control) target value of 3 × 106. Up to 10 of the most intense peptides per full scan were isolated using a 1.4-Th window for fragmentation by higher energy collisional dissociation (normalized collision energy of 27). MS/MS spectra were acquired in centroid mode with a resolution of 30,000, a maximum injection time of 54 ms, and an AGC target value of 1 × 105. Single charged ions, ions with a charge state of more than six, and ions with unassigned charge states were not considered for fragmentation, and dynamic exclusion was set to 20 s to minimize the acquisition of fragment spectra representing already acquired precursors.

### MS data processing (for co-immunoprecipitations)

MS raw data were processed using MaxQuant v1.6.17.0 with default parameters. Acquired spectra were searched against the human ‘one sequence per gene’ database (Taxonomy ID 9606) downloaded from UniProt (12 March 2020; 20,531 sequences), and a collection of 244 common contaminants (‘contaminants.fasta’ provided with MaxQuant v1.6.17.0) using the Andromeda search engine integrated into MaxQuant v1.6.17.0 ^63,64^. Identifications were filtered to obtain false discovery rates below 1% for both peptide spectrum matches (minimum length of seven amino acids) and proteins using a target–decoy strategy^65^ protein quantification and data normalization relied on the MaxLFQ algorithm implemented in MaxQuant v1.6.17.0 ^66^. For protein assignment, spectra were correlated with the UniProt human database v2019, including a list of common contaminants. Searches were performed with tryptic specifications and default settings for mass tolerances in MS and MS/MS spectra. Carbamidomethyl cysteine, methionine oxidation, and N-terminal acetylation were defined as fixed modifications. The match-between-run feature was used with a time window of 1 min. For further analysis, Perseus v2.0.7.0 was used and first filtered for contaminants and reverse entries, as well as proteins that were only identified by a modified peptide.

### Sample preparation of whole cell proteome (WCP)

HeLa CRISPR control and CCPG1 KO CRISPR cells were seeded in 6-well plates and, the following day, were either left untreated or treated with EBSS for 6 hours. Cells were then washed three times with PBS and lysed in 150 µL of lysis buffer (2% SDS, 50 mM Tris-HCl pH 8.5, 10 mM TCEP, 40 mM CAA, protease inhibitors [Roche], and phosphatase inhibitors [Roche]). Lysates were collected and transferred into 2 mL low-binding tubes. Samples were sonicated for 2 min (1 sec on, 1 sec off, 45% amplitude) and boiled for another 5 min at 95 °C. Solubilized proteins were precipitated as described next: 600 μl of ice-cold MeOH was mixed for 2 seconds, and 150 μl of chloroform was added and mixed for 2 more seconds. 3 volumes of water were added and mixed again. Samples were spun for 15 min at 15-20000 g at 4 °C. The first top layer was removed without disturbing the interface, and 750 μl ice-cold methanol was added with a subsequent mix. Samples were then centrifuged for 10 min, as mentioned above. Supernatant was discarded and samples pellet to dry. Pellets were dissolved in 100ul of digestion buffer (8mM Urea, 50mM Tris pH8) to reach 1mg/ml protein concentration. Pellets were resuspended by vortexing. Incubation of 10 minutes at RT at 750rpm on thermomixer. Protein concentration was calculated with BCA method (sample dilution 1:10) and adjusted to 1 μg/ μl. 50ug of total protein was transferred to a new tube. Then, the sample were diluted 1:6 with 50 mM Tris pH8.5. Protein digestion was done adding Lysine-C (1:50 dilution from stock 2 μg/ μl, 10 μl/mg protein) and trypsin (1:100 dilution, stock 0.5 μl/ μl, 20 μl/mg protein) at 37 °C and 550 rpm overnight. The digestion was stop with TFA (final concentration 1%). Cleaning and concentration of the peptides was performed Sep-Pak or StageTip. Perform microBCA assay or peptide assay (both from Thermo/Pierce). For optimal accuracy, resuspend the peptides in around half the volume of 100mM EPPS pH8.2, 10% acetonitrile (ACN) in regard to digested protein (e.g. 25μl if you digested 50μg). Adjust all samples to 1μg/μl with 100mM EPPS pH 8.2, 10% acetonitrile (ACN).

### Sample preparation for Crude Mitochondrial Proteome

HeLa CRISPR control and CCPG1 KO CRISPR cells were seeded in 15cm plates (3x per conditions) and, the following day, were either left untreated or treated with EBSS for 6 hours. For crude mitochondrial isolation, the manufacturer’s instructions for the Thermo Fisher Mitochondria Isolation Kit (Cat. No. 89874) were followed. Cells were washed with 1× PBS to remove debris and media and then centrifuged. The resulting pellet was resuspended in mitochondrial isolation reagent A, briefly vortexed, and incubated on ice for 2 minutes. The sample was then transferred to a Dounce homogenizer, and a few strokes were performed on ice to disrupt the cells without causing complete lysis. Mitochondrial isolation reagent C was added, and the lysate was subjected to differential centrifugation at low speed (700 × g for 10 min at 4°C) to pellet nuclei and cell debris. The supernatant was transferred to a new tube and centrifuged at higher speed (10,000 × g for 15 min at 4°C) to pellet the crude mitochondria. The supernatant was removed and the pellet was resuspended in 100ul of lysis buffer (2%SDS, 50mM Tris pH 8.5, 10mM TCEP, 40mM CAA, protease inhibitors (Roche) and phosphatase inhibitor (Roche)). Samples were homogenized with continuous mixing until viscosity decreased (10–20 strokes) and then boiled for 10 min at 95°C. The samples were sonicated for 2 min (1 sec on, 1 sec off, 45% amplitude) and boiled again for 5 min at 95°C. Solubilized proteins were precipitated using methanol–chloroform. Pellets were dissolved in 100 µL of digestion buffer (8 M urea, 50 mM Tris-HCl, pH 8). Protein concentration was determined using the BCA assay (sample diluted 1:10) and adjusted to 1 µg/µL. A total of 50 µg of protein was transferred to a new tube and diluted 1:6 with 50 mM Tris-HCl, pH 8.5. Protein digestion was performed by adding Lys-C (1:50 dilution from 2 µg/µL stock; 10 µL/mg protein) and trypsin (1:100 dilution from 0.5 µg/µL stock; 20 µL/mg protein) and incubating overnight at 37°C with shaking at 550 rpm. Digestion was stopped by adding TFA to a final concentration of 1%. Peptides were subsequently desalted using Sep-Pak cartridges.

### TMT labeling of protein digests (for WCP and Crude Mitochondrial proteome)

De-salted peptides were dried and resuspended in TMT-labeling buffer (200 mM EPPS pH 8.2, 20% acetonitrile). 10μg of peptides per condition were subjected to TMT labeling with 1:2.5 peptide TMT ratio (w/w) for 1 h at room temperature. The labeling reaction was quenched by addition of hydroxylamine to a final concentration of 0.5% and incubation at room temperature for 15 min. Successful TMT labeling was verified by mixing equimolar ratios of peptides and subjecting the mix to single shot LC-MS/MS analysis. Peptides were fractionated using high-pH liquid-chromatography on a micro-flow HPLC (Dionex U3000 RSLC, Thermo Scientific). Pooled and purified TMT labelled peptides were resuspended in Solvent A (5mM ammonium-bicarbonate, 5%ACN) and separated on a C18 column (XSelect CSH, 1mm × 150mm, 3.5 µm particle size; Waters) using a multistep gradient from 3-60% Solvent B (5mM ammonium-bicarbonate, 90% ACN) over 65 minutes at a flow rate of 30 µl/min. Eluting peptides were collected every 43 seconds from minute 2 for 69 minutes into a total of 96 fractions, which were cross-concatenated into 24 fractions (whole cell proteome) or 8 fractions (crude mito proteome). Pooled fractions were dried in a vacuum concentrator and resuspended in 2% ACN, 0.1% TFA for LC-MS analysis.

### MS data acquisition (for WCP and Crude Mitochondrial Proteome)

Tryptic peptides were analyzed on an Orbitrap Fusion Lumos coupled to an easy nLC 1200 (ThermoFisher Scientific) using a 25 cm long, 75µm ID fused-silica column packed in house with 1.9 µm C18 particles (Reprosil pur, Dr. Maisch), and kept at 50°C using an integrated column oven (Sonation). HPLC solvents consisted of 0.1% Formic acid in water (Buffer A) and 0.1% Formic acid, 80% acetonitrile in water (Buffer B). Assuming equal amounts in each fraction, 1 µg of peptides were eluted by a non-linear gradient from either 7% to 40% or 7% to 28% Buffer B over 90-100 minutes (WCP or crude mitochondrial proteome, respectively) followed by a step-wise increase to 90% Buffer B in 26 minutes which was held for another 9 minutes. A synchronous precursor selection (SPS) multi-notch MS3 method was used in order to minimize ratio compression as previously described^67^. Full scan MS spectra (350-1400 m/z) were acquired with a resolution of 120,000 at m/z 200, maximum injection time of 100 ms and AGC target value of 4 × 105. The most intense precursors with a charge state between 2 and 6 per full scan were selected for fragmentation (“Top Speed” with a cycle time of 1.5 seconds) and isolated with a quadrupole isolation window of 0.7 Th. MS2 scans were performed in the Ion trap (Turbo) using a maximum injection time of 50 ms, AGC target value of 15000 and fragmented using CID with a normalized collision energy (NCE) of 35%. MS3 acquisition was performed on the 10 most intense MS2 fragment ions with an isolation window of 0.7 Th (MS) and 2 m/z (MS2). Ions were fragmented using HCD with an NCE of 50% and analyzed in the Orbitrap with a resolution of 50,000 at m/z 200, scan range of 100-500 m/z, AGC target value of 150000 and a maximum injection time of 50 or 86 ms (total or crude mitochondria proteome, respectively). Repeated sequencing of already acquired precursors was limited by setting a dynamic exclusion of 60 seconds and 7 ppm and advanced peak determination was deactivated. All spectra were acquired in centroid mode.

### MS of WCP-MS data processing and statistical analysis

Raw data were analyzed using Proteome Discoverer v2.4 (Thermo Fisher Scientific) or MaxQuantv1.6.17.0 ^63,68,69^. Acquired MS2 spectra were used to screen the human reference proteome (Taxonomy ID 9606) downloaded from UniProt (17 April 2022; “One Sequence Per Gene”, 20509sequences) and a collection of common contaminants (244 entries from MaxQuant’s“contaminants.fasta”) using SequestHT, allowing a precursor mass tolerance of 7 ppm and a fragment mass tolerance of 0.5 Da after recalibration of mass errors using the Spectra RC-node^63^ ^70^. In addition to standard dynamic modifications (methionine oxidation and N-terminal loss) and static modifications (cysteine carbamidomethylation), TMT labeling of N-termini and lysine 9 side chains were set as further static modifications. The false discovery rate (FDR) was controlled using Percolator (< 1% FDR on at the level of peptide-spectrum matches). Only matches with a signal-to-noise ratio > 10 and a co-isolation < 50% as well as synchronous precursor selection (SPS) matches > 50% derived from unique peptides were used for protein quantification. For statistical analysis, contaminant proteins were removed and only proteins quantified in all replicates in all experimental groups were used for downstream analysis. Further statistical analysis was carried out using Perseus v1.6.7.0 For volcano plot analysis, significantly enriched interactors were analyzed in VolcanoR^71^. For Gene Ontology (GO) analysis, significantly enriched proteins (p<0.05) were analyzed in ShinyGO 0.80 online tool (http://bioinformatics.sdstate.edu/go). MS proteomics data were deposited in the PRIDE partner repository of the ProteomeXchange Consortium using the indicated dataset identifier^72,73^.

### MS data analysis (Crude Mitochondrial Proteome)

MS raw data were analyzed using FragPipe v22.0, with MSFragger v.4.1^74^and Philosopher v.5.1.1^75^. Acquired spectra were searched against the human reference proteome (UP0000005640) downloaded from UniProt (07/08/2024; 20468 entries) with a precursor mass tolerance of 20 ppm and fragment mass tolerance of 20 ppm. Identifications were filtered to obtain false discovery rates (FDR) below 1% for both peptide spectrum matches (minimum peptide length of 7) and proteins using a target-decoy strategy. For all searches, carbamidomethylated cysteine was set as a fixed modification and oxidation of methionine and N-terminal protein acetylation as variable modifications with allowing up to 3 modifications per peptide. Strict trypsin cleavage was set as protein digestion rule. Isobaric quantification was performed using IonQuant v.1.10.27^76^. Data were further processed using FragPipe Analyst ^77^ and plotted with custom R script.

### Lysosomal immunoprecipitation (LysoIP)

LysoIP was performed in parental HeLa cells and HeLa cells expressing TMEM192-3xHA, as described previously^78^. Six million cells were seeded per condition in three biological replicates. Cells were transfected with the GFP-CCPG1 plasmid overnight on the day of seeding. On the following day, cells were treated with either bafilomycin A1 (200 nM, 5 h), EBSS, rotenone (1 µM, 5 h), or combinations thereof, and harvested in KPBS (136 mM KCl, 10 mM KH_2_PO_4_, adjusted to pH 7.25 with KOH).

For Western blot analysis, an input fraction (2.6%) was collected prior to homogenization, lysed in 4X Laemmli buffer, and boiled for 10 min at 95 °C, 0.8% input was applied on the gel. Cells were homogenized with 25 strokes in a 10-mL tissue homogenizer and centrifuged at 1,000 × g for 2 min at 4 °C. The supernatant, containing the cellular organelles, was incubated with 20 µL of anti-HA magnetic beads on a rotator shaker for 15 min at 4 °C. The beads were washed four times with KPBS. For Western blot analysis, proteins were eluted with 2X Laemmli buffer and samples were boiled for 10 min at 95 °C.

For mass spectrometry analysis, the beads were eluted in 3% sodium deoxycholate buffer, followed by reduction with 5 mM TCEP (tris(2-carboxyethyl) phosphine) and alkylation with 20 mM CAA (chloroacetamide). Samples were then digested with trypsin for 16 h at 37 °C with shaking at 500 rpm. Digestion was stopped by adding 1% trifluoroacetic acid (TFA) in isopropanol. Peptides were cleaned up and desalted using SDB-RPS (styrene-divinylbenzene-reversed-phase sulfonate) StageTips for subsequent mass spectrometry analysis.

### Fluorescence microscopy

Cells were grown on 12-mm glass coverslips or in 96-well plates and washed with PBS prior to fixation in 4% paraformaldehyde (PFA) for 10 min at room temperature. After additional PBS washes, cells were permeabilized with 0.1% saponin in PBS and blocked with 1% FBS for 1 h at room temperature. Primary antibodies, diluted in 1% FBS/0.1% saponin in PBS, were incubated overnight at 4°C. Following three PBS washes, secondary antibodies were applied and incubated for 1 h at room temperature. Cells were then washed three more times before nuclear staining with DAPI for 10 min, followed by a final PBS wash prior to mounting with Fluoromount-G. Secondary antibodies were used at a 1:1,000 dilution: anti-rabbit Alexa 488, anti-mouse Alexa 546, and anti-rat Cy5. Images were acquired using a Leica SP8 confocal microscope equipped with a ×63 oil-immersion lens and analyzed with ImageJ software (version 1.51w). For ER-phagy flux assays, doxycycline-inducible mCherry–GFP–CCPG1 U2OS cells and RFP-GFP-KDEL U2OS cells were analyzed using a Yokogawa CQ1 (vR1.08.01) confocal imaging cytometer. Mitochondrial area and circularity, as well as SPLICs ER–mitochondria remodeling parameters (area, counts, and density), were assessed by confocal microscopy in cells either detecting endogenous TOMM20 or transiently expressing constructs in knockdown cells. Image analysis was performed using Fiji Software.

Images were acquired using a Leica STELLARIS confocal microscope with identical acquisition settings between conditions. The conditions of imaging were subsequently processed using Leica LIGHTNING adaptive deconvolution to improve spatial resolution and signal-to-noise ratio. The same deconvolution settings were applied to all images used for quantitative analysis.

### SRAI flux reporter analysis

For screening, HeLa cell lines stably expressing the indicated reporters — tagged with the SRAI (signal retaining autophagy indicator) probe at either the C- or N-terminus — were generated (see **Supplementary Table 2**), imaged, and analysed as previously described^28^. Briefly, HeLa cells were transduced with lentivirus expressing the SRAI-tagged constructs, and pools of positive cells were selected by FACS. For knockdown experiments, cell pools underwent siRNA-mediated silencing of *Ccpg1* (or a non-targeting siControl) prior to plating as previously described.

For each experiment, 50,000 cells were plated on glass-bottom 96-well plates (IBL, Cat# P96-1.5H-N). For the SRAI-CCPG1 line, doxycycline (Bio-Techne, 4090; 0.5 µg/ml, 24 h) was added to the medium at the time of plating. The following day, cells were subjected to the indicated treatments — full nutrient medium or starvation (EBSS, 6 h), rotenone (Sigma-Aldrich, R8875; 1 µM, 6 h), tunicamycin (Sigma-Aldrich, T7765; 10 µg/mL, 6 h), or haloperidol (Sigma-Aldrich, H1512, 10 µM, 6h) — before being fixed with 4% paraformaldehyde (PFA) in PBS (pH 7.2) for 10 min at room temperature.

Cells were washed twice with PBS and imaged using automated acquisition on a Nikon Ti2 inverted microscope (Nikon Instruments UK Ltd., Kingston Upon Thames, UK) equipped with a 40X NA 0.75 objective and a Nikon NIS-Elements JOBS module^79^. Widefield illumination was provided by a CoolLED PE-4000 LED light source combined with LED-CFP/YFP/mCherry-3X-A (Pinkel) filter sets, and images were captured using a Photometrics Prime BSI sCMOS camera. The TOLLES:YPet index was quantified using a custom ImageJ macro, as previously described^79^. Mean values from three to four biological replicates were presented as heatmaps generated in GraphPad.

### Immunohistochemistry of pancreas tissue and TOMM20 analysis

Pancreata from 8-weeks old *Ccpg1*^+/+^ or *Ccpg1*^GT/GT^ sibling mice on a C57BL/6 background were excised and fixed at room temperature overnight in 10 % neutral buffered formalin. They were then embedded in paraffin and standard sections cut onto glass slides for staining. Prior to processing, sections were de-waxed with xylene and rehydrated through decreasing v/v ethanol immersions. Heating in sodium citrate buffer (pH 6.0) was performed as epitope retrieval. Sections were then permeabilised in 0.1% Triton-X-100/PBS for 10 minutes and blocked for 1 hour at room temperature in 1xPBS / 5% goat serum / 0.3% Triton-X-100. Sections were then washed thrice with 0.1% Triton-X-100 in PBS for 2 min and incubated with endogenous mouse IgG blocking (0.13 mg/ml AffiniPure Fab Fragment (Jackson ImmunoResearch, 115-007-003) in PBS) for 1 h at room temperature. Primary antibodies (TOM20-V5 and REG3-V5 see table above) were diluted in protein block (Abcam, ab64226) and incubated overnight at 4 °C. After 3 × 5 min PBS washes, slides were incubated with secondary antibody (1:200 in PBS) for 1 h at RT (see table). Following PBS washes, sections were incubated with DAPI (1:1000) for 10 minutes, samples were mounted with Dako mounting media (Agilent, S3023) and a glass coverslip and sealed with nail varnish. For observation of TOM20, pancreas sections were imaged using a Zeiss FV30-HSD confocal microscopy system a 60x objective lens (Objective Lens na = 1.35, name = UPLSAPO 60, refractive index = 1.518 Lens wd = 0.15) and z drive. The z step size was 1 µm and captured each channel at each point in the stack. To count spherical TOM20 structures, all images were processed with FIJI v2.1.0 software (ImageJ, NIH). A macro was run to quantify particles with a size = 0.10-3 and circularity = 0.0-1.00 in each TOM20 image. To obtain TOM20 intensity quantifications, 3D reconstructed z-stacks were analysed using Imaris software (v10.0). In detail, Z-stacks were opened in Imaris in their native format and were automatically reconstructed into a multi-channel 3D model during input, with no further image pre-processing. Total maximum intensity values were exported for each field. Both the number of spherical structures and total intensity values were normalised to the number of cells per field. This was determined using DAPI staining for nuclear count using the Structures tool in Imaris. A one-sample t-test (theoretical mean = 1) was conducted on each set of data wherein the control mice were normalised to 1.

For the CCPG1 rescued mice experiment, the sample TOMM20 – V5 images were captured with an Olympus FV3000 Confocal Laser Scanning Microscope (Evident) using Fluoview FV31S-SW (version 2.4.1.198) Software. The microscope is composed of an IX83 inverted frame and galvanometer scanhead coupled to a laser bed containing 405, 432, 488, 515, 561 and 640 nm laser lines. Fluorescence images were captured using an Olympus UPlanSApo 60× 1.35NA Oil objective (Evident). The z step size was 1 µm and captured each channel (A405, A488, A561) at each point in the stack.

REG3-V5 images were acquired using a 40x Plan Fluor 0.75 N.A. lens on the multimodal Imaging Platform Dragonfly (Andor technologies) equipped with 405, 445, 488, 514, 561, 638 and 735nm lasers built on a Nikon Eclipse Ti2 inverted microscope body with Perfect focus system (Nikon Instruments). Data were collected in Spinning Disk 40µm pinhole mode on the Sona-6 Extreme (4.2B-6) sCMOS camera using Andor Fusion acquisition software. Z stacks were collected using the Nikon Ti2 focus drive.

### EM from cells and tissue sample preparation and image acquisition

HeLa siControl, siCCPG1 and siSIGMAR1 cells were grown in 6 well plates and treated with EBSS and Bafa1 for 6 hours. The cells were wash rapidly twice in PBS. Fixation of cells was with 1% of glutaraldehyde in 0.2M HEPES buffer for 1h (pH 7.2-7.4) and then fixed cells were washed 5 times with PBS. 1%BSA in PBS was added to the fixed cells and they were collected with the use of a scraper. Cells pellet was obtained upon centrifugation at 10.000rpm. The cell pellet was washed 5 times for 3 min with PBS without resuspending the pellet. The pellet sample was stored in PBS at 4°C.

Pancreas tissue from 8-weeks old *Ccpg1*^+/+^ or *Ccpg1*^GT/GT^ sibling mice on a C57BL/6 background were intracardiac perfusion fixed for 10 minutes with freshly-made 2% (w/v) paraformaldehyde, 2% (w/v) EM-grade glutaraldehyde in Sorenson’s Phosphate Buffer (0.1M NaH2PO_4_, 0.1M Na_2_HPO_4_, pH 7.4). The pancreas was then excised, diced into approximately 1mm^3^ pieces, and post-fixed overnight in a new volume of fixative. Samples were retained in this fixative, at room temperature, while shipping to the electron microscopy facility. Similar procedure of fixation was performed for the CCPG1 rescued mice experiment (Luciferase-V5, WT CCPG1 and 5XMutant).

Both Hela cells fixed pellets as well as small pieces of pancreas tissue were post-fixed in a mixture of 1% osmium tetroxide and potassium ferrocyanide, dehydrated in ethanol and acetone and embedded in epoxy resin as described previously 88. Thin 50-60 nm sections were cut using a Leica EM UC7 ultramicrotome. Ribbons of sections were transferred on Formvar-coated copper slot grids and enhanced contrasted with 5% uranyl acetate in methanol/water and lead citrate. EM images were acquired using a FEI Tecnai-12 electron microscope (FEI, Eindhoven, Netherlands) equipped with a VELETTA CCD digital camera (Soft Imaging Systems GmbH, Munster, Germany) or a Zeiss TEM900 microscope operated at 80 keV in the bright-field mode and equipped with a Troendle 2K camera.

### Ultrastructure analysis of TEM images

TEM images were analyzed using Fiji software. All the morphological mitochondria parameters such as Mitochondria area, contacts length, and distance from ER and mitochondria were determined by direct measurements of these structures. Mitochondria were identified based on established ultrastructural criteria, including the presence of a double membrane, characteristic mitochondrial matrix, and recognizable cristae. These criteria were applied consistently across all experimental conditions, including mitochondria displaying reduced matrix electron density or altered morphology. Mito-autolysosomal structures were identified as membrane-bound autophagic compartments containing recognizable mitochondrial material. The percentage of mitochondria associated with mito-autolysosomal structures was quantified relative to the total mitochondrial number within the analyzed fields. All the data is the average of more than 10 different fields from each condition and every membrane structure was considered to avoid bias analysis in both Hela cells and pancreatic mice tissue.

### Measurement of mitochondrial membrane potential and ROS

For live cell imaging, after 2 days of knock down for SIGMAR1 and CCPG1, mPDACs cells were seeded onto 35 mm Petri dishes with a glass bottom. Cells were incubated with Mitotracker Green and co-incubated with TMRE or mitoSOX (Invitrogen) for half an hour. Subsequently, cells were washed in phosphate-buffered saline (PBS) and normal DMEM was replaced with phenol-red-free DMEM (Thermo). Dishes were live-imaged on a 37 °C warming stage, for no longer than 5 minutes and images were collected using a Leica Microscope Confocal SP8 fitted with a 60xoil-immersion lens. Images were analyzed and processed using the ImageJ software.

### CryoET sample preparation

HCT116 cells were seeded using a modified version of a previously available protocol^80^. Briefly, cells were cultured in presence of DMEM and the respective resistances for each cell line. Briefly, EM grids (200 Mesh Au SiO2 R1/4, Quantifoil) were glow discharged on both sides for 90 s using a Pelco easiGlow device in negative polarity and 15 mA current. Then, grids were transferred to a cell culture hood and coated with 2 µg/ml of Fibronectin (Sigma-Aldrich, Merck KGaA F0895-1MG) for 1 hour at room temperature under UV light for sterilization. Then, grids were washed once with PBS and transferred into 8-well µ-slide (ibidi, Cat. No: 80826-90) containing DMEM medium. Cells were seeded on the grids at a concentration of around 30.000 cells/ml in a final DMEM volume of 300 µL. Cells were allowed to attach to the grids over-night in the incubator. Next day, the media was exchanged and cells were treated with EBSS (ThermoFisher) and Bafilomycin A (LC laboratories) for 3-4 hours prior to vitrification. Vitrification was carried out using an EM GP2 grid plunger (Leica Microsystems) at a temperature of 37°C and 70% chamber humidity. As fiducials, 4 µl of a 1:20 dilution in Dulbecco PBS Buffer of autofluorescent 1 µm diameter Dynabeads (Dynabeads MyOne carboxylic acid No. 65011, ThermoFisher Scientific) were applied to each grid. Grids were then immediately blotted from the back for 4 s followed by plunge freezing in liquid ethane at −184°C. Grids were then stored in liquid nitrogen until further use. Prior to loading, grids were clipped into Autogrid rings with two markings on the sides of the gird to aid for orientation prior to next steps.

### Plasma-FIB milling for lamella preparation

Samples were loaded into an Arctis cryo-Plasma Focus Ion Beam (cryo-PFIB) instrument (Thermo Fisher Scientific) equipped with an integrated Fluorescence Light Microscope (iFLM) for correlative lamella preparation prior to Transmission Electron Microscopy (TEM). Grids were loaded so that one of the markings was facing the top when inside the multispecimen casette holder. After loading the casette and performing an inventory, all further steps were performed using the Arctis WebUI software (versions 1.4-1.6,^81^). Xenon was used for all steps described in this protocol. A custom template was built for performing automated milling at manually selected positions. Briefly, an initial grid overview was taken to evaluate grid quality (cells, ice thickness…). Then, an electron tileset was taken at 90° from the grid using the SEM with a dwell time of 3 μs. Then, a three step “sputter-GIS-sputter” layer was applied to the front of the grids to protect the leading edge of future lamellas during milling. The sputtered platinum layer was applied by redeposition at 12 kV and 70 nA for 120 s, followed by 120 s of platinum coating using the gas injection system (GIS) loaded with trimethyl(methylcyclopentadienyl) platinum (IV), followed by another 90 s of sputter coating with platinum using the same parameters as before. Then, positions were manually placed at cells in suitable positions of the grids for automated milling. During the correlative-milling workflow, eucentric eight was set for each lamella site and milling angles between 12-16° were selected prior to the acquisition of an Ion Beam (IB) image. After this, for each position, a Z-stack was acquired with the iFLM using the green and red channels making use of the automated focus and automated exposure functions integrated within WebUI for each site. Z-stacks of 10 µm were acquired with a Z step size of 500 nm. Specifically, for SPLICS-LC3B samples, WebUI 1.6 was used with the automatic deconvolution of images which was used to increase the signal of the images prior to the correlation. For the correlation, Points of Interest (POI) were placed at the location of fluorescent puncta within cells together with fiducial markers for each lamella site (the 1 µm Dynabeads were used as fiducials as they are visible in both fluorescence and IB image). A correlation step was performed manually for each site within webUI and lamella patterns were placed at the most reliable position indicated by the correlation with the fiducial markers. At least 3 fiducials were used for each POI. Finally, each sample was milled to a final thickness of 150-180 nm. Milling was performed automatically in WebUI in a stepwise manner: first, stress-relief cuts were milled at a distance of 6 μm to each side of the intended lamella using a 1.0-nA ion beam and a depth correction factor of 2. Three milling steps were then used to remove the material above and below the intended lamella position: rough milling at 1.0 nA to 1-μm thickness (depth correction of 2), followed by a second step at 0.3 nA to thin down to 500 nm (depth-correction of 1), and a final step at 0.1 nA to thin down to 300 nm (depth-correction 0.88). Then, lamellas were automatically polished at a current of 30 pA to 150 to 180 nm. In some cases, manual milling was performed after the automated milling to (1) remove cellular debris left from the previous steps due to thicker specimens or (2) to make the lamella thinner if low-kV (3 kV) SEM images showed charging on the surface of lamellas. Finally, after lamella polishing, a final SEM imaging step was performed prior to a final iFLM Z-stack using the same parameters as prior to milling to confirm the presence of fluorescence signal within the lamella. For SPLICS-LC3B samples, images were also deconvolved after acquisition within WebUI 1.6.

### Tomogram acquisition

Tilt series were acquired using either a Titan Krios G4 (Thermo Scientific) cryo-transmission electron microscope operated at 300 kV equipped with a Selectris X energy filter and Falcon 4i direct electron detector or a Titan Krios G2 (Thermo Scientific) equipped with a Gatan BioQuantum energy filter and K3 camera. After grid transfer for the Arctis PFIB, a grid map montage was acquired at 125x magnification, followed by lamella montages acquired at 8700x magnification. Additionally, an intermediate magnification (470x) was used to acquire images of full grid squares containing milled lamella. A 2D correlation step was performed in 3D-Correlation Toolbox ^82^ to guide the selection of positions for tilt-series acquisition. Low-dose tilt series were collected using a dose symmetric scheme correcting for the lamella pre-tilt corresponding to the milling angle of each lamella using SerialEM (version 4.2)^83^ using the following parameters: for data acquired in the Falcon 4i detected, a 53000x magnification (corresponding to a pixel size of 2.414 Å), 10 eV energy slit; for the K3 camera, a 53000x magnification (corresponding to a pixel size of 2.176 Å) and 10 eV energy slit. In both cases, a 2° tilt-increment was used with a defocus range of −1.5 to −4.5 µm and ~150 e^−^/Å^2^ total dose. During the acquisitions, 10 frames were acquired for each tilt image and frame alignment was performed on-the-fly in SerialEM yielding aligned tilt-stacks at the end of each tilt series.

### Tomogram reconstruction and segmentation

Tilt series were processed by reordering tilt images and performing CTF estimation on already on-the-fly aligned tilt images. Tilt images with poor CTF estimation or bad quality due to artifacts (ice, drift…) in the field of view were removed manually. Tilt series alignment and reconstruction using AreTomo2 (version 1.0.0)^84^ and IMOD (version 4.12.62)^85^ using patch tracking and weighted back projection reconstruction at bin4 using 10 iterations of SIRT-like filtering. Membrane segmentations of cellular membranes were performed directly on fSIRT reconstructed tomograms using the MemBrain-Seg software (https://github.com/teamtomo/membrain-seg/) ^86^ manually refined using Napari^87^ and displayed using the ArtiaX plug-in (https://github.com/FrangakisLab/ArtiaX/ ^88^ in ChimeraX (version 1.9) ^89^. Vesicle measurements were performed in IMOD. For display purposes, denoising was performed on half tomograms, resulting from frame-splitting during MotionCor2^90^ frame alignment of the original frames stored during data acquisition. These even and odd tomograms were used for IsoNet2^91^ denoising using the noise-to-noise training workflow.

### Labelling of antibodies with DNA oligonucleotides

Antibodies were self-labelled with azide-modified docking strands according to a previously published protocol^92^. Unlabelled secondary antibody (AffiniPure donkey anti-mouse IgG, #715-005-150; AffiniPure donkey anti-rabbit IgG, # 711-005-152; AffiniPure donkey anti-sheep IgG, # 713-005-003; AffiniPure goat anti-rabbit IgG, # 111-005-003 were ordered from Jackson ImmunoResearch (USA) In brief, the antibody was concentrated with an Amicon spin filter (MWCO 100 kDa, Merck, Germany) when the antibody concentration was below 2 mg/mL. DBCO-sulfo-NHS ester linker (Jena Bioscience, Germany) was dissolved in dimethylformamide (DMF) (Sigma-Aldrich, Germany) and further diluted in 1x phosphate buffer saline (PBS). Linker and antibody were mixed in a 10:1 molar ratio and incubated for 90 min at 4°C gently shaking. The unbound linker was removed using Zeba desalting columns (Thermo Fisher Scientific, Germany). Azide-modified docking strands (sequences given in Table 3; Table 4) (P1 docking linker for Donkey anti-Mouse IgG; P1 docking linker for Goat anti-Rabbit IgG; P3 docking linker for Donkey anti-Sheep IgG; P5 or docking linker for Donkey anti-Rabbit IgG) and linker-antibody conjugate were incubated in a 10:1 molar ratio overnight at 4°C while slightly shaking. The exact sequences of the docking strands can be found in table 3 and 4. The next day, unbound DNA was removed using an Amicon spin filter (MWCO 100 kDa). The docking strand-labelled antibody was stored at 4°C.

**Table 3:** Sequences of docking and imager strands used for DNA-PAINT and CLSM measurements. ^92^.

| Name | Sequence | Supplier |
|---|---|---|
| P1 docking strand | 5’-azide-TTATACATCTA-3’ | Metabion, Germany |
| P3 docking strand | 5’-azide-TTTCTTCATTA-3’ | Metabion, Germany |
| P5 docking strand | 5’-azide-TTTCAATGTAT-3’ | Metabion, Germany |
| P1 imager strand Cy3B | 5’-TAGATGTAT-Cy3B-3’ | Metabion, Germany |
| P3 imager strand Cy3B | 5’-GTAATGAAGA-Cy3B-3’ | Metabion, Germany |
| P5 imager strand Cy3B | 5’-CATACATTGA-Cy3B-3’ | Eurofins, Germany |
| P1 imager strand ATTO488 | 5’-CTAGATGTAT-ATTO488-3’ | Eurofins, Germany |
| P3 imager strand ATTO655 | 5’-GTAATGAAGA-ATTO655-3’ | Eurofins, Germany |
| P5 imager strand ASOrange | 5’-ATACATTGA-ASOrange-3’ | Biomers, Germany |

**Table 4:**
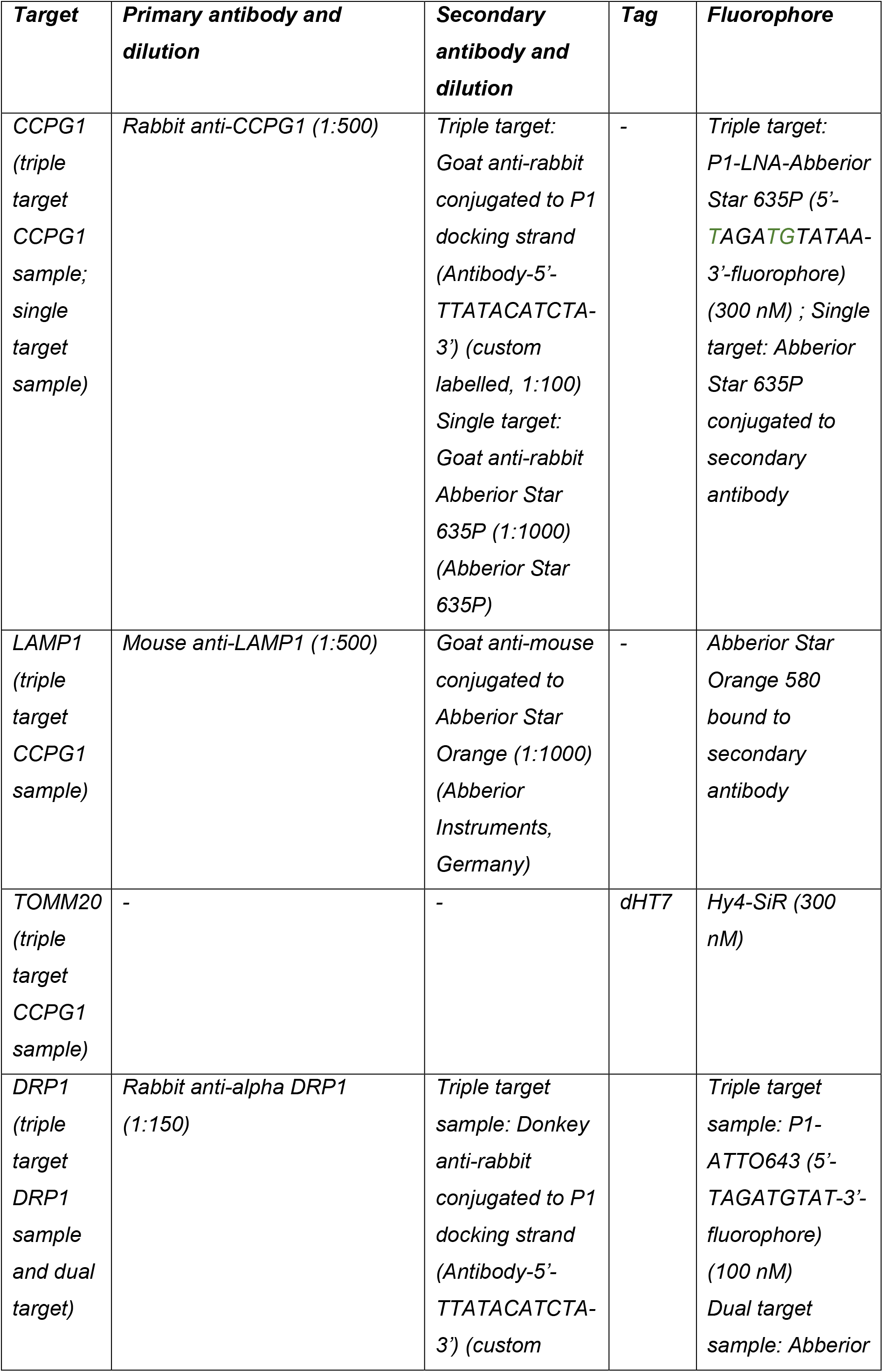
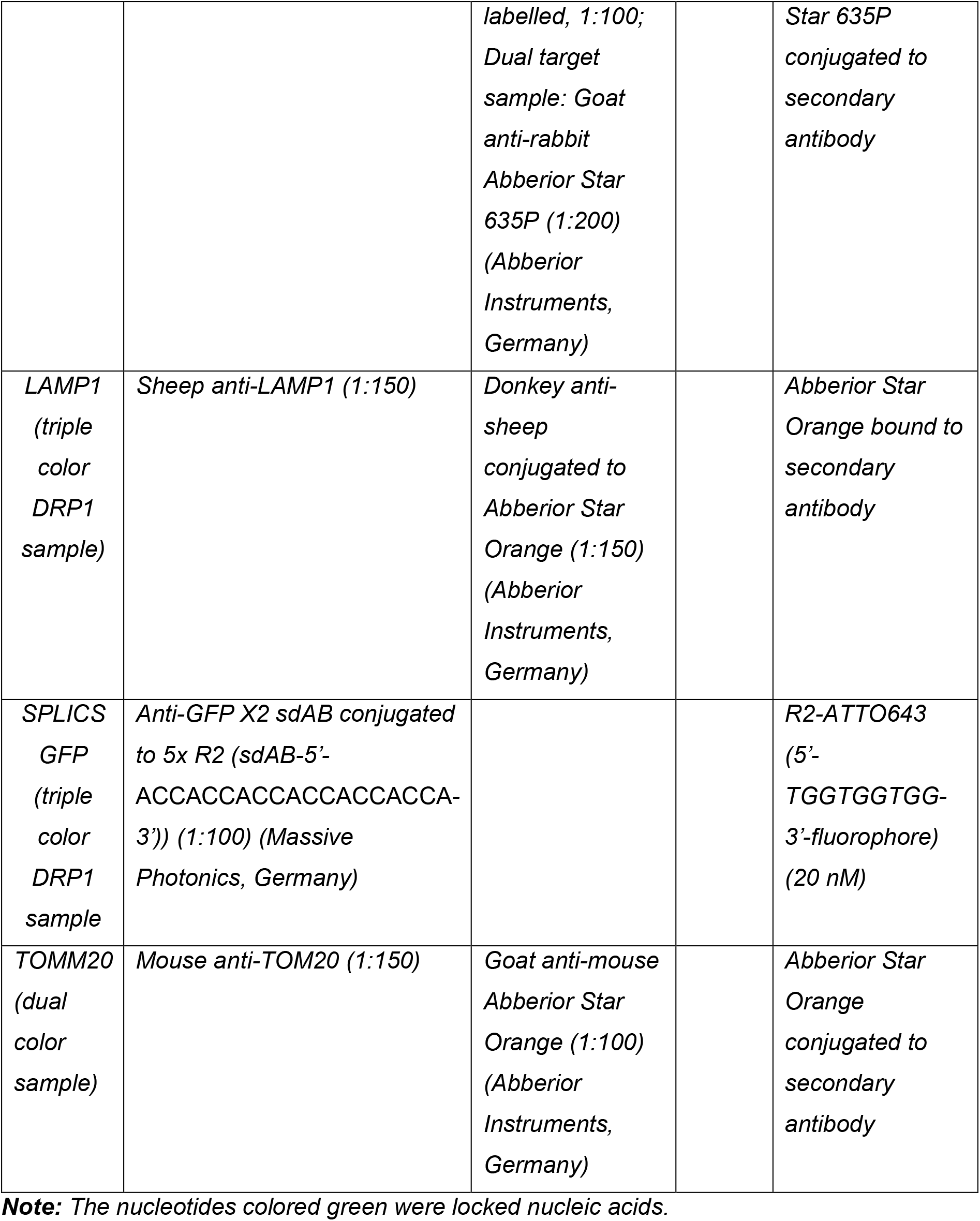
Antibodies and DNA sequences used for sample preparation, single- and multi-target STED microscopy.

| ***Target*** | ***Primary antibody and dilution*** | ***Secondary antibody and dilution*** | ***Tag*** | ***Fluorophore*** |
|---|---|---|---|---|
| *CCPG1 (triple target CCPG1 sample; single target sample)* | *Rabbit anti-CCPG1 (1:500)* | *Triple target: Goat anti-rabbit conjugated to P1 docking strand (Antibody-5’-TTATACATCTA-3’) (custom labelled, 1:100)*<br>*Single target: Goat anti-rabbit Abberior Star 635P (1:1000) (Abberior Star 635P)* | *-* | *Triple target: P1-LNA-Abberior Star 635P (5’-TAGATGTATAA-3’-fluorophore) (300 nM) ; Single target: Abberior Star 635P conjugated to secondary antibody* |
| *LAMP1 (triple target CCPG1 sample)* | *Mouse anti-LAMP1 (1:500)* | *Goat anti-mouse conjugated to Abberior Star Orange (1:1000) (Abberior Instruments, Germany)* | *-* | *Abberior Star Orange 580 bound to secondary antibody* |
| *TOMM20 (triple target CCPG1 sample)* | *-* | *-* | *dHT7* | *Hy4-SiR (300 nM)* |
| *DRP1 (triple target DRP1 sample and dual target)* | *Rabbit anti-alpha DRP1 (1:150)* | *Triple target sample: Donkey anti-rabbit conjugated to P1 docking strand (Antibody-5’-TTATACATCTA-3’) (custom labelled, 1:100; Dual target sample: Goat anti-rabbit Abberior Star 635P (1:200) (Abberior Instruments, Germany)* |  | *Triple target sample: P1-ATTO643 (5’-TAGATGTAT-3’-fluorophore) (100 nM)*<br>*Dual target sample: Abberior Star 635P conjugated to secondary antibody* |
| *LAMP1 (triple color DRP1 sample)* | *Sheep anti-LAMP1 (1:150)* | *Donkey anti-sheep conjugated to Abberior Star Orange (1:150) (Abberior Instruments, Germany)* |  | *Abberior Star Orange bound to secondary antibody* |
| *SPLICS GFP (triple color DRP1 sample* | *Anti-GFP X2 sdAB conjugated to 5x R2 (sdAB-5’-ACCACCACCACCACCACCA-3’)) (1:100) (Massive Photonics, Germany)* |  |  | *R2-ATTO643 (5’-TGGTGGTGG-3’-fluorophore) (20 nM)* |
| *TOMM20 (dual color sample)* | *Mouse anti-TOM20 (1:150)* | *Goat anti-mouse Abberior Star Orange (1:100) (Abberior Instruments, Germany)* |  | *Abberior Star Orange conjugated to secondary antibody* |

***Note:*** *The nucleotides colored green were locked nucleic acids.*

### Sample preparation for DNA exchange-PAINT

HeLa cells were cultured in DMEM/F12 media supplemented with 10% fetal bovine serum (Corning, USA), 1% (v/v) penicillin-streptomycin (Thermo Fisher Scientific, Germany) and 5% (v/v) Glutamax (Thermo Fisher Scientific, Germany). The cells were incubated at 37 °C and 5% CO_2_ and were passaged every 2-3 days or when they reached 80% confluency. Cells were seeded onto 8-well chambered coverglass (Sarstedt, Germany) at an amount of 1 × 10^4^ cells per well. The cells were incubated overnight at 37 °C and 5% CO_2_ and were stress induced the next day with EBSS and BafA1 for 6 hours. The cells were fixed with 4% formaldehyde (FA) and 0.1% glutaraldehyde (GA) for 20-30 min at 37 °C which was followed by washing three times with PBS. The fixed cells were then permeabilized using antibody incubation buffer (5% FBS in 1x PBS). After fixation and permeabilization, the cells were incubated with the respective primary antibodies (three color sample: Sheep anti-RTN4 (1:150) (AF6034, R&D systems, Germany), rabbit anti-LC3B (1:150) (MBL)/rabbit anti-LAMP1 (1:150) (abcam), mouse anti-TOMM20 (BD Biosciences); two color sample: mouse anti-LC3B (1:150) (MBL)/rabbit anti-LAMP1 (1:150) (abcam)) for 1.5 hr at room temperature (RT) while shaking. This was followed by washing thrice with 1x PBS and the cells were incubated with the respective secondary antibodies (three color sample: P3 donkey anti-sheep IgG (1:100), P5 donkey anti-rabbit IgG (1:100), P1 donkey anti-mouse IgG (1:100); two color sample: F1 sdAB anti-mouse kappa light chain (1:200) (MASSIVE-sdAB-FAST 6-PLEX, Massive Photonics, Germany), F3 sdAB anti-GFP (1:200) (MASSIVE-TAG-Q ANTI-GFP, Massive Photonics, Germany)) overnight at 4 °C. This was followed by washing thrice with 1x PBS and post fixation with 4% formaldehyde for 10 min at RT. The sample was washed thrice with PBS. This was followed by adding gold beads (Gold nanoparticles, 100 nm diameter, Product #A11-100-NPC-DIH-1-25, NanoPartz, USA) as fiducial markers were diluted 1:5 in 1x PBS and incubated for 10 min. The sample was then stored in 1x PBS for imaging.

### DNA exchange-PAINT acquisition

DNA exchange-PAINT acquisitions were performed on a home-built widefield setup based on a Nikon Eclipse Ti microscope. The excitation light was generated by a DPSS laser at 561 nm (SAPPHIRE 561-300 CW CDRH, Coherent, USA) with the required excitation power controlled by an acousto-optic tunable filter (AOTFnC-400.650-TN, AA Opto Electronic, France). A clean beam profile was achieved by coupling the laser to a fibre collimator (60FC-4-M6.2-33, Schäfter & Kirchhoff GmbH, Germany) and then into a polarisation maintaining single-mode optical fibre (PMC-E-400RGB, Schäfter & Kirchhoff GmbH, Germany) which was subsequently collimated back to a FWHM diameter of 6 mm (60FC-T-4-M50L-01, Schäfter & Kirchhoff GmbH, Germany). The collinear beam was then directed through two telescope lenses (AC255-030-A-ML and AC508-150-A-ML, Thorlabs GmbH, Germany) which focused the beam onto the back focal plane of the objective (CFI Apochromat TIRF 100XC Oil, 1.45NA, Nikon, Japan). A mirror mounted on a motorised translation stage (MTS50-Z8, Thorlabs GmbH, Germany) was used to vary the illumination angle between widefield, HILO or TIRF (HILO illumination was used in our acquisition). The excitation light was coupled into the microscope by means of a dielectric beamsplitter (zt405/488/561/640rpc, AHF Analysentechnik, Germany) which also transmitted the emission light into the detection beam path. The axial focus was maintained using an autofocus system (Ti-PFS, Nikon) and the lateral position was adjusted using a motorised stage (Ti-S-ER, Nikon) combined with a piezo stage (Nano-Drive, MadCityLabs, USA). After spectral filtering with a bandpass filter (610/60 ET, Chroma Technology Corp., USA) the emission light was projected onto an Andor Ixon Ultra EMCCD camera (DU-897U-CS0, Andor, North Ireland). The samples were excited with a laser intensity of 73.4 W/cm^2^. Exchange of buffers was made using syringes connected to the chambered coverglasses using 0.16 cm diameter FEP tubes (Part number: 1548L, IDEX Health & Science, LLC, USA) via a custom milled lid for the chambered coverglass. The injection channel and the suction channel were separate. The sample was washed thrice with 1x PBS between each target acquisition. In the case of three-color exchange PAINT imaging, each acquisition was carried out with 500 pM of the respective imager strands (round 1: P1 bound to Cy3B, round 2: P5 bound to Cy3B, round 3: P3 bound to Cy3B) diluted in 1x PBS with 0.5 M NaCl supplemented with PCA, PCD and Trolox as an oxygen scavenger system. In the case of two-color exchange PAINT imaging, each acquisition was carried out with 400 pM of the respective imager strands (round 1: F1 bound to Cy3B, round 2: F3 bound to Cy3B) diluted in 1x PBS with 0.5 M NaCl supplemented with PCA, PCD and Trolox as an oxygen scavenger system. The sequences for the imager strands are given in Table 3. Similar parameters were used for the acquisition of all targets. The parameters were: EM gain of 200; Camera exposure time of 150 ms for three color sample and 80 ms for the two color sample; Readout mode of 5000 MHz with frame transfer ‘on’. A total of 15000 frames were acquired per round.

### Image analysis of exchange-PAINT samples

All analysis of the acquired PAINT images were performed on Picasso v 0.7.0 and 0.8.0 ^92^. Each image stack acquired was first localized using Picasso Localize. In the case of three-color sample, the following parameters were used: box size 7, baseline 105 photons, sensitivity 4.32, pixel size of 158 nm a min net gradient of 45,000 for LC3B; 60,000 for LAMP1 and 70,000 for RTN4 and TOM20. In the case of the two color sample, the following parameters were used: box size 7, baseline 88 photons, sensitivity 4.75, pixel size of 158 nm a min net gradient of 32000. The localizations were drift corrected using fiducial markers and images of different channels were aligned using them. All localizations were filtered for the following parameters: sx, sy: 0.5-1-5; lpx, lpy: 0-0.2; photons: 0-6000, bg: 0-400. In the case of the three-color sample, clusters were picked with the pick tool (radius 1 pixel) in Picasso Render and only those with minimum 10 localizations spread over the complete measurement time were selected for further analysis. In the case of the two-color sample, the localizations were clustered using DBSCAN in the Picasso Render module with a radius of 75 nm and minimum 100 localizations for LC3B/LAMP1 and 30 nm and 30 localizations for SPLICS.

### CLSM of DNA-PAINT samples

Confocal imaging for the DNA-PAINT samples was performed on a Leica TCS-SP8 (Leica, Germany) equipped with a 63x oil objective (HC PL APO CS2 63x/1.40 NA), an Argon laser at 488 nm, a diode laser at 561 nm and a HeNe laser at 633 nm and photomultiplier tube (PMT) and hybrid PMT (HyD) detectors. Images were acquired at 100 Hz scan frequency with a pixel size set to 57 nm. The z-depth was set to 300 nm. Three imager strands (P1 bound to ATTO488, P5 bound to Abberior Star Orange (ASOrange), P3 bound to ATTO655) were simultaneously added to the sample at 300 nM each, diluted in 1x PBS with 0.5 M NaCl. In all cases, volumes (xyz) were acquired as frame-by-frame acquisition.

### Sample preparation for single- and multi-target STED microscopy

HeLa cells or U-2 OS cells stably expressing TOM20 conjugated to the dead mutant of HaloTag7 (dHT7) were cultured in DMEM or DMEM/F12 media (Gibco, Germany) supplemented with 10% fetal bovine serum (Corning, USA), 1% (v/v) penicillin-streptomycin (Thermo Fisher Scientific, Germany) and 5% (v/v) Glutamax (Thermo Fisher Scientific, Germany) respectively. The cells were incubated at 37 °C and 5% CO_2_ and were passaged every 2-3 days or when they reached 80% confluency. Cells were seeded onto 8-well chambered coverglass (Sarstedt, Germany) at an amount of 1 × 10^4^ cells per well. The cells were incubated overnight at 37 °C and 5% CO_2_ and induced with 100 ng/µL doxycycline the next day. After a day, the cells were stress induced with EBSS and BafA1 for 6 hours. The cells were then fixed with 4% formaldehyde (FA) and 0.1% glutaraldehyde (GA) for 20-30 min at 37 °C which was followed by washing three times with PBS. The fixed cells were then permeabilized using antibody incubation buffer (10% FBS and 0.1% Saponin in 1x PBS). After fixation and permeabilization, the cells were incubated with the respective primary antibodies (table 4) for 1.5 h at room temperature (RT) while shaking. This was followed by washing thrice with 1x PBS and the cells were incubated with the respective secondary antibodies (table 4) overnight at 4 °C. In the case of the triple target DRP1 sample, nanobody against SPLICS GFP was introduced after the secondary antibody incubation and washing with 1x PBS thrice. This was followed by washing thrice with 1x PBS and post fixation with 4% formaldehyde for 10 min at RT.

The sample was washed thrice with 1x PBS and was stored in 1x PBS for imaging. In the case of triple target DRP1 sample, after post fixation 100 nm tetraspeck beads (diluted 1:150 and sonicated for 10 min) (T7279, Thermo Fisher Scientific) were added and incubated for 15 min at RT followed by washing with 1x PBS thrice and was stored in 1x PBS for imaging.

### STED microscopy

The STED microscopy measurements were carried out on an Abberior STED Expert line microscope (Abberior Instruments, Göttingen, Germany) made of an Olympus IX83 inverted microscope (Olympus, Hamburg, Germany) with a UPLXAPO 60x NA 1.42 oil immersion objective (Olympus, Hamburg, Germany), and operated by the Imspector software (v16.3.15521; Abberior Instruments, Göttingen, Germany).

The samples used for CCPG1 cluster size quantification were measured with 5% (7.6 µW) excitation from a 640 nm and stimulated emission using a pulsed STED laser of 775 nm (2D donut point spread function) at 8% (88.5 mW). The pixel size used was 15 nm with a pixel dwell time of 5 µs, line accumulation of 15×15 nm^2^ and a pinhole size of 0.81 Airy Unit (AU). Fluorescence photons ranging between 650 nm to 755 nm were collected using an avalanche photo diode (APD) in line sequential mode with a time gate of 0.75 ns to 8.75 ns.

The samples used for triple target CCPG1 imaging used three different targeting strategies, LAMP1 was with normal immunofluorescence; CCPG1 was with LNA-PRISM^92,93^ ported to STED and TOM20 was with HaloTag-PAINT^94,95^. The images were acquired with two targets at a time in a frame sequential mode. First, LAMP1 and TOM20 were imaged followed by washing thrice with 0.01x PBS with 5 min incubation between washes and then LAMP1 and CCPG1 were imaged. LAMP1 was imaged with 8% (1.3 µW) 561 nm excitation, 7% (77.4 mW) 775 nm depletion with 10 accumulations and 5 µs dwell time. TOM20 was imaged with 5% (7.6 µW) 640 nm excitation, 6% (66.4 mW) 775 nm depletion with 6 accumulations and 5 µs dwell time. CCPG1 was imaged with 5% (7.6 µW) 640 nm excitation, 6% (66.4 mW) 775 nm depletion with 6 accumulations and 5 µs dwell time. In all three cases, the depletion laser had a 2D donut point spread function, the pinhole was set to 0.81 AU and pixel size was 15×15 nm^2^. For LAMP1 fluorescence photons ranging between 571 nm to 618 nm were collected by an APD and for both TOM20 and CCPG1, fluorescence photons ranging between 650 nm to 755 nm were collected by a second APD. In all case the time gate was set from 0.75 ns to 8.75 ns.

The samples used for triple target DRP1 imaging were acquired in two steps steps using two different targeting strategies, LAMP1 was with normal immunofluorescence, DRP1 and SPLICS were with STED-PAINT^96^. First 100 nM of P1-ATTO643 diluted in 1x PBS with 0.5 M NaCl was added to the sample and DRP1 and LAMP1 were measured together, followed by washing thrice with 0.01x PBS with 5 min incubation between washes. Next, 20 nM of R2-ATTO643 diluted in 1x PBS with 0.5 M NaCl was added to the sample and SPLICS and LAMP1 were measured together. Both imaging steps were performed as a line by line acquisition. LAMP1 was imaged with 15% (5.5 µW) 561 nm excitation, 15% (156 mW) 775 nm depletion with 20 accumulations and 10 µs dwell time. DRP1 and SPLICS were acquired with 15% (26.3 µW) 640 nm excitation, 12% (~120 mW) 775 nm depletion with 30 accumulations and 10 µs dwell time. In all three cases, the depletion laser had a 3D top hat point spread function, the pinhole was set to 0.61 AU and pixel size was 25 nm. For LAMP1 fluorescence photons ranging between 571 nm to 618 nm were collected by an APD and for both DRP1 and SPLICS, fluorescence photons ranging between 650 nm to 755 nm were collected by a second APD. In all three cases, the time gate was set from 0.75 ns to 8.75 ns.

The samples used for dual target DRP1-TOM20 imaging were acquired as line by line. DRP1 was measured with 8 % (~12 µW) 640 nm excitation, 20% (210 mW) 775 nm depletion with 5 accumulations and 10 µs dwell time. TOM20 was measured with 10 % (3.63 µW) 561 nm excitation, 25% (257 mW) 775 nm depletion with 20 accumulations and 10 µs dwell time. the depletion laser had a 3D top hat point spread function, the pinhole was set to 0.61 AU and pixel size was 25 nm. For DRP1 fluorescence photons ranging between 650 nm to 755 nm were collected by an APD and for TOM20, fluorescence photons ranging between 571 nm to 618 nm were collected by a second APD. In all three cases, the time gate was set from 0.75 ns to 8.75 ns. The power levels stated in W were measured at the back focal plane.

### Cluster size and density determination

Image analysis to determine cluster size was performed using ImageJ^97^ through custom macro. Briefly a raw image is opened and the brightness/contrast are adjusted automatically followed by background subtraction using “Math” function. The area of a given cell was selected using the “freehand” selection tool and clusters were picked using the “Find Maxima” function. The selected clusters are then fitted using the plugin “GaussFit OnSpot” (Peter Haub and Tobias Meckel) with the shape set to circle, fitmode set to Levenberg Marquard and a rectangle half size of thrice the pixel size. The results give the width and number of clusters and the area of the selection. In the case of DRP1 cluster density determination, multiple regions of interest (ROI) were chosen from the raw image and the above analysis repeated for the whole area of the ROI. The number of clusters determined per µm^2^ of the ROI was used as cluster density. Further analyses and statistics were performed on Origin (OriginLab, Massachusetts, USA) and in the case of CCPG1 clusters, only clusters ≥ 40 nm were used. Populations were tested for normality using Shapiro Wilk test and tested for significance either with one-way ANOVA if they were not normal or with two-sample t-test if they were normally distributed.

### Drift correction for triple target image

In the case of the CCPG1 triple target sample, the images were acquired with two targets (LAMP1 and TOM20) at a time (frame sequentially) followed by washing and the next two targets (LAMP1 and CCPG1). The two acquired LAMP1 images were aligned to each other, with the first LAMP1 image as reference using a custom python script based on pystackreg^98^. The CCPG1 was then shifted by the same shift in the second LAMP1 image using a custom python script. In the case of DRP1 triple target sample, images were acquired with LAMP1 and DRP1 as the first set and LAMP1 and SPLICS as the second set with washing in between and in both cases, the images were acquired sequentially line by line. All images had tetraspeck beads as fiducial markers and were manually drift corrected using the TrakEM2 tool in ImageJ.

### SIM microscopy and analysis

SIM imaging was performed on an Elyra 7 Lattice SIM2 inverted fluorescence microscope (Zeiss). The microscope is equipped with a Plan Apochromat 63×/1.4 oil immersion objective, with 405 nm (diode laser) and 488, 561 and 642 nm lasers, all maintained under temperature control at 37 °C with 5% CO2. Fluorescence emission was collected on two sCMOS cameras with DuoLink camera adapter. Z-stacks were acquired over a range of approximately 2 μm, with z-resolution determined by the software algorithm to achieve maximum resolution with a minimal number of slices (Leap mode). Image channels were aligned using a transformation matrix generated from a calibration measurement with fluorescent beads. The raw image data were processed using the Zeiss SIM2 processing algorithm, which were then adjusted for brightness and contrast using Fiji/ImageJ.

For 3D-reconstruction of the four-color SIM Microscopy, z-stacks were imported to Imaris (version 10.1) and rendered in 3D view for visualization.

Percentage of DRP1 particles were calculated using a custom macro for the batch of images where multiple ROIs of 200×200 pixels were marked in the field of view and the channels for DRP1, ER and mitochondria were thresholded for signal by Otsu method. To determine the percentage of DRP1 puncta associated with the ER, with mitochondria, or with both organelles simultaneously, binary masks were generated for all three channels. The total number of DRP1 puncta (N_total_) was determined, and puncta overlapping the ER mask (N_ER_), the mitochondrial mask (N_mito_), and both masks simultaneously (N_both_) were counted. The graphs were plotted for percentage of DRP1 overlap with ER, mitochondria and both.

The percentage of DRP1 overlapping with SPLICS was measured using a custom macro where freehand ROIs were drawn around individual cell boundaries in the field of view and percentage of DRP1 with SPLICS was calculated. Also, Pearson’s correlation between DRP1 and SPLICS was calculated and plotted.

### Subcellular fractionation to obtain ERMCSs (MAMs)

Cellular fractionation was performed as previously described in ^27^. In brief, cells were trypsinized and pelleted. The cell pellet was resuspended in IB_cells_1 buffer (225 mM mannitol, 75 mM sucrose, 0.1 mM EGTA and 30 mM Tris-HCl, pH 7.4) at 4 °C. Cells were homogenized using a Teflon pestle and centrifuged at 700 g for 5 minutes at 4 °C. The pellet is discarded and the supernatant is centrifuged again at 7000 g for 10 minutes at 4 °C obtaining the cytosolic fraction in the supernatant, which will be further processed for isolation of the ER. The obtained pellet is resuspended in IB_cells_2 buffer (225 mM mannitol, 75 mM sucrose and 30 mM Tris-HCl, pH 7.4) and centrifuged at 7000 g for 10 minutes, at 4 °C. The obtained pellet is the crude mitochondrial faction, which was further processed in MRB buffer (250mm of manitol, 5 mM of HEPES and 0,5mM of EGTA) and Percoll medium (250mm of manitol, 25mM of HEPES and 1mM of EGTA and Percoll) to separate MAMs and pure Mitochondria after centrifugation at 100000 g for 30 min, at 4 °C. The supernatant of the cytosolic fraction is further processed to sub-fractionate the ER by centrifugation at 20000 g for 30 minutes, at 4 °C. The resulting pellet is discarded and the supernatant is further centrifuged at 100000 g for 1 hour, at 4 °C, obtaining the ER fraction in the pellet and the cytosolic fraction in the supernatant. All the samples were further processed for SDS-PAGE analysis.

### Proximity ligation assay (PLA)

The PLA Duolink® In Situ Detection Reagents (DUO92014, Sigma-Aldrich), Duolink® In Situ PLA® Probe Anti-Rabbit PLUS (DUO92002, Sigma-Aldrich) and Duolink® In Situ PLA® Probe Anti-Mouse MINUS (DUO92004, Sigma-Aldrich) were used and experiments performed according to the manufacturer’s protocol. The primary and secondary antibodies were each used for 1 hour at room temperature at a dilution of 1:100 in antibody solution (0.1 % Saponin (47036; Sigma-Aldrich), 5 mM MgCl_2_, 10 % FBS in PBS). The secondary antibody solution also contained Alexa Phalloidin 647 and DAPI.

### qRT-PCR

For the confirmation of CCPG1 knockdown in mPDACs, cells were seeded on 6-well dishes and treated as indicated. RNA isolation and DNAse treatment were performed according to the kit’s protocol (Qiagen #74104, #79254). RNA concentration was measured, and 500 ng was used for cDNA synthesis. For cDNA synthesis, the BioRad Kit was used (#170-8891). For the qRT-PCR, 1 µL of cDNA was used. Per reaction: 1 µL cDNA, 5 µL of SYBR green mix (Roche, #04707516001), 0.5 µL of forward primer (from 10 mM stock), 0.5 µL of reverse primer (from 10 mM stock), 3 µL dH_2_O. Each condition has a technical triplicate.

Primers used are:

- GAPDH Fw: GACTATAACCCTGGCTCTATG
- GAPDH Rv: GAATGGACTGTGGTCATTAG
- CCPG1 Fw: TCCAGTAGCGATGACACCAG
- CCPG1 Rw: CTGGATTGTGCCATAGAAATGTC

### Statistical analysis

All experiments have a minimum of three biological replicates. Data are presented as the mean with error bars indicating the s.d. (standard deviation). The statistical significance of differences between experimental groups was assessed with Student’s T-test. Differences in means were considered significant if p<0.05. Differences with p<0.05 are annotated as *, p<0.01 are annotated as ** and p<0.001 are annotated as ***. Statistical values measurements were performed using Origin software with corresponding R packages, Excel or PRISM.

### Data availability

All data is available in the main manuscript text or the supplementary materials and will be made available upon request. The MS proteomics data have been deposited to the ProteomeXchange Consortium^99^ via the PRIDE partner repository^100^ with the dataset identifiers PXD051844 (Fig. 1a, b), PXD051977 (Fig. 2 k, Extended Data Figure 2l-n), PXD051843 (Extended Data Figure 3a-c), (Extended Data Figure 3g-h-i), PXD071006, PXD083749 (Figure 3e) and PXD083725 (Figure 4a-d, Extended Data Fig. 7a-b).

## Author contributions

The present study was conceived by ID, AC-P, and AG with additional input and feedback from SW. The experiments and analysis were performed by AC-P and AG. AC-P and AG prepared all the constructs. AG and AC-P carried out all the cell biology and biochemical experiments, generated all the inducible cell lines and samples for MS experiments, and analyzed the results, including MS data, with the support of RK, RR, IYAO, BM, YF, UGM, MK, and TM. NJ-M for performing the SRAI assays, together with CSC and MD, performed the mouse experiments under the supervision of SW. SAP-C, GH, and RMB performed structural modeling, simulations, and data analysis. A.B. carried out the super-resolution experiments and analysis under the supervision of MH. PV performed SIM imaging experiments and analysis under the supervision of AJG-S. The electron correlative microscopy analysis was performed by JL under the supervision of FW. Electron Microscopy experiments were analyzed by EP, with additional analysis made by MB and SE. ACP and AG wrote the manuscript with contributions from all authors.

## Acknowledgments

We thank Daniela Höller and Anne Claire-Jacomin for initial critical reading and revision of the manuscript. We thank all members of the Quantitative Proteomics Unit at IBC2 (Goethe University, Frankfurt), in particular Georg Tascher for support and expertise in Proteomics methodology and data analysis, Martin Adrian-Allgood for technical help and measurements, Kristina Wagner for preparing LC columns and David Krause for help in (bio)informatics. We thank the Central Electron Microscopy Facility at the Max Planck Institute of Biophysics, including Sonja Welsch and her team for constant technical support during sample preparation and cryo-ET data acquisition. We thank the Imaging Facility at the Max Planck Institute of Biophysics and Dr. Andreas Jenner from the facility for his technical support on SIM imaging. We thank Michael Rennie, Matthew Pearson and Morwenna Muir from the Flow Cytometry Facility, Advanced Imaging Resource and Bioresearch and Veterinary Services facilities, respectively, at the Institute of Genetics and Cancer, University of Edinburgh, for their technical support. We thank the Deutsche Forschungsgemeinschaft (German Research Foundation, DFG) for funding the LC-MS system (easy nLC 1200, QExactive HF) used in this study (Project-ID: 259130777, SFB1177 – Selective Autophagy), for funding the LC-MS system (easy nLC1200, Orbitrap Fusion LUMOS) used in this study (FuGG Project-ID: 403765277). We thank the European Fonds for regional development (REACT-EU, IWB-EFRE-program Hessen, 20008763) for the VanquishNeo, Orbitrap Ascend LC-MS systems used in this study and for INST 161/1195-1 FUGG and co-funded by the by the European Union / EFRE, Hessen financing the STELARIS confocal microscope (FLIM Module) from Leica. SAP-C is supported by the cluster project ENABLE, funded by the Hessian Ministry for Science and arts. The work from IYAO was supported by the São Paulo Research Foundation (FAPESP), grant #2025/03598-1. We thank the Deutsche Forschungsgemeinschaft (German Research Foundation, DFG) for funding the INST 161/1020-1 FUGG for financing the Abberior STED expert line microscope and the CRC project on selective autophagy DFG Project-ID 259130777-SFB1177 to I.D., G.H., M.H. and R.M.B. G.H. thanks the Max Planck Society for support. SAP-C and R.M.B. thank the Center for Supercomputing, Goethe University Frankfurt, for computing time on the Goethe-NHR cluster. Additionally, this work was supported by a Spinning disk confocal microscope MRC funded (project MC_PC_MR/X012212/1), UKRI project grant (BB/N000315/1) and a Cancer Research UK Senior Fellowship (29576) to S.W.

## Supplementary figures legends

**Extended data Figure 1.**
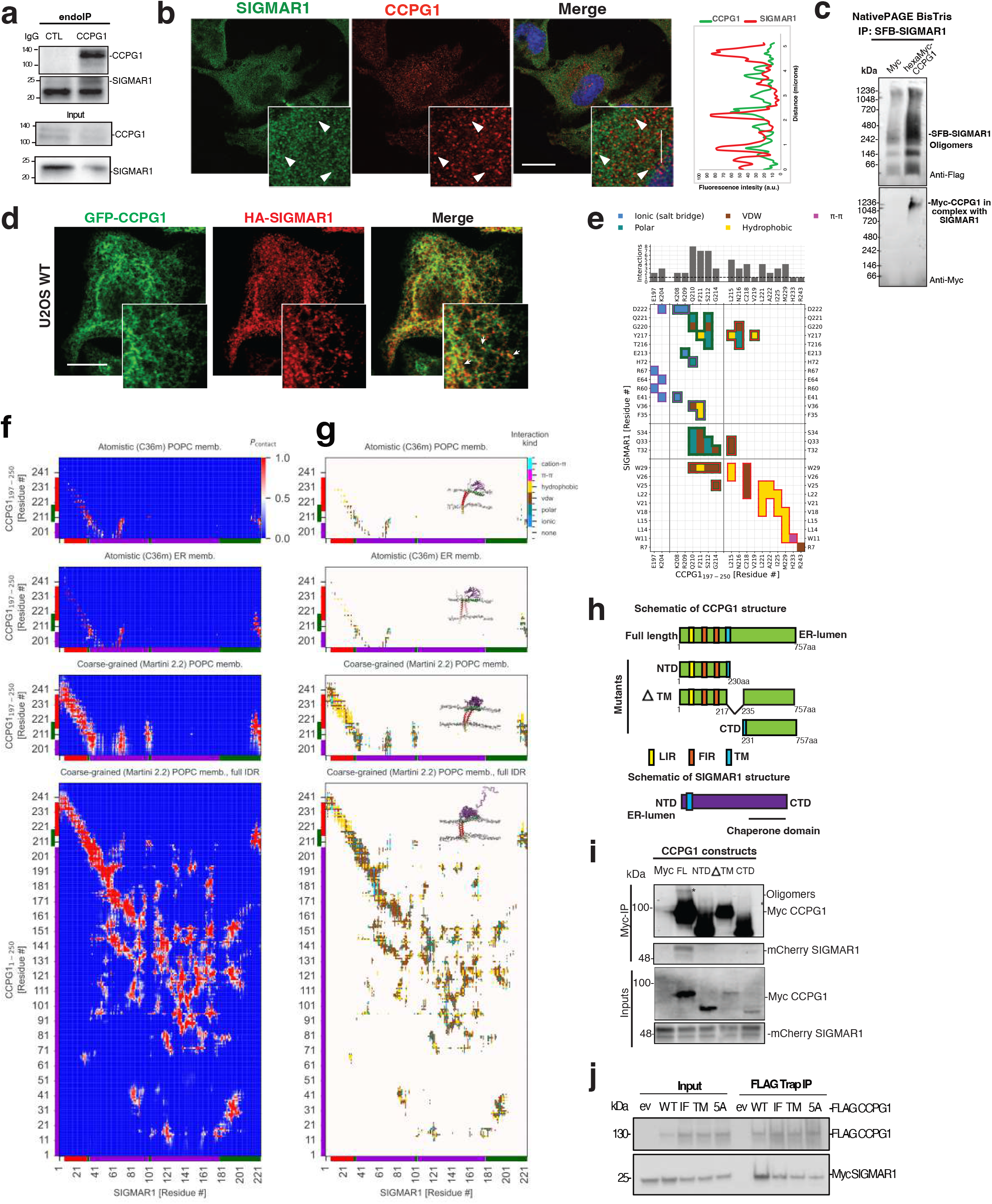
SIGMAR1–CCPG1_197-250_ complex is stabilized predominantly by interactions in the hydrophobic core of a POPC lipid bilayer. **a)** Native-PAGE analysis of SIGMAR1-CCPG1 in complex and oligomerization state of co-immunoprecipitation and eluted SFB-SIGMAR1 in complex with myc-CCPG1. **b)** Representative immunofluorescence analysis of CCPG1 and SIGMAR1 co-stained in Hela cells. On the right, Histogram analysis of fluorescence intensity distribution of both signals indicating co-localization. Arrowheads show CCPG1 and SIGMAR1 puncta colocalization. **c)** SDS-PAGE and Western blot of endogenous co-immunoprecipitation of CCPG1 and SIGMAR1. **d)** Confocal fluorescence microscopy analysis of HA-SIGMAR1 (red) and GFP-CCPG1 (green). Scale bar, 10 µm. SIGMAR1 and CCPG1 signals are distributed along the ER network. Positive HA SIGMAR1^+^ and GFP-CCPG1^+^ punctate structures are indicated with arrows (inset boxes). **e)** Heatmap of the pairwise residue contact map, obtained by averaging intermolecular contacts from all-atom MD trajectories. The plot highlights residue interactions (colored by interaction type) with high probability (*P*_contact_ > 0.05), obtained from all-atom MD simulations of PC-rich (1 µs, *n* = 2) and ER membranes (1 µs, *n* = 2). The number of interactions mediated by CCPG1-interface residues is also plotted above. **f)** Heatmaps of pair-wise residue contact map from four different MD simulations: (from top to bottom) all-atom MD simulations of CCPG1_197-250_-SIGMAR1 in POPC membrane (1 µs, *n* = 2), all-atom MD simulations of CCPG1_197-250_-SIGMAR1 in ER membrane, (1 µs, *n* = 2), coarse-grained MD simulations of the same protein-constructs (CCPG1_197-250_-SIGMAR1; 10 µs, *n* = 3), and coarse-grained MD simulations of CCPG1_1-250_-SIGMAR1 including the IDR (10 µs, *n* = 3). Three main PPI interaction zones are highlighted along the CCPG1 and SIGMAR1 protein axes. **g)** Filtered heatmaps from f highlighting the residue interaction type with *P*_contact_ > 0.5. Insets show representative snapshots of the complex structure. **h)** Schematic of CCPG1 structure and its deleterious mutants: NTD_1-230_ amino acids (N-terminal and transmembrane domain, ΔTM), Delta TM_217-235_ amino acids, and C-terminus domain: CTD_231-757_ amino acids. **i)** SDS-PAGE and Western blot analysis of co-immunoprecipitation of myc-CCPG1 mutants and mCherry-SIGMAR1. **j)** SDS-PAGE and Western blot analysis of co-immunoprecipitated CCPG1 WT, IF (interfacial), TM (transmembrane), and 5A (IF+TM) residues to alanine mutants and their binding with Myc-SIGMAR1.

**Extended data Figure 2.**
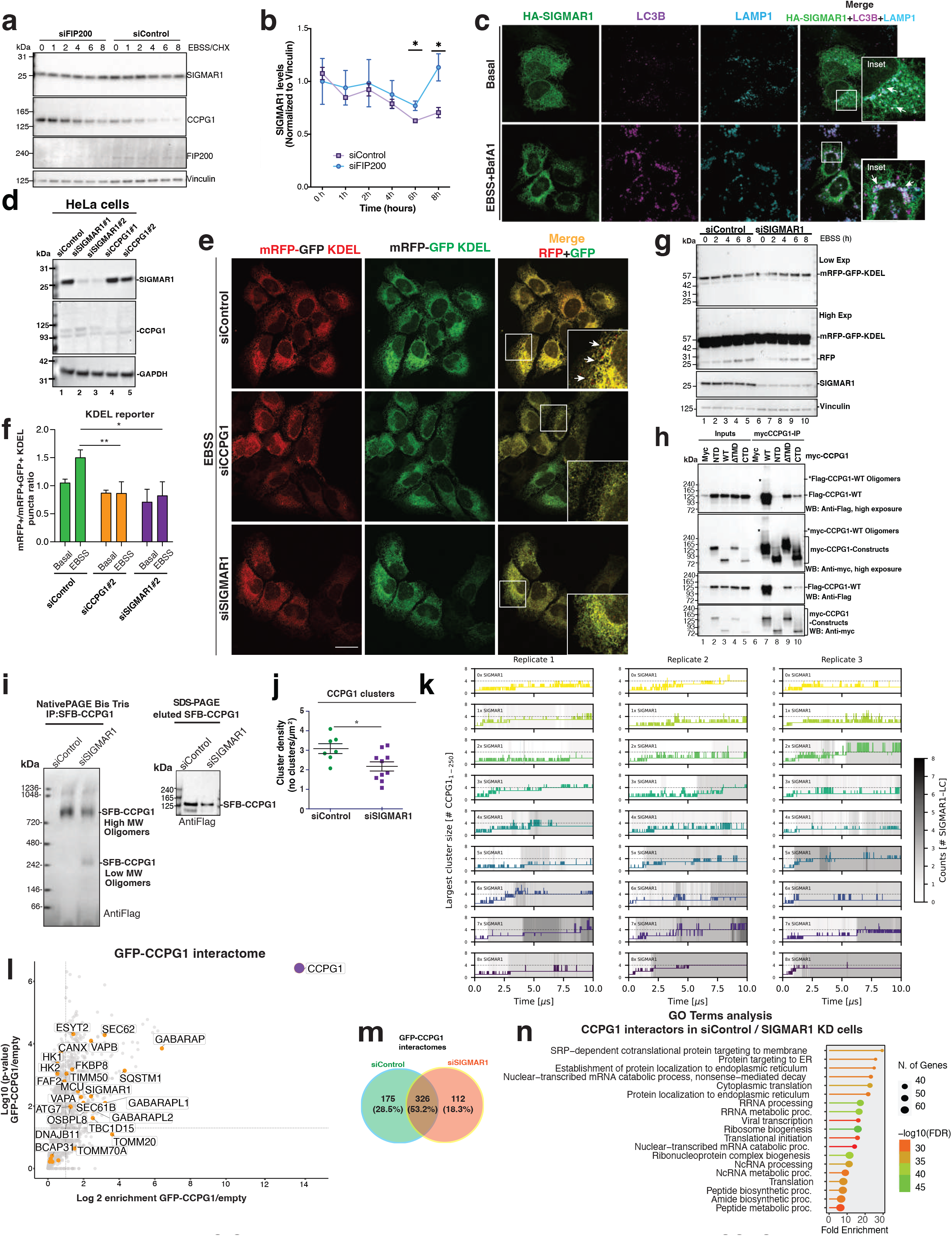
SIGMAR1 is substrate of autophagy and contributes to CCPG1-driven ER-phagy. **a)** Western blot analysis of SIGMAR1 and CCPG1 turnover in siControl and siFIP200 knock down cells for 72h under starvations conditions. **b)** densitometric quantification of the endogenous signal of SIGMAR1 in **a)** (data represent the mean ± S.D. n=2 independent experiments; two-way ANOVA, Bonferroni post-hoc test). **c)** Immunofluorescence analysis of U2OS Trex cells expressing HA-SIGMAR1 (green) and its localization with LC3B+ (red) and LAMP1+ (magenta) structures under basal and EBSS plus BafA1 treatment for 6h. **d)** Representative western blot analysis of knock down efficiency of siControl, siSIGMAR1 #1, siSIGMAR1 #2, siCCPG1 #1 and siCCPG1 #2 in Hela cells. **e)** Representative images of the ER-phagy reporter mRFP-GFP KDEL in HeLa cells transfected with siControl, siCCPG1 and siSIGMAR1.The double positive signal is localized in the ER and red only puncta within the lysosomes (cells were treated for 6 h with EBSS). **f)** ER-phagy flux was quantified as the ratio between mCherry^+^GFP^−^ and mCherry^+^GFP^+^puncta. n=3 independent experiments. Total number of cells per condition siControl (basal:8; EBSS:8); siCCPG1 (basal:6; EBSS:10) and siSIGMAR1 (basal:5; EBSS:7) for KDEL reporter which was analyzed by confocal microscopy analysis. **g)** Western blot analysis of the KDEL reporter in siSIGMAR1 KD cells. **h)** Co-immunoprecipitation assay of CCPG1 oligomers. Cells co-expressed FLAG-CCPG1 and the myc-CCPG1 WT and the mutant CCPG1 versions NTD (1-230 amino acids), delta TMD (deletion of 217-235) and CTD (231-757 amino acids). The Myc-trap IP was analyzed by western blot. **i)** Native-PAGE analysis of CCPG1 oligomers in siControl and SIGMAR1 deficient cells. SDS-PAGE of total lysates expressing SBP-CCPG1 **j)** CCPG1 cluster density (cluster number / cell area in µm^2^) of siControl (green) and siSIGMAR1 (magenta) cells. (n = 7 for siControl; n = 10 for siSIGMAR1). **k)** SIGMAR1 promotes CCPG1_1-250_ clustering. The size of the largest CCPG1_1-250_ cluster, defined by the increasing number of CCPG1 molecules, is plotted as a function of time for all 3 replicates. Top-to-bottom plots show increasing numbers of SIGMAR1 molecules in the simulations, from 0 to 8. The shade of grey represents the number of SIGMAR1 molecules co-clustered with the largest CCPG1_1-250_ cluster. Note that increased grey intensity often precedes an increase in the size of the largest cluster, indicating that SIGMAR1 molecules mediate enhanced association of CCPG1 molecules, thereby increasing cluster size. **l)** Volcano plot of significant interaction partners of GFP-CCPG1 in siControl. Data were analyzed with MaxQuant and Perseus; n = 3. Interaction partners with log_2_ enrichment > 1.0 and –log_10_ *P* > 1.3 were plotted and compared (one-sided unpaired Student’s *t*-test). The bait CCPG1 is marked in green, and the most significant interaction partner besides autophagy adaptors (GABARAPs, marked in red), ERMCS proteins (SIGMAR1, VAPs and RMDN3 in blue). All other proteins identified as significant interaction partners are represented by gray dots. **m)** Venn diagram of interactors of GFP-CCPG1 compared in presence or absence of SIGMAR1. Numbers represent significantly enriched interaction partners (log_2_enrichment > 1.0 and –log_10_ p > 1.3, one-tailed unpaired Student’s test). *n* = 3 independent experiments. **n)** Biological process pathway analysis of downregulated interaction partners of GFP-CCPG1 in SIGMAR1 KD cells (fold enrichment). The FDR of significant hits compared from siControl and siSIGMAR1 interaction partners of CCPG1 (log_2_enrichment > 1.0 and –log_10_ > 1.3, one-tailed unpaired Student’s test). *n* = 3 independent experiments.

**Extended data Figure 3.**
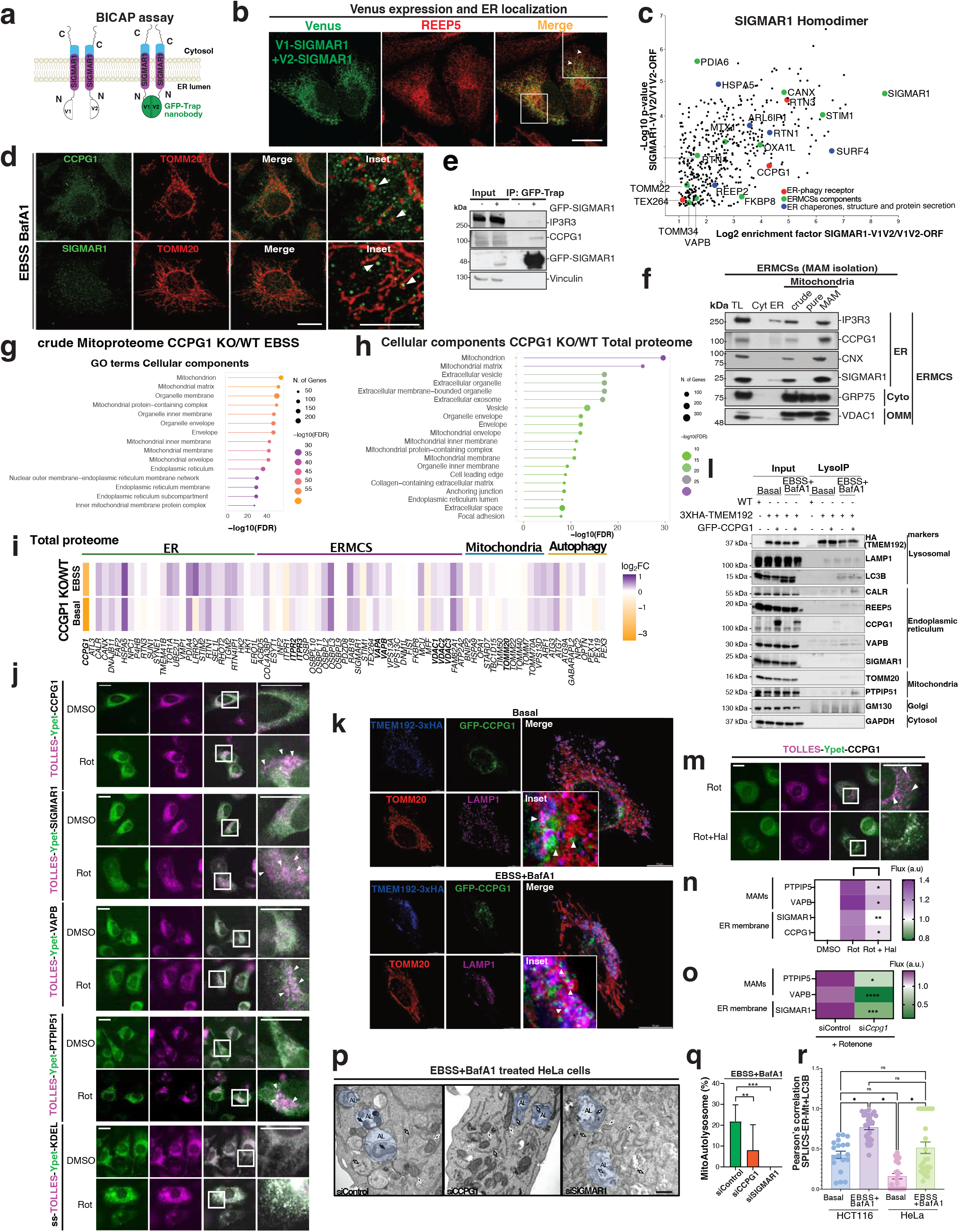
ERMCS degradation depends on CCPG1-SIGMAR1 complex. **a)** Schematic representation of the bimolecular complementation affinity purification assay (BICAP) of SIGMAR1 dimers. Top diagram, the full-length SIGMAR1 was fused to the N-terminal of the two non-fluorescent complementary fragments of the Venus fluorescent protein (V1-SIGMAR1 and V2-SIGMAR1). **b)** Immunofluorescence analysis of V1-V2 SIGMAR1 interactions and localization at the ER (REEP5+). Scale bar, 10 µm. **c)** Volcano plot showing proteins enriched in Venus-SIGMAR1 homodimers, highlighting ERMCSs components (green), ER-phagy receptors (red) and ER chaperones; structure and proteins secretion (blue). Data represent three independent experiments, one-sided unpaired Student’s t-test. **d)** Immunofluorescence analysis of endogenous CCPG1 and SIGMAR1 localized close proximity to mitochondria (TOMM20+), arrowheads in insets show the co-localization. Scale bar, 10 µm. **e)** Co-immunoprecipitation assay of GFP-SIGMAR1 interaction with CCPG1 and IP3R3, analyzed by western blot. **f)** Subcellular membrane fractionation to enrich ERMCS contacts determined CCPG1 and SIGMAR1 localization there. IP3R3, CNX, GRP75 and VDAC1 were included as controls (ER, cytosol and mitochondria, respectively). **g)** Pathway analysis of crude mitochondria proteome of CCPG1 KO compared to WT cells under starvation. Cellular component pathways analysis showed an accumulation in mitochondrion components and endoplasmic reticulum. **h)** Cellular component analysis comparing WT and CCPG1 KO cells depicting the changes in mitochondrion and endoplasmic reticulum in deficient cells. The FDR of significant hits compared from CCPG1 KO versus WT cells (log_2_enrichment > 1.0 and – log_10_ > 1.3, single-tailed unpaired Student’s test). *n* = 3 independent experiments. **i)** Heatmap showing differentially abundant proteins in the total proteome of CCPG1 KO versus control cells under starvation conditions. Data were analyzed using FragPipe and visualized in R (n = 3). Upregulated proteins associated with ER–mitochondria contact sites are highlighted, including ER proteins (CANX, SIGMAR1, HK1, CHCHD3, SEL1L, and ATP2A2) and mitochondrial proteins (TOMM20, TOMM40, TOMM70, VDAC1, VDAC2, and VDAC3). **j)** Widefield fluorescence images (TOLLES, magenta; YPet, green) of HeLa cells stably-expressing indicated proteins tagged with SRAI (n=3, representative images shown). Arrowheads indicate TOLLES-only foci (proteins that have undergone flux). Scale bars,20μm. **k)** Representative LIGHTNING-deconvolved images showing Lyso-IP expressing cells TMEM92-3XHA (blue) and GFP-CCPG1 (gree) and colocalization with TOMM20 (red) and LAMP1 (magenta) in basal and EBSS+BafA1 (4h). Scale bar 10 µm. **l)** Western blot analysis of Lyso-IP enrichment in cells expressing GFP-CCPG1 compared to non-expressing cells in Basal and EBSS+BafA1 conditions. Lysosomal (HA-TMEM192, LAMP1), autophagosome (LC3B) Endoplasmic reticulum (Calreticulin, REEP5, CCPG1, VAPB and SIGMAR1), Mitochondria (TOMM20 and PTPIP51) and Golgi-GM130 and Cytosol-GAPDH markers as control as well validated substrates of lysosomal degradation. **m)** Widefield fluorescence images (TOLLES, magenta; YPet, green) of HeLa cells stably-expressing indicated proteins tagged with SRAI (quantification in N), as in A (n = 4, representative images shown). Arrowheads indicate TOLLES-only foci (proteins that have undergone flux). Scale bars, 20 μm. **n)** Heatmap of fluorescence ratios (flux indices, see Methods) of HeLa cells stably-expressing SRAI fusions with the indicated proteins under DMSO or Rotenone (Rot, 1 μM) +/− Haloperidol (Hal, 10 μM) for 6 h (n = 4 independent replicates, ± s.e.m., 6 fields of 20-100 cells per replicate, normalised to DMSO, 2-tailed student t-test for Rot vs Rot+Hal, ** = p < 0.01, * = p ≤ _0.05). **o)** Heatmap of fluorescence ratios (flux indices, see Methods) of HeLa cells stably-expressing SRAI fusions with the indicated proteins transfected with si*Ccpg1* (or siControl) after DMSO or Rotenone (Rot, 1 μM) for 6 h (n = 3 independent replicates, ± s.e.m., 6 fields of 20-100 cells per replicate, normalised to siControl DMSO, 2-tailed student t-test for siControl vs si*Ccpg1*, **** = p < 0.0001, *** = p < 0.001, * = p ≤ _0.05). **p)** Representative TEM images of Hela WT (siControl), siCCPG1 and siSIGMAR1 cells treated with EBSS plus BafA1 for 6h. AL-autolysosomes depicted by false blue color, ER and mitochondria are indicated by white and black arrows respectively. Scale bar 500 nm. **q)** Quantification analysis of the mitochondria localized in autolysosomes from siControl, siCCPG1, siSIGMAR1 cells under EBSS plus BafA1. **r)** Pearsons’ colocalization analysis of Contact-phagy SPLICS-ER-Mt with LC3B+ structures in Hela and HCT116 cells (number of cells basal *n* = 3; ebss bafa1 n=4), HCT116 Basal n=3 and n=4). Data mean ± s.e.m. from one independent experiment; Kruskal– Wallis/Dunn’s post-test.

**Extended data Figure 4.**
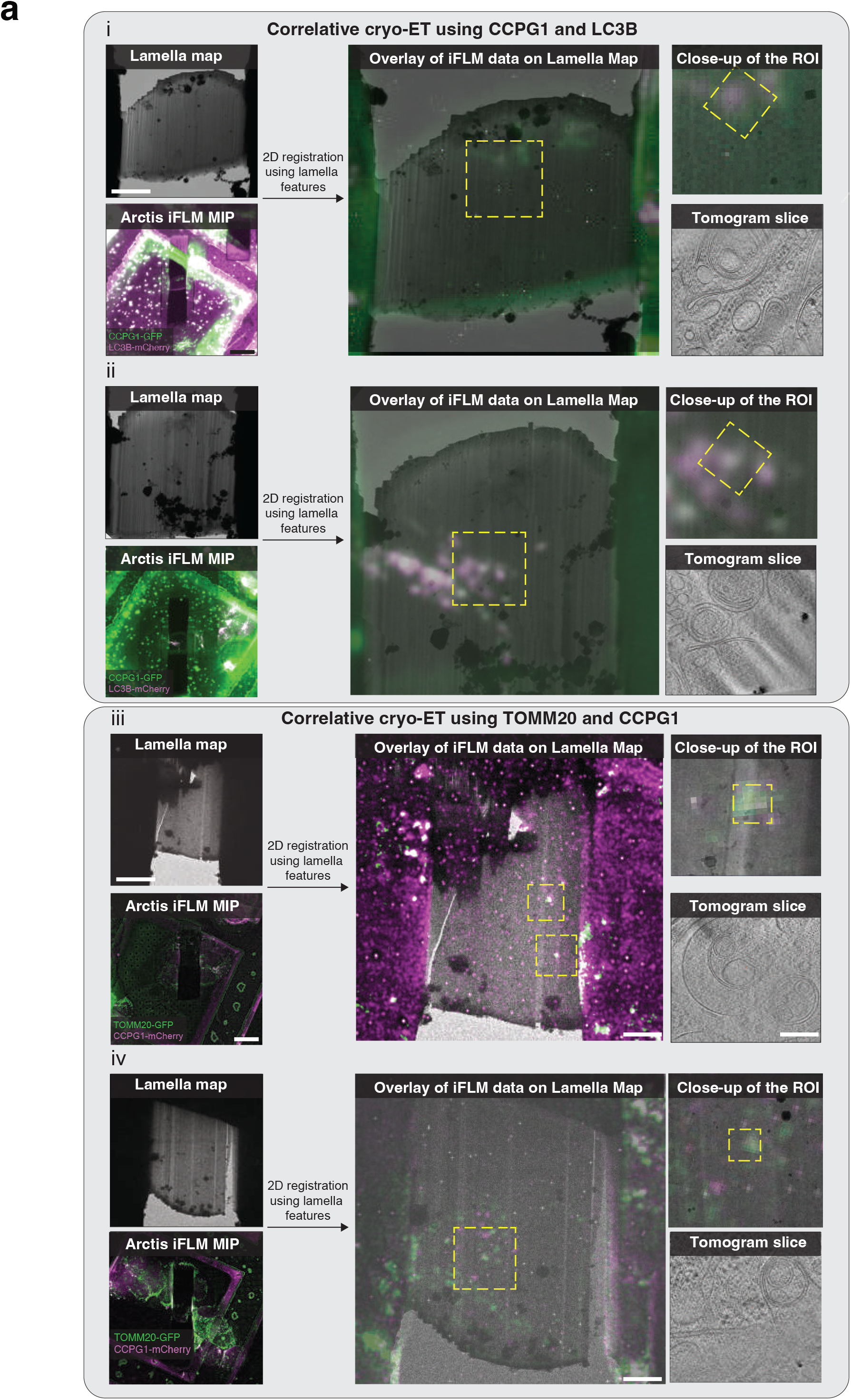
Correlative cryo-ET images of HCT116 cells labelled with contact-phagy markers. **a)** Images acquired for the correlation of GFP-CCPG1 and mCherry-LC3B (i and ii) and TOMM20-GFP and mCherry-CCPG1 (iii and iv). For each correlation, a medium magnification stitched lamella map acquired at the TEM is shown above the final iFLM stack showing on lamella fluorescence. The middle image shows an overlay of the lamella map with fluorescence. On the right-hand side, a close-up image around the region of interest (ROI) is shown with the fluorescence overlay and finally a slice of the tomogram reconstructed from the tilt series at that position is shown below.

**Extended data Figure 5.**
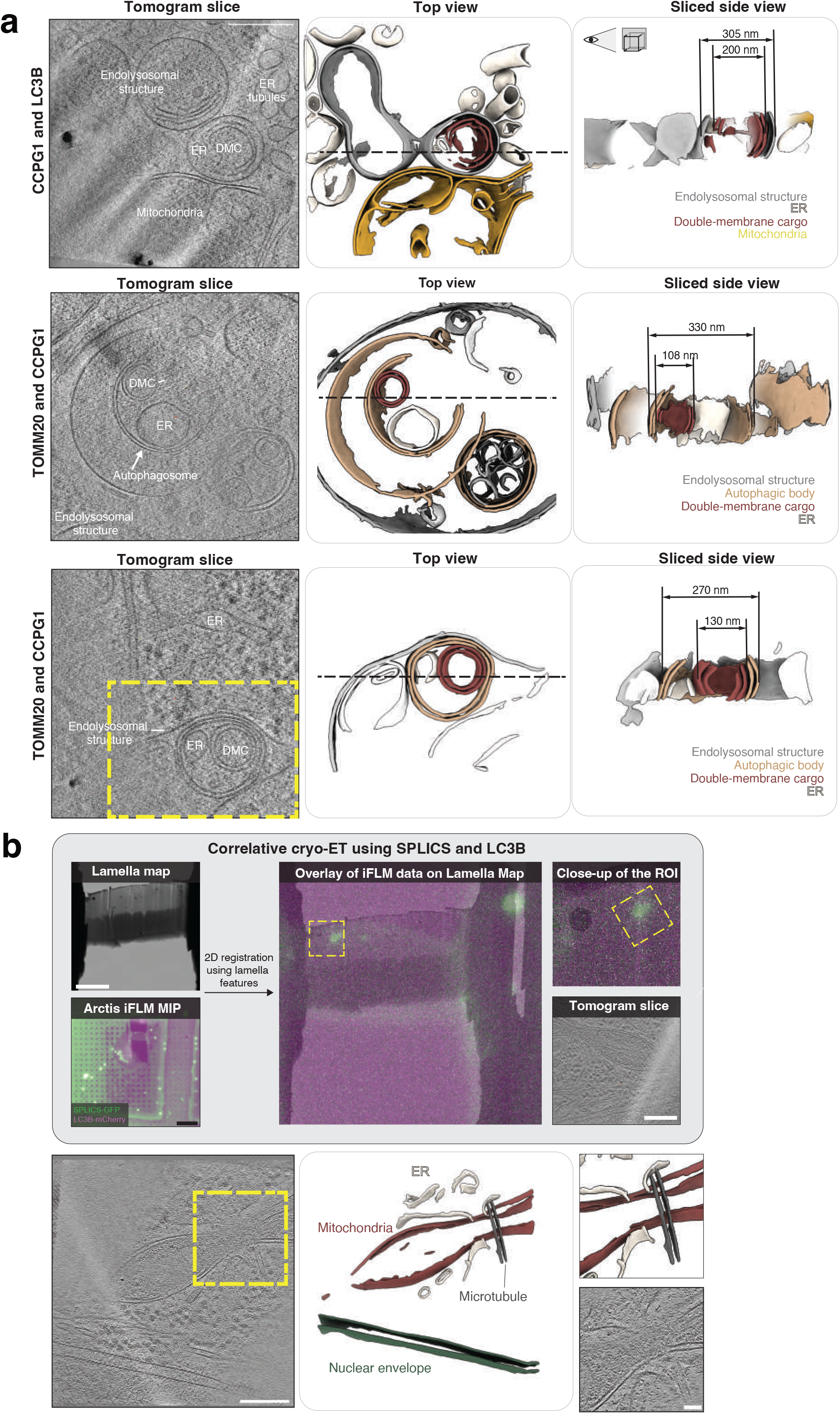
Correlated tomograms of HCT116 cells treated with EBSS and BafA1 display autophagic bodies with mixed content. **a)** (left) Tomographic slice of a tomogram of HCT116 cells after 3 hours of EBSS starvation and BafA1 treatment in a region positive for LC3B and CCPG1 (**top**) or in regions positive for TOMM20 and CCPG1 (**middle** and **bottom**). Double-membrane cargo (DMC) next to the ER is shown, some containing autophagosomes or autophagic bodies inside endo-lysosomal structures. (middle) A membrane segmentation of the tomograms is shown with membranes colored according to their organelle (in the lower panel only the bottom part of the tomogram within the yellow rectangle is segmented). (right) A sliced side view of the membrane segmentations with the indicated size of the autophagic bodies and the diameter of the DMC is shown for each example. **b)** Correlative cryo-ET images of HCT116 cells labelled with the SPLICS-ER-Mt system and LC3B. Images acquired for the correlation of SPLICS-ER-Mt (green) and mCherry-LC3B. A medium magnification stitched lamella map acquired at the TEM is shown above the final iFLM stack showing on lamella fluorescence. The middle image shows an overlay of the lamella map with fluorescence. On the right-hand side, a close-up image around the region of interest (ROI) is shown with the fluorescence overlay and finally a slice of the tomogram reconstructed from the tilt series at that position is shown below. Slice of a tomogram in a position positive for the SPLICS-ER-Mito marker. The region highlighted in yellow shows a constriction site in the mitochondria where the ER is present together with a microtubule.

**Extended data Figure 6.**
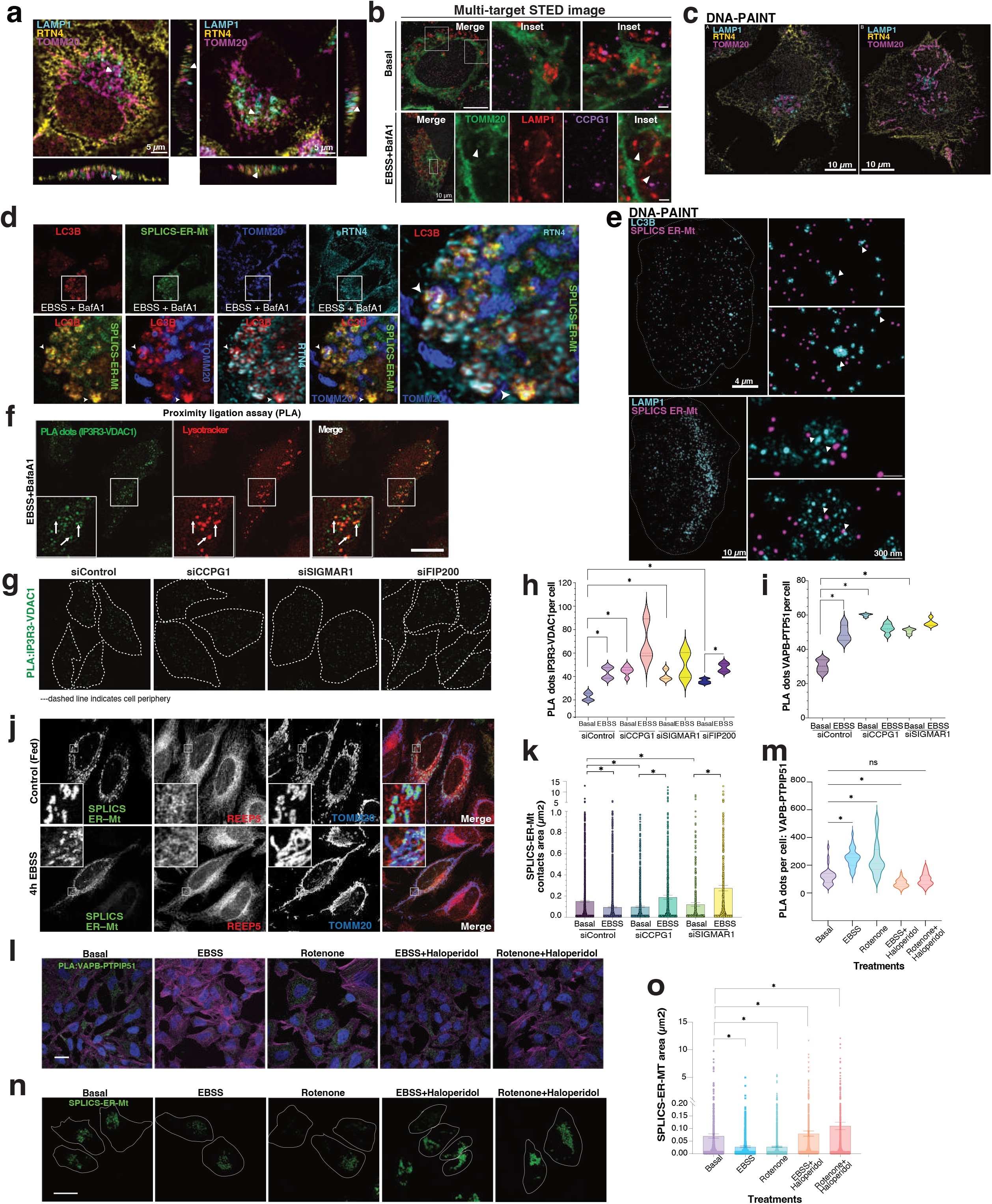
Single-molecule localization microscopy (SMLM) analysis and confocal microscopy analysis of contact sites degradation. **a)** Confocal images of RTN4+, TOMM20+ and LC3B+ or LAMP1+, arrowheads indicate colocalization of the signals. Orthogonal view of the projected image in yz and xz. Scale bar 5 µm. **b)** Multi-target STED image of U2OS cell expressing HaloTag TOMM20 treated with DMSO (Basal) and EBSS together with BafA1 co-stained for TOMM20 (green, HaloTag dead mutant coupled to ligand conjugated to SiR), LAMP1 (red, indirect immunofluorescence) and CCPG1 (magenta, LNA-PRISM). Arrows points to an exemplary ERMCSs inside a lysosome. Scale Bar, 10 µm, insets. **c)** Representative exchange-PAINT images of HeLa cells treated with EBSS+BafA1 (6 h) stained for LC3B or LAMP1 with RTN4 and TOMM20. Scale bar 10 µm. Zoomed in sections from A exemplifying LAMP1 positive vesicles or LC3B positive regions with TOMM20 or RTN4 signal either inside or in close proximity (>60 nm). **d)** Representative confocal image after LIGHTNING deconvolution of SPLICS-ER-Mt expressing cells colocalizing with LC3B (red), TOMM20 (blue) and RTN4 (cyan). Inset shows merge of all the channels, arrowheads show colocalization. Scale Bar, 2,5 µm. **e)** Representative exchange-PAINT images of HeLa cells expressing the fluorescence reporter SPLICS-ER-Mt contacts stained for LC3B (=3) (Top panel), LAMP1 (n=3) (Bottom panel) in EBSS and BafA1 treated cells (6h). Scale bar: 4, 10 µm and 300 nm. Arrowheads indicate localizations positive for ERMCSs in close proximity to LC3B signal or inside LAMP1 positive vesicles. **f)** Confocal images of Proximity ligation assay (PLA) for IP3R3 (ER) and VDAC1 (Mitochondria OMM), imaged alongside mCherryLAMP1. Arrows indicates PLA signal localizes within the LAMP1+ structures. **g)** Representative image analysis of the PLA dots IP3R3-VDAC1 per cells in siControl, siCCPG1, siSIGMAR1, and siFIP200 KD cells. The dashed line indicates cell periphery. **h)** Violin plot of PLA dots of IP3R3-VDAC1 coupling per cell in siControl, siCCPG1 and siSIGMAR1 cells treated with EBSS (6h). Data are mean ± s.e.m. from three independent experiments Basal n=984 for siControl, n=370 for siCCPG1 and n=334 for siSIGMAR1and EBSS n=441 for siControl, n=431 for siCCPG1 and n=186 for siSIGMAR1; Kruskal–Wallis/Dunn’s post-test. **i)** Violin plot of PLA dots of VAPB and PTPIP51 coupling per cell in siControl, siCCPG1 and siSIGMAR1 cells treated with EBSS (6h). Data are mean ± s.e.m. from three independent experiments Basal n=984 for siControl, n=370 for siCCPG1 and n=334 for siSIGMAR1and EBSS n=441 for siControl, n=431 for siCCPG1 and n=186 for siSIGMAR1; Kruskal–Wallis/Dunn’s post-test). **j)** Representative images of SPLICS-ER-Mt contact reporter localization with REEP5+ and TOMM20+ under basal and starvation conditions (EBSS for 6h). Inset shows the morphological changes on the contact sites. Scale bar, 10 µm. **k)** Violin plot of SPLICs ER-Mt contact sites density per cell. Data are mean ± s.e.m. from three independent experiments Basal siControl (n=12), siCCPG1 (n=15) and siSIGMAR1 (n=13) and EBSS siControl (n=12), siCCPG1 (n=13) and siSIGMAR1 (n=12); Kruskal–Wallis/Dunn’s post-test. **l)** Representative images of PLA VAPB-PTPIP51 changes upon EBSS, Rotenone (1 μM), in presence of the SIGMAR1 agonist (Haloperidol, 10uM) compared to Basal conditions. Scale Bar, 10 µm. **m)** Violin plot of PLA dots of VAPB and PTPIP51 coupling per cell in EBSS, Rotenone (1 μM), in presence of the SIGMAR1 agonist (Haloperidol, 10uM) compared to Basal conditions. Data are mean ± s.e.m. from three independent experiments Basal n=30; EBSS n=32; Rotenone n=31; EBSS+Haloperidol n=41; Rotenone+Haloperidol n=38). Kruskal–Wallis/Dunn’s post-test). **n)** Representative images of SPLICS-ER-MT remodeling by monitoring the area changes upon EBSS, Rotenone and inhibited by SIGMAR1 agonist (Haloperidol) for 4h compared to Basal conditions. Scale Bar, 10 µm. **o)** SPLICS-ER-MT remodeling by monitoring the area changes upon EBSS, Rotenone and inhibited by SIGMAR1 agonist (Haloperidol) compared to Basal conditions (Basal n=20; EBSS n=17; Rotenone n=18; EBSS+Haloperidol n=19 and Rotenone+Haloperidol n=22). Data represent the mean ± s.e.m. from three independent experiments; Kruskal–Wallis/Dunn’s post-test.

**Extended data Figure 7.**
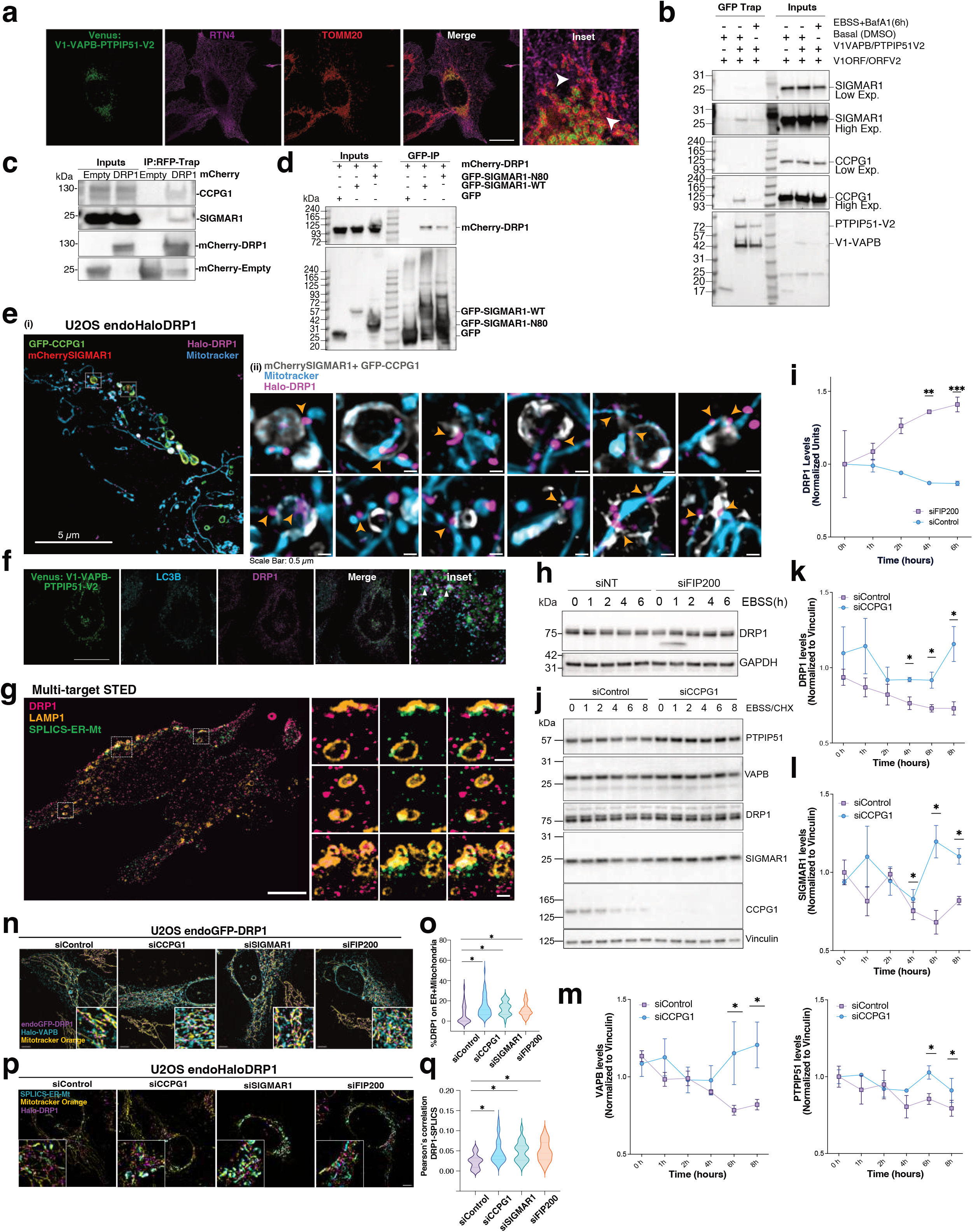
DRP1 associates to CCPG1-SIGMAR1 complex and the interactions depends on SIGMAR1 chaperone domain. **a)** Representative LIGHTNING-deconvolved images showing complemented Venus VAPB-PTPIP51 signal that localizes with ER (RTN4+) and mitochondria (TOMM20+) markers. Arrow heads show the triple co-localization of the Venus signal. Scale Bar, 10 µm. **b)** Representative western blot analysis of the Venus VAPB-PTPIP51 immunoprecipitation with GFP-trap beads showing the expression of VAPB and PTPIP51 and their interaction with SIGMAR1 and CCPG1. **c)** Co-immunoprecipitation assay of RFP-trap immuno-isolated of mCherry-DRP1interacting with endogenous SIGMAR1 and CCPG1. **d)** Co-immunoprecipitation assay and western blot analysis of GFP-trap immuno-isolated SIGMAR1 WT and N80 mutant (chaperone domain deletion) showed reduced interaction with mCherry-DRP1. **e)** SIM microscopy in U2OS cells showing endogenous Halo-DRP1 (magenta, labelled with JF646), transfected mCherry-SIGMAR1 (red), transfected GFP-CCPG1 (green), and mitochondria (cyan; labelled with BioTracker^TM^ 405 Blue mitochondria dye). Scale bar, 5 µm. **(i)** indicate ROI area used for 3D-rendering in **main** Fig. 4f and **(ii)** insets collected from different cells showing the CCPG1 and SIGMAR1 structures (grey) associated to DRP1 (magenta) and mitochondria (cyan). Scale bar, 0.5 µm. Zoomed images examples showing DRP1 association to CCPG1 and SIGMAR1 vesicles in close proximity to mitochondria at constriction point (early stage) or fission (later stage). **f)** Representative LIGHTNING-deconvolved images showing complemented Venus VAPB-PTPIP51 signal localizing with LC3B (cyan) and DRP1 (magenta). Arrow heads show the triple co-localization of the Venus signal. Scale Bar, 10 µm. **g)** Representative multi-target STED image of HeLa cells co-stained for DRP1, LAMP1 and SPLICS marker. The inset ROIs in the representative image are zoomed in on the right, with DRP1 and LAMP1; SPLICS and LAMP1 and all three targets together respectively. Scale bar: 10 µm and 1 µm. **h)** Western blot analysis of DRP1 protein levels in siControl and siFIP200 knockdown cells for 72 h under EBSS treatment for 0, 2, 4, 6 and 8 h. **i)** Quantification analysis of western blot **h** data represents the mean ± s.d. n=2 independent experiments. Two-way ANOVA, Bonferroni post-hoc test. **J)** Western blot analysis of DRP1 protein levels in siControl and siCCPG1 knockdown cells for 72 h under EBSS treatment for 0, 2, 4, 6 and 8 h. **k), l) and m)** Quantification analysis of western blot **j)** data represent the mean ± s.d. n=2 independent experiments. Two-way ANOVA, Bonferroni post-hoc test. **n)** SIM images from U2OS-endomEGFP-DRP1, with knockdown for CCPG1, SIGMAR1, FIP200 and control; showing the distribution of DRP1 (magenta) on mitochondria (yellow, labelled with MitoTracker Orange) and ER marker Halo-VAPB (cyan, labelled with JF646). Scale bar, 5 µm. **o)** Violin plots showing percentage of DRP1 associated with ER and mitochondria contact sites, from the SIM images of U2OS-endomEGFP-DRP1, with knockdown for CCPG1, SIGMAR1, FIP200 and control shown in panel N. (n=19, n= 14, n= 24 and n=20, respectively); Data represent the median (middle line) and the interquartile range (upper/lower lines as 75^th^ (Q3) and 25^th^ (Q1) percentile, respectively) from two independent set of experiments) represent the mean ± s.e.m. from two independent experiments; Kruskal–Wallis/Dunn’s post-test; *p<0.05 **p)** SIM images from U2OS-endoHalo-DRP1, with knockdown for CCPG1, SIGMAR1, FIP200 and control; showing the distribution of DRP1 (magenta) on SPLICS (long) and mitochondria. Scale bar, 5 5 µm. **q)** Plot showing Pearson’s correlation between DRP1 and SPLICS calculated from SIM images shown in panel C. (n=28, n=15, n=29 and n=28); Data represent the median (middle line) and the interquartile range (upper and lower lines as 75^th^ (Q3) and 25^th^ (Q1) percentile, respectively) from two independent set of experiments) represent the mean ± s.e.m. from two independent experiments; Kruskal–Wallis/Dunn’s post-test; *p<0.05.

**Extended Data Fig. 8.**
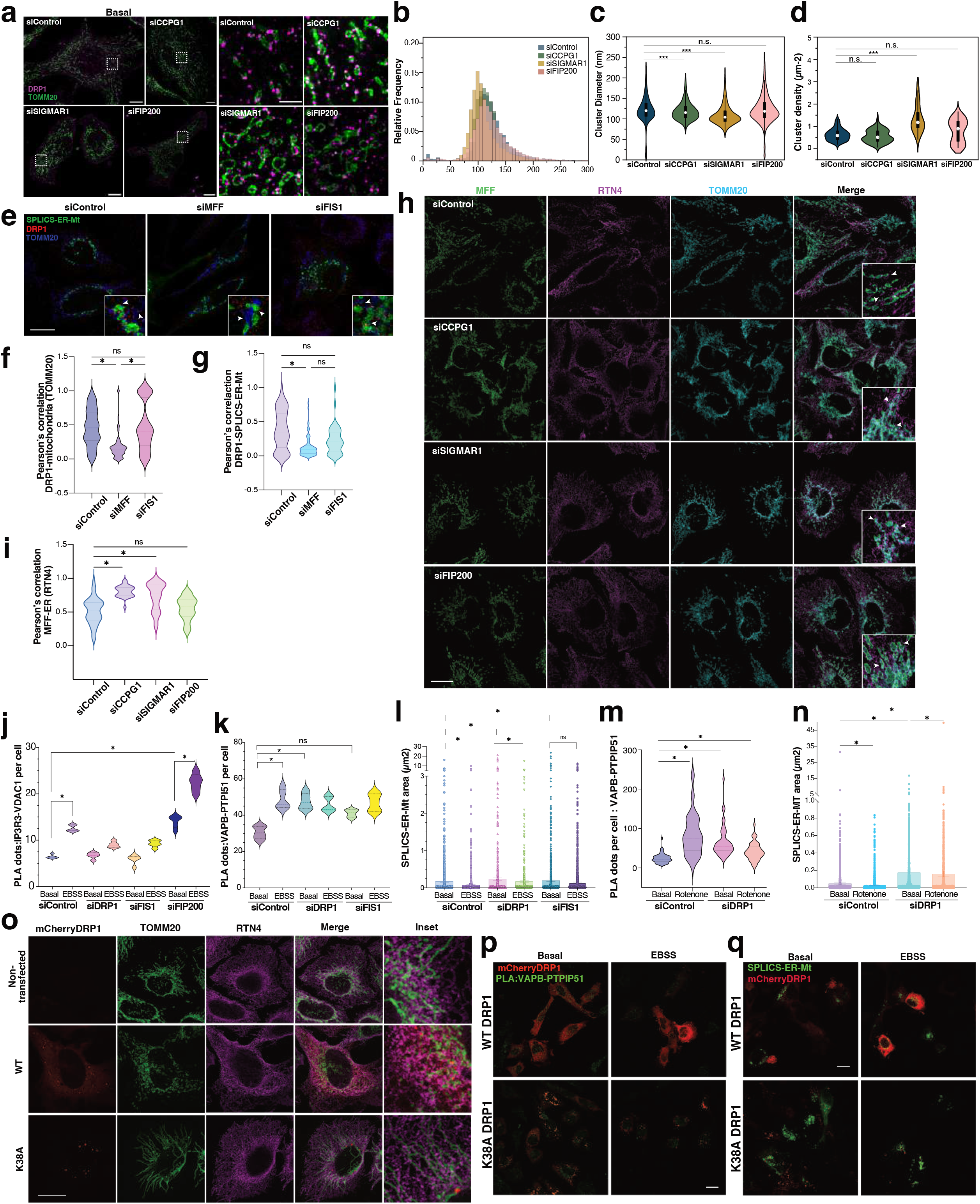
DRP1 and MFF localization at ERMCS are altered in CCPG1, SIGMAR1 and FIP200 deficient cells. **a)** Representative STED images of HeLa cells with control and knockdown of CCPG1, SIGMAR1 and FIP200 respectively co-stained for DRP1 (magenta) and TOMM20 (green) right panel Zoom-in of the ROIs shown in left panel showcasing the distribution of DRP1 and the shape of mitochondria across the knockdown conditions. Scale bar: 10 µm and 2 µm. **b)** Distribution of cluster diameter of DRP1 under various knockdown conditions. **c)** Violin plot showcasing DRP1 cluster diameter across different knockdown conditions. ***-p<0.0001, n.s. not significant. **d)** Cluster density (µm^−2^) of DRP1 across different ROIs in a cell under various knockdown conditions (siControl, n=9 images; siCCPG1, n=5 images; siSIGMAR1, n=4 images; siFIP200, n=5 images). **e)** Representative LIGHTNING-deconvolved images showing DRP1 localization in cells expressing SPLICS-ER-Mt (ERMCS marker) and stained mitochondria (TOMM20) in siMFF and siFIS1 knockdown cells. Arrowheads within the insets show the localization/recruitment of DRP1 to ERMCS or mitochondria. Scale bar: 10 µm. **f)** Violin plot of Pearson’s correlation analysis of DRP1 and TOMM20 in siControl compared to siMFF and siFIS1 knockdown cells (n=10, n=8 and n=12 cells respectively) represent the mean ± s.e.m. from two independent experiments; Kruskal–Wallis/Dunn’s post-test. **g)** Violin plot of Pearson’s correlation analysis of DRP1 and SPLICS-ER-Mt in siControl compared to siMFF and siFIS1 knockdown cells (n=10, n=8 and n=12 cells respectively) represent the mean ± s.e.m. from two independent experiments; Kruskal–Wallis/Dunn’s post-test. **h)** Representative LIGHTNING-deconvolved images showing MFF co-localization with ER (RTN4) and mitochondria (TOMM20) markers. Arrowheads within the insets show the localization of MFF with ER or mitochondria. Scale bar: 10 µm. **i)** Violin plot of Pearsons correlation analysis of MFF and RNT4 in siControl compared to siCCPG1, siSIGMAR1 and siFIP200 knockdown cells (n=8, n=9 and n=12 cells respectively). Data represent the mean ± s.e.m. from two independent experiments; Kruskal– Wallis/Dunn’s post-test. **j)** PLA dots analysis of IP3R3 and VDAC1 in siControl (basal n=979 and EBSS n=809), siDRP1 (basal n=910 and EBSS n=576) and siFIS1 (basal n=1420 and EBSS n= 294+399+316) knock down cells. Data represent the mean ± s.e.m. from three independent experiments; Kruskal–Wallis/Dunn’s post-test. **k)** PLA dots analysis of VAPB-PTPI51 in siControl (basal n=984 and EBSS n=441), siDRP1 (basal n=983 and EBSS n=847) and siFIS1 (basal n=983 and EBSS n=864) knock down cells. Data represent the mean ± s.e.m. from three independent experiments; Kruskal–Wallis/Dunn’s post-test. **l)** SPLICS-ER-MT remodeling by monitoring the area changes upon starvation in siControl (basal n=16 and EBSS n=12), siDRP1 (basal n=15 and EBSS n=12) and siFIS1 (basal n=11 and EBSS n=10) knockdown cells. Data represent the mean ± s.e.m. from two independent experiments; Kruskal–Wallis/Dunn’s post-test. **m)** Violin plot of PLA dots analysis of VAPB-PTPIP51 in cell siControl and siDRP1 knockdown cells in basal and rotenone induced stress. (siControl Basal n=44, Rotenone n=21 and siDRP1 Basal n=45, Rotenone n=34, respectively). Data represent the mean ± s.e.m. from two independent experiments; Kruskal–Wallis/Dunn’s post-test. **n)** SPLICS-ER-MT remodeling by monitoring the area changes upon rotenone in siControl and siDRP1 knockdown cells in basal and rotenone induced stress. (siControl Basal n=22, rotenone n=20 and siDRP1 Basala n=14, rotenone n=27). **o)** Representative images analysis of cells expressing WT DRP1 and the catalytic inactive mutant K38A DRP1 and the changes in mitochondria and ER interfaces. Scale Bar, 10 µm. **p)** Representative images of cells expressing mCherryDRP1 WT and K38A mutant and the changes in PLA dots VAPB-PTPIP51 in basal and starvation conditions (6h). Scale Bar, 10 µm. **q)** Representative images of cells expressing mCherryDRP1 WT and K38A mutant together with the SPLICS-ER-Mt fluorescence ERMCS reporter in basal and starvation conditions (6h). Scale Bar, 10 µm.

**Extended data Figure 9.**
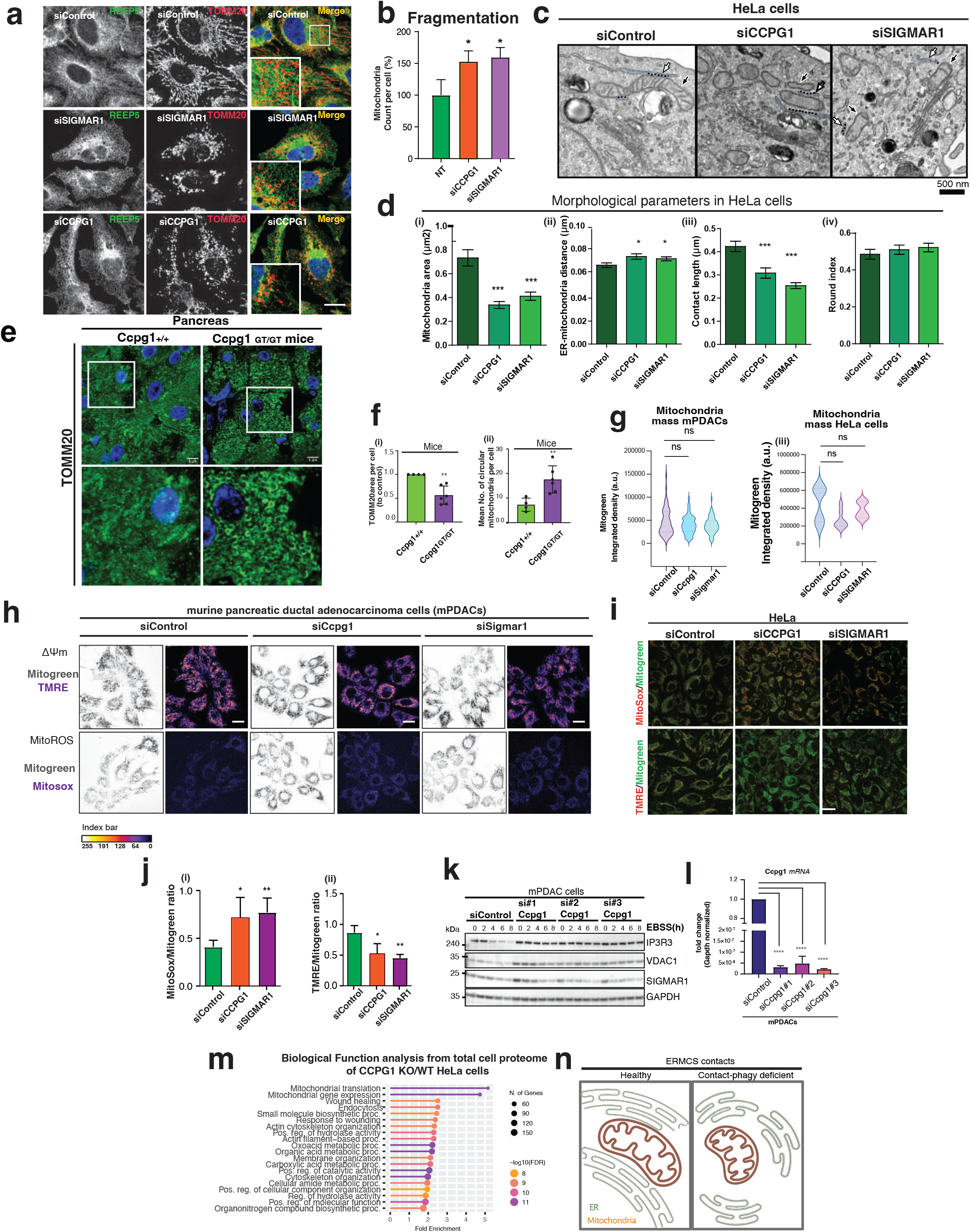
Importance of CCPG1-SIGMAR1 complex in Mitochondria Homeostasis. **a)** Confocal image analysis of ER (REEP5+) and Mitochondria (TOMM20+) distribution in siControl, siCCPG1, and siSIGMAR1 KD HeLa cells. **b)** Quantification analysis of mitochondria TOMM20^+^ structures for siControl (n=891), siSIGMAR1 (n=934) and siCCPG1 (n=943) number of cells. **c)** TEM images of siControl, siCCPG1 and siSIGMAR1 depicting ER and mitochondria contacts (dashed black line). ER (white arrows) is depicted in blue false color. Mitochondria are indicated by black arrows. **d)** Morphological mitochondria parameters were analyzed from TEM images (siControl, n=10; siCCPG1, n=10; siSIGMAR1, n=10); **i)** Mitochondria area (μm)^2^, **ii)** ER-mitochondria distance (μm), **iii)** contact length (μm) and **iv)** round index. **e)** Immunohistochemical analysis in the pancreas of WT and Ccpg1 ^GT/GT^ mice. Fixed tissue sections were stained with TOMM20. **f)** Quantitative analysis of TOM20 staining showing the relative area (top panel (i)) and the mean number of circular mitochondria per cell (bottom panel (ii)). Each panel point is the mean from different animal. **g)** Representative confocal images of mPDAC cells Control, siCcpg1 and siSigmar1 stained with the MitoSOX and TMRE dyes. The index bar indicates the fluorescence level. **h) and i)** Mitochondrial membrane potential and superoxide levels were measured in siControl, siCCPG1, and siSIGMAR1 KD HeLa cells (Mitosox: siControl n=972; siCCPG1 n=1139; siSIGMAR1 n=1360. TMRE: siControl n=1044; siCCPG1 n=1139; siSIGMAR1 n=1280). For this, we used MitoSox and TMRE, respectively, and quantified by analyzing the fluorescence intensity. **j)** Western blot analysis of total protein levels in siControl and siCcpg1 KD cells. ERMCS protein levels were detected under time course with EBSS treatment for 0, 2, 4, 6 and 8h. **k)** Relative mRNA expression levels of CCPG1 normalized to GAPDH in mPDACs siControl, siCcpg1 #1, siCcpg1 #2 and Ccpg1 #3 cells. Data are presented as the mean with error bars indicating the s.d. Statistical significance of differences between experimental groups was assessed with ANOVA/Bonferroni post-test. Differences with p<0.0001 are annotated as ****; n = 3. **l)** Biological function pathway analysis from total cell proteome of CCPG1 KO/WT Hela Cells. Top hits were analyzed using the Shiny GO platform and sorted by FDR (log_2_enrichment > 0.25 and –log_10_ p > 1.3, one-tailed unpaired Student’s test). *n* = 3 independent experiments. **m)** Model indicates that the absence of CCPG1 and SIGMAR1 impairs the Contact-phagy leading to a dysfunctional mitochondrion evidenced by spherical and swollen morphology accompanied of smaller ERMCSs contacts and increased mitochondria distance to ER.

**Extended data Figure 10.**
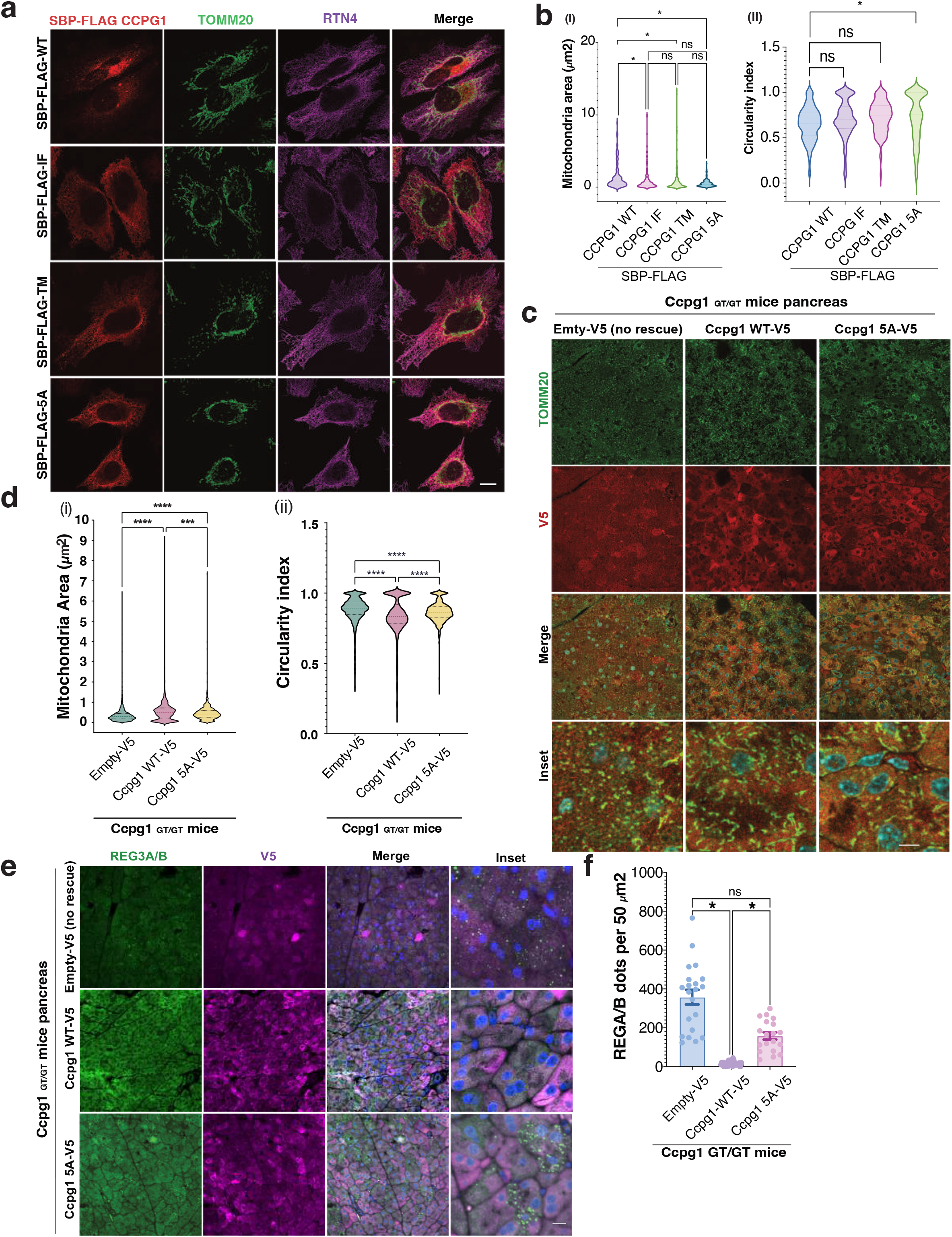
Re-expression of CCPG1 WT rescue the mitochondrial and ERMCSs abnormalities while a CCPG1-deficient SIGMAR1 binder failed. **a)** Representative immunofluorescence analysis of CCPG1-deficient SIGMAR1 binding mutants (WT, interfacial (IF), transmembrane (TM) and IF+TM(5A)), mitochondria (TOMM20) and ER (RNT4). **b)** Morphological analysis of mitochondria area (i) and circularity (ii) in cells expressing the CCPG1-deficient SIGMAR1 binding mutants (WT n=8, IF n=7, TM n=5 and 5A n= 5 cells per group). **c)** Representative immunofluorescence images of TOMM20+, V5 and DAPI from Ccpg1^GT/GT^ mice pancreas re-constituted with Empty-V5, Ccpg1 WT-V5 and Ccpg1^5A^-V5 mutant. (n=3 animals per group **d)** Bar plot results of mitochondria parameter from immunofluorescence analysis form pancreas sections of the re-expressed Ccpg1 WT and the Ccpg1 5A-SIGMAR1 deficient binding mutant (Ccpg1 ^GT/GT^ re-expressing the Luciferase empty vector n=3, reconstituted with Ccpg1 WT n=3 and Ccpg1 5A n=3 mice per group). **e)** Representative images of REG3A/B+, V5, DAPI staining from Ccpg1^GT/GT^ mice pancreas re-constituted with Empty-V5, Ccpg1WT-V5 and Ccpg1^5A^-V5 mutant. **f)** Quantification analysis of REG3A/B in Ccpg1^GT/GT^ mice pancreas re-constituted with Empty-V5 Ccpg1WT-V5 and Ccpg1^5A^-V5 mutant. Data represent the mean ± s.e.m. from n=3 animals per group, Kruskal– Wallis/Dunn’s post-test.

## Supplementary information

**Table 1. Per-residue CCPG1–SIGMAR1 contact summary from coarse-grained simulations of full-length CCPG1 (residues 1–250) with SIGMAR1 in a POPC membrane (n = 3)**. Contacts were retained using a contact-probability threshold (>0.5) and a minimum of 3 partner residues. Cytosolic CCPG1 residues are subject to two additional, independent criteria: they retain their full contact list only if they form at least 15 contacts, whereas residues below this density are retained exclusively through salt bridges (ionic interactions), requiring a minimum of 2. Columns list SIGMAR1 partners, structural region, summed contact probability (± propagated error), and interaction-type breakdown.

**Table 2. Per-residue CCPG1–SIGMAR1 contact summary from all-atom simulations of the truncated CCPG1 construct (residues 197–250), pooling POPC and ER-membrane replicas (n = 4)**. Contacts were retained using a contact-probability threshold (>0.05) and a minimum of 1 partner residue. Cytosolic CCPG1 residues are subject to two additional, independent criteria: they retain their full contact list only if they form at least 15 contacts, whereas residues below this density are retained exclusively through salt bridges (ionic interactions), requiring a minimum of 2. Columns list SIGMAR1 partners, structural region, summed contact probability (± propagated error), and interaction-type breakdown

